# SciPhy: A Bayesian phylogenetic framework using sequential genetic lineage tracing data

**DOI:** 10.1101/2024.10.01.615771

**Authors:** Sophie Seidel, Antoine Zwaans, Samuel Regalado, Junhong Choi, Jay Shendure, Tanja Stadler

**Author notes:** Equal contribution (Sophie Seidel, Antoine Zwaans).

## Abstract

CRISPR-based lineage tracing offers a promising avenue to decipher single cell lineage trees, especially in organisms that are challenging for microscopy. A recent advancement in this domain is lineage tracing based on sequential genome editing, which not only records genetic edits but also the order in which they occur. To capitalize on this enriched data, we introduce SciPhy, a simulation and inference tool integrated within the BEAST 2 framework. SciPhy utilizes a Bayesian phylogenetic approach to estimate time-scaled phylogenies and cell population parameters. After validating SciPhy using simulations, we apply it to two lineage tracing datasets for which we estimate time-scaled trees together with cell proliferation rates. Using simulated and real lineage tracing data obtained from a monoclonal culture of HEK293T cells, we compare SciPhy to other lineage reconstruction methods, and find that SciPhy consistently constructs distinct, and more accurate lineage trees. In particular, for HEK293T cells, SciPhy trees stand out for their later estimated cell division times. In addition, SciPhy reports uncertainty as well as proliferation rates, neither of which are available within a UPGMA analysis. Our second example applies SciPhy to the study of murine gastruloids, and showcases the use of complex models of time-varying population growth to capture realistic aspects of this in-vitro model of early mammalian development. Together, these examples showcase the application of advanced phylogenetic and phylodynamic tools to explore and quantify cell lineage trees, laying the groundwork for enhanced and confident analyses to decode the complexities of biological development in multicellular organisms. SciPhy’s codebase is publicly available at https://github.com/azwaans/SciPhy.

## 1 INTRODUCTION

The interplay between cell division, cell death and differentiation is at the core of the development of all multicellular organisms. Tracing cell populations and their lineage relationships via DNA-based molecular recording has recently emerged as a powerful approach to study this complex process (VanHorn and Morris (2021)). Among the technologies developed for lineage recording, CRISPR-Cas9-based approaches stand out for generating stochastic, heritable indels in genetic barcodes during development and proliferation (McKenna et al. (2016); McKenna and Gagnon (2019); Chan et al. (2019); He et al. (2022); Bowling et al. (2020); Alemany et al. (2018); Spanjaard et al. (2018)). However, early approaches relying on *unordered indels* combined with a short recording duration limit the reconstruction of highly resolved cell ‘lineage trees’ (see Glossary, Tab. 1) (Loveless et al. (2021); Choi et al. (2022)).

**Table 1.** Glossary. Translation of phylogenetic terminology to cell biology terminology.

| Phylogenetic Terms | Cell Biology Terms |
| --- | --- |
| Tree topology | Lineage tree |
| Clock rate | Edit insertion rate by prime editor (e.g. per hour) |
| Bifurcation, branching point | Cell division event |
| Branching time | Cell division time |
| Time-scaled tree | Lineage tree where branches are scaled in real time |
| Monophyletic | Monoclonal |

Recent work (Loveless et al. (2021), Choi et al. (2022)) addresses some of these limitations by introducing prime-editing based recording systems that enable lineage tracing over longer time periods. These methods rely on *ordered insertions* of nucleotide sequences at target sites mediated by the prime editor and prime editing guide RNA (pegRNA) (Anzalone et al. (2019)). Each insertion at a target site deactivates the current site and activates the next for subsequent editing. This ensures irreversible edit accumulation and records the order of edits within the target. These new lineage tracing technologies have the potential to enable the reconstruction of accurate, highly resolved single-cell lineage trees.

To date, lineage tree reconstruction using such data Choi et al. (2022); Loveless et al. (2021) has relied on UPGMA (Unweighted Pair Group Method with Arithmetic Mean, Sokal et al. (1963)) in tandem with custom distance metrics. This approach is agnostic to specific properties of the editing process, such as variable propensities of introducing particular inserts and variable edit insertion rates (see Glossary, Tab. 1). UPGMA, being a distance-based clustering method, only leverages pairwise distances between cells, ignoring all higher order information (Felsenstein (2004)). In contrast, Bayesian phylogenetic inference methods for unordered edit lineage tracing data (Seidel and Stadler (2022)) enable the estimation of lineage trees with cell division times (see Glossary, Tab. 1) by mechanistically modelling barcode generation. This approach was shown to provide accurate lineage reconstruction, attributed to explicitly incorporating *a priori* knowledge of the editing process and the experimental conditions under which traced cell populations were studied.

In this article, we derive a mechanistic model of sequential insertion-based editing (ordered inserts) for lineage tracing data. This model forms the basis of our new inference framework, ‘SciPhy’ **S**equential **C**as-9 **I**nsertion based **Phy**logenetics, implemented in the Bayesian inference software BEAST 2 (Bouckaert et al. (2014)). BEAST 2 uses a Markov Chain Monte Carlo framework to jointly estimate time-scaled lineage trees (see Glossary, Tab. 1) and editing dynamics. This integration further makes phylodynamic analyses of single-cell population dynamics directly accessible to the community.

We showcase the versatility of our validated framework with a comprehensive analysis of two lineage tracing datasets: the first derived from a monoclonal (see Glossary, Tab. 1) HEK293T culture (Choi et al. (2022)) and the second from the in vitro differentiation of a single mouse embryonic stem cell (ESC) into a multicellular mouse gastruloid, a protocol that models aspects of early mammalian development. Our analyses reveal that complex editing dynamics govern the recording process in both datasets and estimate the growth rates of the cell populations over time. We observe significant differences between our lineage tree estimates and those obtained using UPGMA, underscoring the impact of the reconstruction method on the inferred cellular relationships and growth dynamics.

## 2 RESULTS

We introduced a mechanistic editing model and likelihood calculation for sequential lineage tracing data as the core component of SciPhy (Fig. 1). This model assumes that stably integrated barcodes accumulate edits sequentially and irreversibly at a constant Cas9 nicking rate. Subsequent pegRNA-mediated insertions occur with varying probabilities. We derived a phylogenetic likelihood to calculate the probability of observing a barcode alignment given a phylogeny and SciPhy parameters. Details of the model, its parameterization, and Bayesian inference implementation in BEAST 2 are in Methods and Materials.

**Figure 1.**
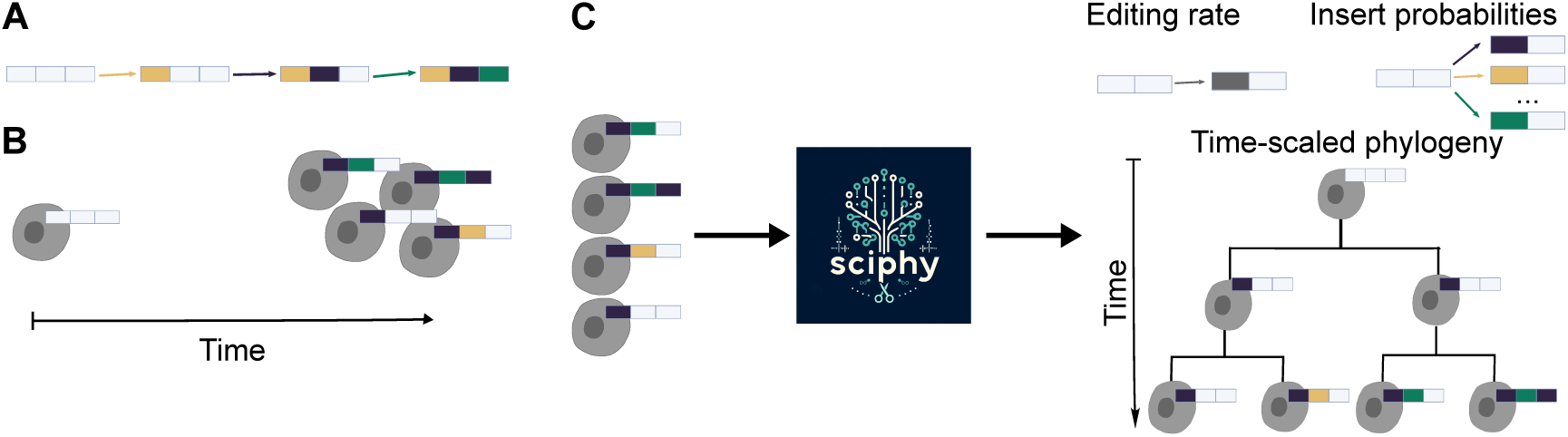
SciPhy is a framework for phylogenetic and phylodynamic analysis of sequential edit lineage tracing data. It implements a mechanistic model of sequential editing based recording systems (**A**), which enables their use for Bayesian phylodynamic inference in BEAST 2. These alignments are obtained following typical pipelines of genetic lineage tracing experiments: cells that harbour copies of the recording barcode, or ‘DNA Tape’, accumulate insertions during growth (**B**). At the end of these experiments, single-cell sequencing generates barcode alignments (**C**), from which SciPhy can infer parameters of the recording system: the rate and probability at which different insertions are acquired.

### 2.1 In-silico validation

We validated SciPhy using well-calibrated simulations Dawid (1982) to ensure that it draws samples from the correct posterior distribution. This process involved simulating trees and edited barcodes for each cell, and using them as input data to estimate back the lineage tree as well as the editing parameters under the SciPhy model. Editing parameters were drawn from distributions representing realistic editing dynamics. The same distributions were also used as priors in the inference (see Methods for details). Correct tree inference was assessed through three key summary statistics of inferred lineage trees, namely, the tree height, tree length, and tree balance, covering the properties of branch lengths and tree shape.

We calculated the coverage for each editing model parameter and tree summary statistic, defined as the fraction of datasets for which SciPhy inferred a 95% highest posterior density (HPD) interval containing the true (simulation) parameter. The coverages for all parameters fall within the expected range (Mendes et al. (2024), Supplementary table 6), demonstrating that our method is correctly implemented.

To evaluate the inference power across a range of parameter values, we plotted the simulated (true) parameters against the estimated parameters (Fig. 2 A-E). The correlations for the editing rate and the insertion probabilities (with a Pearson’s R value of 0.99 and 0.98, respectively, see Supplementary table 6 for 95% confidence intervals) confirm that SciPhy extracts meaningful signal from the data. Likewise, a high correlation for all three tree summary statistics is achieved (for tree height, tree length and tree balance, Pearson’s R of 0.90, 0.99 and 0.99 respectively), showing that signal for these features is also derived from the data.

**Figure 2.**
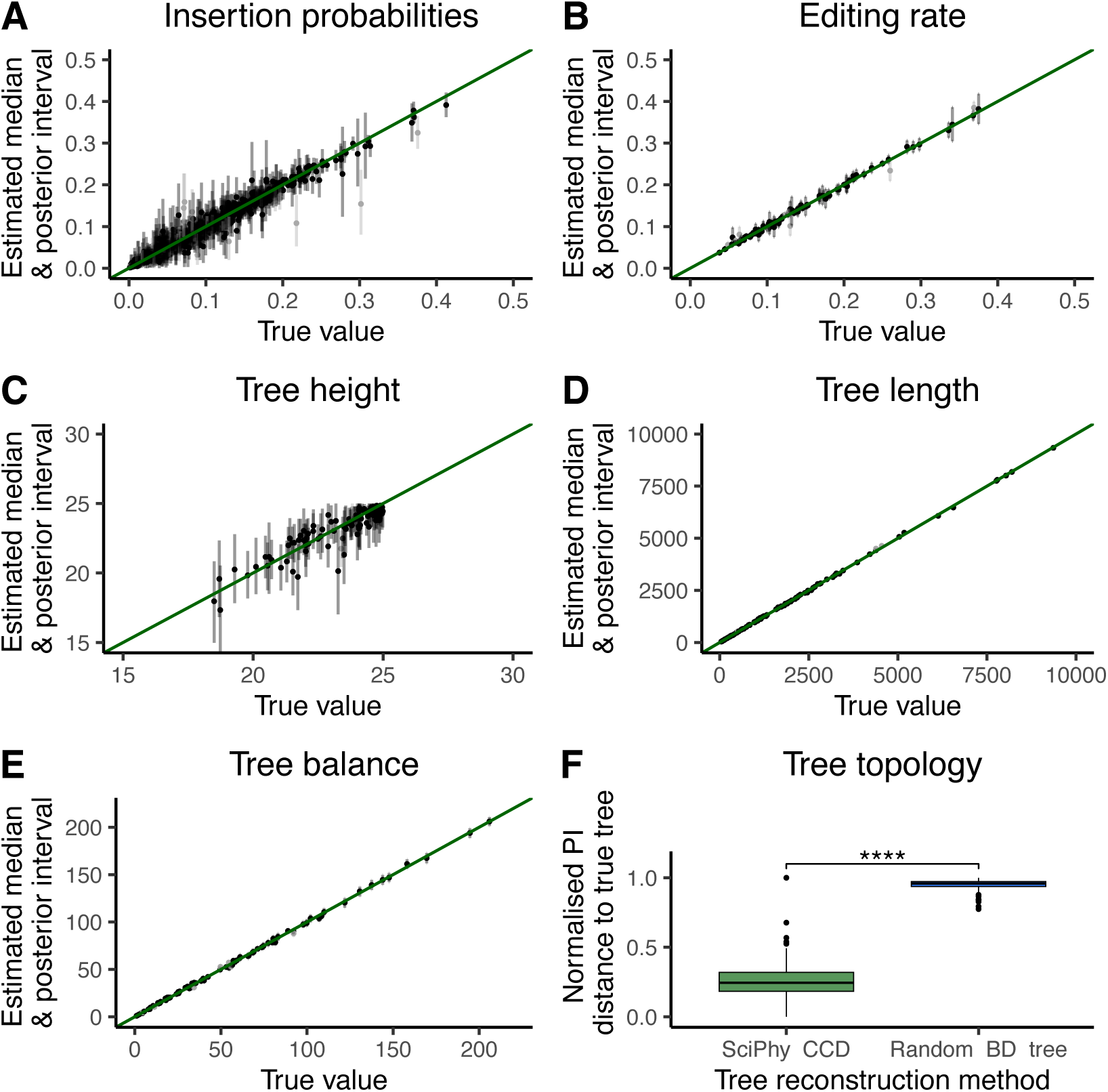
Validation results. We report the results of the in-silico validation for the editing rate (**A**), the insertion probabilities (**B**), tree height (**C**), tree length (**D**), tree balance (**E**) calculated using the B1 index, and tree topology (**F**) based on 100 simulations. In all panels **A-E**, we plot the estimated posterior medians against the true values. The grey intervals denote the 95% highest posterior density interval around the median. Estimated median posterior are plotted as black dots when the 95% highest posterior density interval contains the true values and as grey dots otherwise. The green line represents the performance of an ideal estimator. Panel **F** shows the topological distance (measured using the Phylogenetic Information metric) to the ground truth simulated tree with trees inferred with SciPhy (SciPhy CCD) or obtained by simulation under a simple birth-death model (Random BD tree), and annotated with the significance of a paired t-test between both methods.

We also assess SciPhy’s ability to extract signal for lineage relationships from the data by comparing topological distances (using the Phylogenetic Information (PI) distance, Smith (2020)) between random trees obtained by simulations under a simple birth-death model as well as SciPhy’s tree estimates (summarised as point estimates with a conditional clade distribution algorithm (CCD), Berling et al. (2025)) and the true simulated topologies. SciPhy consistently and significantly recovers topologies closer to the ground truth, thus confirming its ability to learn lineage relationships from this input data (Fig. 2 F).

Finally, we evaluate SciPhy’s robustness to two major sources of missing data found in lineage tracing datasets, namely, transgene silencing and sequencing dropout. To do so, we simulate alignments under varying levels of heritable tape loss (modeling transgene silencing) and loss upon sampling (modeling sequencing dropout), which both result in incomplete recovery of tape sequences per cell (see Materials and Methods for a description of this simulation process). Trees reconstructed using incomplete barcode alignments are farther from the ground truth tree than those that use complete ones as input (see Appendix fig. 23, top and 24), considering both the tree topology alone (using the PI distance) and branch length estimation (using the weighted Robinson Foulds (wRF) distance). We find that the PI distance for tree topology is particularly sensitive to tape loss and the resulting alignment sparsity (see Appendix fig. 23, bottom left and 24, left), while the wRF metric considering both topology and branch lengths results in a weaker increase in distance as tape loss probability increases.

### 2.2 Phylodynamic analysis of monoclonal expansion

We analyze a dataset where DNA Typewriter was used to trace lineage in a 25-day monoclonal expansion of HEK293T cells (Fig. 3, A). Data filtering was performed as previously (Choi et al. (2022), see Methods for details). Using SciPhy, we conducted a phylodynamic analysis to estimate the lineage tree, editing dynamics of the integrated barcodes, or ‘DNA Tapes’, and the growth dynamics of the sampled cell population (see Methods for details).

**Figure 3.**
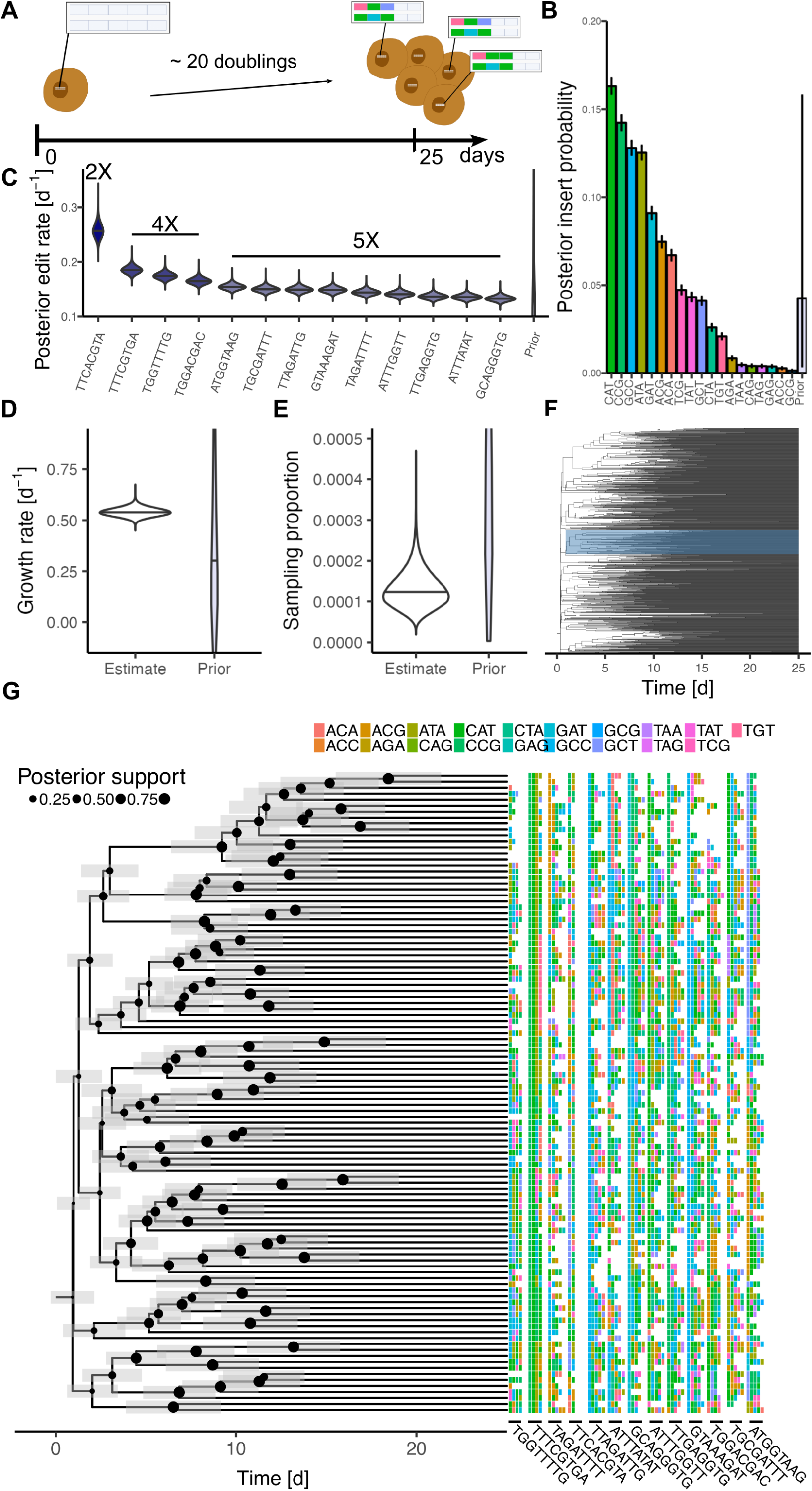
SciPhy captures complex editing dynamics and infers developmental parameters. (**A**) We apply SciPhy to a dataset from monoclonal expansion of HEK293T cells harboring the DNA Typewriter recording system. We report SciPhy’s parameter estimates as the posterior distributions of the editing model parameters, namely the insertion probabilities (**B**) and the editing rates per day per tape (**C**). The error bars on the insertion probabilities indicate the 95% highest posterior density interval. Editing rates for truncated tapes are colored with darker shades and annotated with the corresponding number of sites (respectively 2, 4, and 5 sites). We report estimates of the cell population dynamics under the birth-death-sampling model, namely the growth rate (**D**) and the sampling proportion (**E**). In (**F**) we plot the time-scaled Conditional Clade Distribution (CCD) summary tree for 1000 cells and highlight a clade that is shown enlarged in (**G**).We show a sub-clade of the CCD tree and highlight the posterior support in the node placement by the node thickness and the uncertainty, i.e. 95% HPD intervals, in the cell division time as grey shadings. We plot the alignment facing the tree tips.

We estimated SciPhy’s editing model parameters for this dataset, namely an editing rate for each DNA Tape and the overall insertion probabilities. Our estimates show substantial variation in the insert probabilities for different inserts (Fig. 3, B). Notably, the ‘CAT’ insert had the highest probability (16%), whereas ‘GCG’ had the lowest probability (≤ 1%). Although this variation, potentially due to differential pegRNA expression or hexamer efficiencies, was documented in the initial publication (Choi et al. (2022)), the UPGMA tree building method used there did not take it into account. Under the SciPhy model, we infer that when an insert is introduced, there is approximately a 50% chance that it will be one of the four most prevalent inserts (CAT, CCG, GCC, ATA). Consequently, these particular edits have a high chance of being introduced concurrently in otherwise independent lineages. Unlike UPGMA, SciPhy can account for this process by modelling these insert probabilities, potentially avoiding the reconstruction of incorrect lineage relationships.

We further estimate an editing rate, or clock rate, for each DNA Tape, which denotes the rate at which inserts are introduced at target sites within that tape (Fig. 3, C). Most DNA Tapes show clock rates of around 0.15 d^−1^, corresponding to an average of 4 edits per tape over the entire experiment, a value that aligns with our dataset observations (see Appendix Fig. 5).

Interestingly, all four truncated DNA Tapes, i.e. tapes with less than 5 target sites (see Methods for details), showed elevated clock rates (Fig. 3, C, first 4 tapes). Among these, the most truncated tape, ‘TTCACGTA’ —shortened to just two sites— had the highest rate. The observed pattern of editing rate ordering by tape length is unexpected under a null-model where all tapes have similar editing rates independent of tape length. We therefore asked whether early edits occur at a different rate from late edits. We compiled a dataset in which all tapes were shortened to length two and compared the estimated editing rates with those of the original data encompassing all sites (Appendix Fig. 6). Interestingly, the 2x tape ‘TTCACGTA’ is edited the fastest in both analyses, indicating that its elevated editing rate is not due to the first two edits occurring faster across tapes of any length. Given that we only have a single instance of the 2x tape, it is difficult to draw definite conclusions, but one possible explanation is that the 2x tape has been integrated into a genomic location that facilitates a high editing rate.

We then compared the 4x and the 5x tapes. In the analysis using all sites, the three highest editing rates among the 4x and 5x tapes were all 4x tapes. This result is surprising and statistically significant under the assumption of equal rate (p(k=3)=0.005, see Methods for test details). However, in the analysis using only 2 sites, this order was lost, with only one 4x tape among the three tapes with fastest editing rates. This result was not statistically significant under the assumption of equal rates (p(k=1)=0.5). Therefore, given the editing rate estimates under SciPhy, the difference in editing rate between 4x tapes and 5x tapes may be due to a difference in editing speed across the last 2-3 sites, for instance, slower editing at the 5th site of the 5x tapes. To further explore this scenario, we simulated data in which the 5th site was edited at 20% of the original rate. In this scenario, but not under equal edit rates, we observed two 4x tapes being among the 3 tapes with the highest inferred editing rates (see Appendix Fig 7), supporting our earlier statistical analysis. However, as our HEK293T analysis is based on a single experimental replicate, additional replicates would be required to draw definitive conclusions on the relative edit rates.

Along with estimating the editing model, we studied the growth dynamics of the HEK293T cell population by fitting a birth-death-sampling model to the set of sequences. These sequences are a subsample of 1000 cells from a population reported to consist of approximately 1.2 ∗ 10^6^ cells Choi et al. (2022). To incorporate this knowledge, we initially fixed the sampling proportion to 8 × 10^−4^, assuming constant population growth throughout the experiment. This model estimated a cell population growth rate between 0.42 and 0.47 days^−1^ (95% highest posterior density (HPD) interval, see Appendix Fig. 16) “Sciphy + Fixed sampling”), corresponding to doubling times of 35 to 40 hours. This growth rate in a model of exponential population growth implies a population of ≈ 10^5^ ± 10^5^ (mean ±*σ*) cells by the experiment’s end, which is an order of magnitude smaller than the reported 1.2 ∗ 10^6^ cells. After confirming that this growth rate estimate was driven by the data in tape alignments (see Appendix Sampling from prior), we hypothesized that this disagreement may stem from model misspecification. Therefore, we relaxed two of the main assumptions made in our model separately: first, the assumption of constant growth over the experimental period and, second, the assumption that we have accurate knowledge of the true sampling proportion.

First, we ran the same analysis with piecewise constant birth and death rates allowed to vary every two days over the timeline of the experiment. The estimated growth rates imply a pattern where the median growth rate is highest during the first 2 days of the experiment, with a median growth rate of 1.69 d^−1^ or doubling time of 9.6 hours (95% HPD of growth rate [0.57,2.76] d^−1^), and declines for 5 days after which it stabilizes around 0.3 d^−1^ until the end of the experiment (see Appendix Fig. 15). Under this set of rates, the cell population grows to an expected size of ≈ 9 ∗ 10^5^ cells, closer to the observed 1.2 ∗ 10^6^ cells. However, the fast doubling time in the first days is unrealistic for HEK293T cell growth from a single cell. It should further be noted that a constant growth rate over the experimental period is not excluded under these results.

Second, we relaxed our assumption on accurate knowledge of the sampling proportion, by jointly estimating it with the growth rate. This analysis inferred a growth rate between 0.49 and 0.58 days^−1^, or a doubling time that ranges between 28 and 33 hours (95% HPD interval, Fig.3, D). This doubling time is consistent with documented HEK293T doubling times Biodata (2024), and leads to an average population size between 2.5 ∗ 10^5^ and 2.3 ∗ 10^6^ (95% HPD) after 25 days, which agrees with the reported 1.2 ∗ 10^6^ cells.

Interestingly, the inferred sampling proportion under the last model is approximately four to 16 times lower than initially assumed (95% HPD interval: [5.2 × 10^−5^, 2.2 × 10^−4^], Fig. 3, E). Under the birth-death model parameterized for this analysis, a lower-than-expected sampling proportion may stem from measurement errors in estimating the total cell population size —potentially caused by cell clumping during counting— or from non-uniform sampling, such as biases in cell selection, sequencing, or filtering. Given that the reported population size of 1.2 × 10^6^ cells was measured using both a haemocytometer and a cell counter, which should provide reliable counts, it is more plausible that biases in the sampling process led to this reduced estimate. In such cases, the inferred sampling proportion likely reflects an ‘effective’ sampling proportion, accounting for these underlying biases rather than the true biological sampling proportion. Finally, we performed sensitivity analyses using alternative prior assumptions on the editing model parameters and found that our results remained robust across these settings (Appendix Fig. 21).

Taken together, the three analyses presented here suggest that a model of constant growth adequately describes the population dynamics under study.

### 2.3 Comparison to state-of-the-art

We compare SciPhy with existing methods for phylogenetic tree topology reconstruction on simulated and real data in Figure 4. For the comparison on simulated data, we reuse a subset of 9 datasets from the validation study (see Methods for details). We contrast SciPhy with current approaches that used order-aware UPGMA clustering. We also compared against methods that do not use the ordering information from the DNA Typewriter data, namely TiDeTree and UPGMA clustering based on an order unaware distance matrix. As a baseline, we included random trees simulated under a birth–death model with the same number of tips.

**Figure 4.**
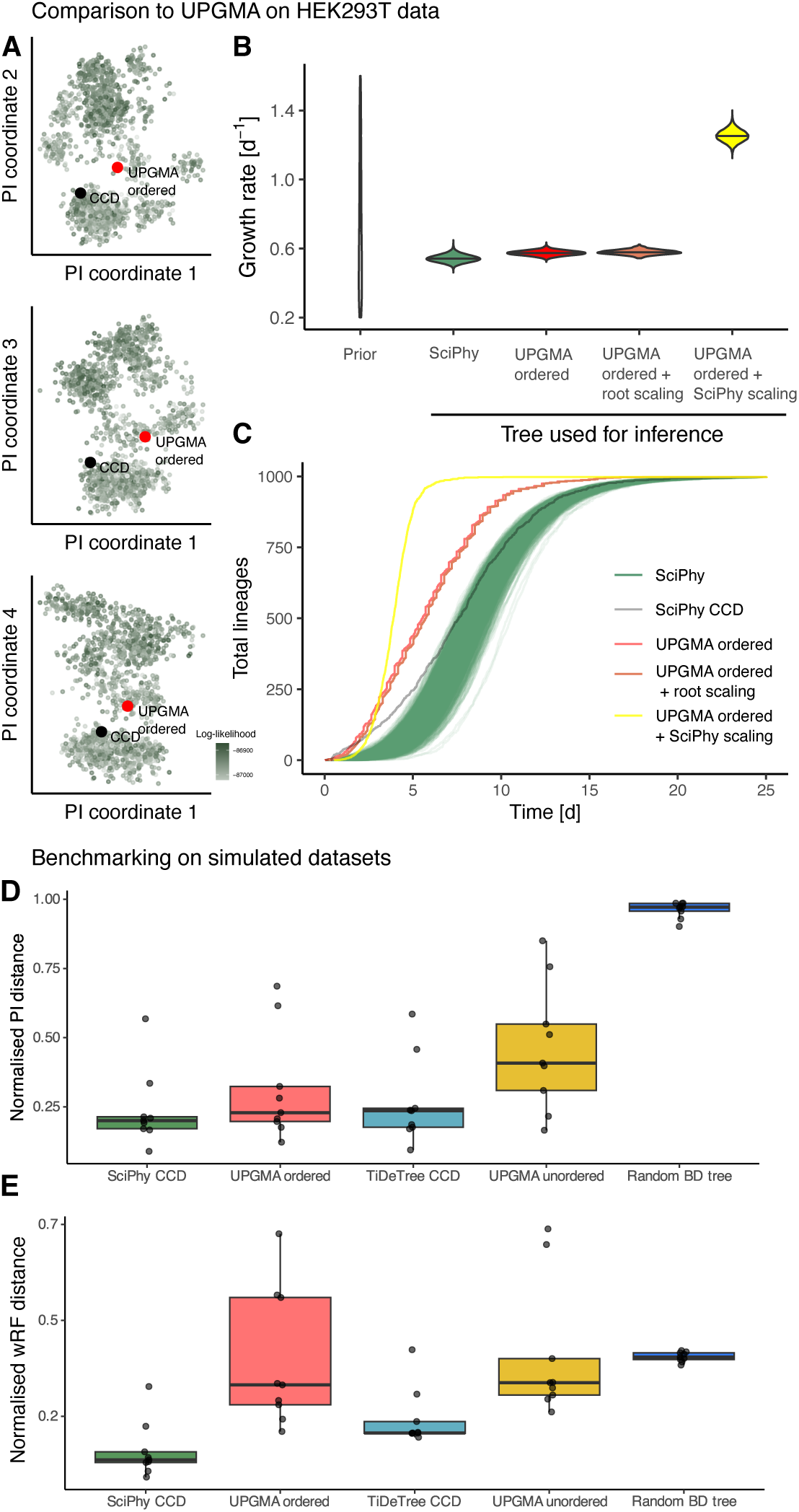
SciPhy produces distinct and more accurate estimates on real and simulated data, respectively. In panels (**A-C**) we show inference outputs obtained using UPGMA and SciPhy using the HEK293T dataset as input. In (**A**), we compare the SciPhy posterior tree set (green), including a summary tree, the CCD tree (black), to the tree reconstructed with the order-aware UPGMA method (red). We do so by calculating the Phylogenetic Information distance that captures topological distance between all pairs of trees. We then use multidimensional scaling (principal coordinates analysis) to project these distances into 12 dimensions. Here we show the results of the first coordinate against coordinates 2 to 4 from top to bottom. The remaining pairwise coordinate are shown in Appendix Fig. 10. In (**B**) we compare growth rates jointly estimated with the SciPhy posterior tree set inferred under the constant birth-death-sampling model (green, and the summary CCD in grey), order-aware UPGMA (red), order-aware UPGMA with scaled root height, (orange) and order-aware UPGMA scaled using SciPhy (yellow). In (**C**), we plot the total number of lineages ( y-axis) through time for each day (x-axis) for the trees reconstructed by each of the methods. In **D** and **E** we extend this comparison using simulated data, and additionally compare SciPhy to counterpart methods that do not model ordered editing, i.e. TideTree, and standard, order-unaware UPGMA (’UPGMA unordered’).

We find that SciPhy’s tree point estimate based on the conditional clade distribution (CCD) achieved the lowest median Phylogenetic Information (PI) distance to the ground true tree, indicating the highest reconstruction accuracy across methods (Fig. 4 D). Notably, TiDeTree performed comparatively well despite not having access to the true order of edit acquisition. However, we expect the performance gap between any method only modelling unordered edits and SciPhy to widen in scenarios where the edit distribution is more skewed—i.e. where some edits are much more frequent than others—since having access to the true order of edits becomes increasingly informative in such settings.

To evaluate the combined performance of tree topology and branch length estimation, we compared the estimated trees in terms of the weighted Robinson Foulds distance (wRF, Fig. 4 E). Here, SciPhy clearly outperforms all other methods. This advantage arises from SciPhy’s explicit model of sequential edit acquisition within a tape. In contrast, both TiDeTree and unordered UPGMA treat all sites as independent and are thus misspecified. Ordered UPGMA incorporates site ordering in the distance matrix calculation (i.e. a higher similarity comparing sites within tape sites is only achieved if the previous sites are identical). However, ordered UPGMA clustering only uses pairwise-distances and does not use the known ancestral state of each tape, nor does it use the knowledge of the editing process to inform branch length inference as SciPhy does (see Appendix Fig. 3 and 4 for more assessment of the accuracy under varying parameter regimes).

To showcase how the differences highlighted above on simulated data may translate to empirical analyses, we repeated the comparison between SciPhy and the order-aware UPGMA algorithm using the HEK293T dataset as input to both methods. Again, to investigate whether they recover similar ancestral relationships between cells, we used the Phylogenetic Information metric to compute pairwise distances between sets of SciPhy posterior trees and UPGMA, visualized in 4 coordinates (Fig. 4 A and see Appendix Fig. 10). Within this metric space, the UPGMA tree falls separate from the bulk of our inferred set of posterior tree topologies in half of the pairwise coordinates (coordinates 1-2, 1-3, 1-4, see Appendix Fig. 10 for all pairwise coordinates), indicating that both methods capture similar overarching topological features, but that SciPhy recovers distinct cellular relationships looking at a finer resolution. We find similar patterns of segregation between the SciPhy posterior and UPGMA trees using different distance metrics. Using the Robinson Foulds distance, the UPGMA topology shows an overlap with SciPhy posterior tree estimates in all but the first two pairwise coordinates (Appendix Fig. 8). Consistently, the Clustering Information distance showed separation between UPGMA and SciPhy trees in two out of 6 pairwise coordinates (coordinates 1-2, 1-4 in Appendix Fig. 9.

Next, we also evaluated SciPhy against the order-aware UPGMA with respect to branch length inference for the HEK293T dataset. UPGMA produces trees with relative branch lengths that are not anchored to an absolute time scale. In contrast, SciPhy calibrates branch lengths to absolute time, using the experiment’s duration as a temporal reference. For fair comparison, we adapted the UPGMA tree to include temporal information by scaling it in three different ways. First, we scale its root (the most recent common ancestor (MRCA) of all cells, and representing the timing of the first division event) to 25 days to match the experiment duration (labelled “UPGMA ordered”). However, it is likely that this first division event happened after the start of the experiment. Therefore, we also scaled the UPGMA root to the median root height of SciPhy trees (labelled “UPGMA ordered + root scaling”). Finally, we also estimate branch lengths using SciPhy on the fixed UPGMA topology (labelled “UPGMA ordered + SciPhy scaling”).

Adopting the same approach as before to visualize trees in pairwise dimensions but using the weighted Robinson Foulds distance, acounting for branch lengths, we observe that UPGMA trees diverge from trees in our posterior set in all pairwise coordinates (see Appendix Fig. 12). Given the topological similarities previously noted, this implies that UPGMA predicts branch lengths patterns distinct from those in trees from SciPhy’s posterior set.

To further illustrate these branch length differences, we display the total number of lineages through time (LTT) for each reconstructed phylogeny (Fig. 4 B). Notably, SciPhy posterior trees display a slower increase in the number of lineages compared to UPGMA. For example, 500 lineages are reached after 5.5 days according to the UPGMA tree (5.7 days in the scaled UPGMA) but only after a median of 8.8 days (95% HPD interval of [8.0, 9.7]) days under SciPhy. Similarly, in the UPGMA tree, the final number of 1000 lineages is reached 1.1 (median, 95% HPD interval [0.31, 2.1]) days earlier compared to trees estimated by SciPhy. As a result, there is no overlap in LTT between the SciPhy tree posterior and the UPGMA tree for most of the experimental duration.

Branch lengths are the key signal informing population dynamics in phylodynamic analyses. Given the divergence in estimated branch lengths between UPGMA and SciPhy, we investigated the impact on growth rate estimates. We compared growth rate estimates based on four input tree sets: (1) the posterior distribution of trees (”SciPhy”) (2) the UPGMA tree scaled to the full experimental duration (”UPGMA ordered”), (3) the UPGMA tree scaled to the median root height of SciPhy trees “UPGMA root”) and (4) the UPGMA tree with SciPhy-estimated branch lengths (”UPGMA + SciPhy”) (Fig. 4 C). The median growth rate inferred on fixed UPGMA trees (95% HPD interval of [0.54, 0.60] (2), [0.55, 0.60] (3) and [1.19,1.32] (4)) is higher than the one inferred under SciPhy. Notably, the growth rate inferred on the UPGMA topology with branch lengths scaled with SciPhy is the highest, leading to doubling times between 12.5 and 14.0 hours, which excludes reported values for this cell line.

Overall, these results highlight that, consistent with the differences underlined on simulated data, empirical trees reconstructed using SciPhy and UPGMA for the HEK293T dataset differ in topology, although the relative impact of these disparities is challenging to quantify in the absence of a ground truth for this empirical dataset. There are notable differences in the branching times, which can lead to qualitative differences in the estimated growth rates.

### 2.4 Estimating growth dynamics during gastruloid development

We analysed a dataset where DNA Typewriter was employed to record the development of gastruloid formation. The experiment began with a single mouse embryonic stem cell (mESC), which was cultured as a single colony on top of mouse embryonic fibroblasts over four days, after which it further expands as a spherical 3D culture for three days. Then, Chiron (CHIR) treatment is applied for one day, initiating symmetry breaking and elongation in the gastruloid. Finally, the cells grow for four more days when they are collected for sequencing (Fig. 5 A). Our analysis focuses on 780 cells, each of which contained the same set of 8 most frequent DNA tapes.

**Figure 5.**
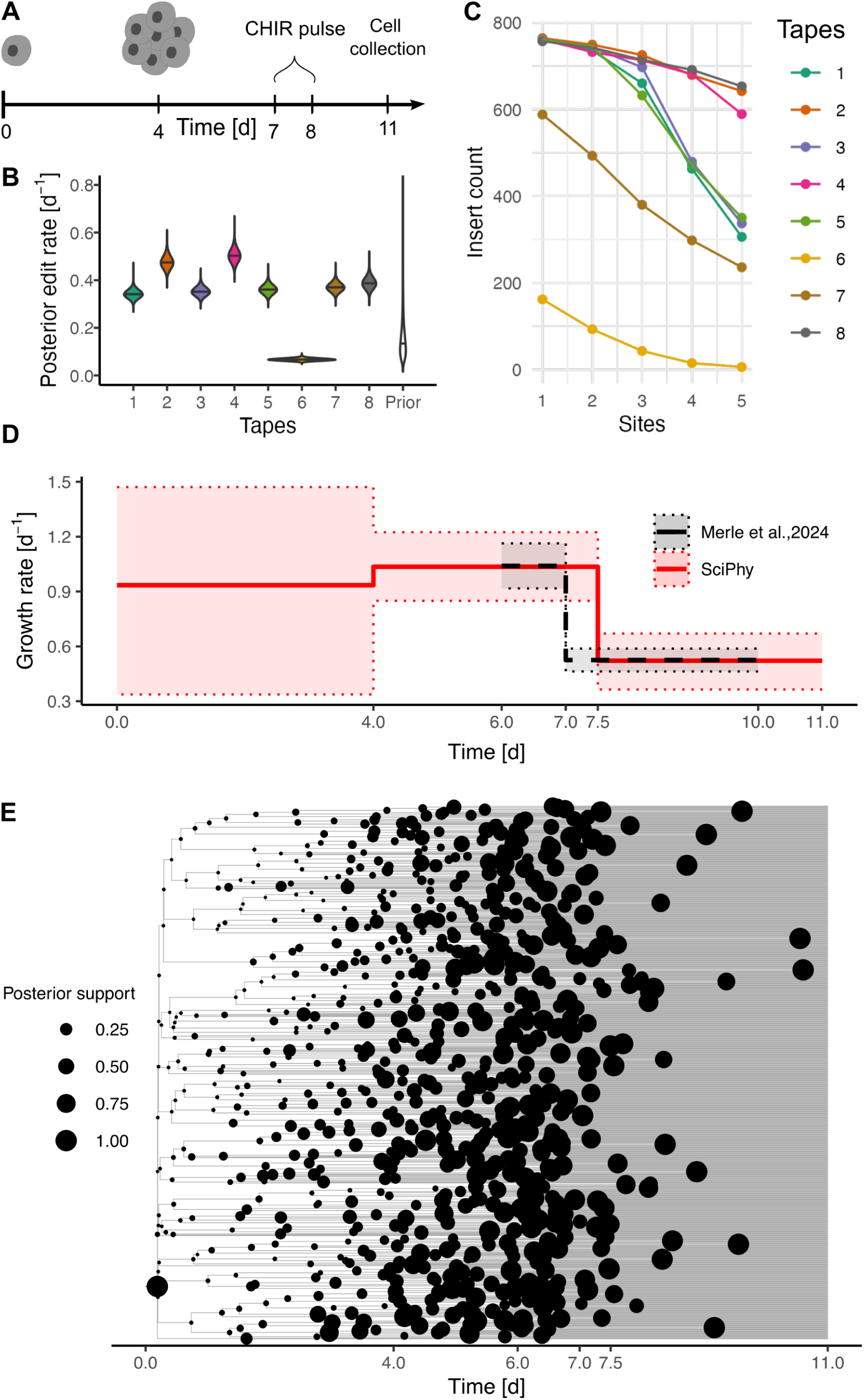
SciPhy estimates slow-down in growth rate upon CHIR stimulation. (**A**) We apply SciPhy to data collected from a monoclonal expansion experiment of gastruloid growth. Until day 4, cells grow to form an aggregate in which they continue growing until day 7, when a chir pulse is applied. At day 11 the cells are collected for sequencing. (**B**) We show the insert count for each tapes across all sites. In (**C**), we display estimated posterior edit rate per day for each tape. In (**D**) We show the estimated time-varying growth rate of the cell population (median and 95% HPD) against the growth rates (mean ±*σ*) measured in Merle et al. (2024), binned to match our time-intervals (see Methods 4.9). (**E**) Is the inferred CCD lineage tree inferred under SciPhy for this dataset.

Initially, we visualised the dataset by displaying the number of inserted sequences per site for each tape (Fig. 5 B). The data reveal differential editing across tapes. For instance, tapes 2, 4 and 8 exhibit high levels of editing across all sites, whereas the other tapes show a stronger decline in the number of inserts at the later sites. Notably, tape 6 is sparsely edited, with less than 200 cells being edited at the first site.

In addition to the overall number of inserts, the number of uniquely edited tape sequences provides more intuition about the information content in the different tapes. Thus, we also examined the number of unique sequences for each tape (Appendix Fig. 19). Interestingly, the most heavily edited tapes (2, 4, 8) show the smallest number of unique sequences, indicating that their editing saturated early in development when fewer cells were present. Conversely, a higher number of unique sequences were found in tapes 1, 3, 5 and 7, despite the fact that they were less edited. This could be due to editing being slower and thus lasting longer and marking more cells. Despite being the least edited tape, tape 6 shows a number of unique sequences comparable to the heavily edited tapes.

We apply SciPhy to estimate the editing model parameters, the time-scaled phylogeny and the dynamic growth rate over time. We find that the estimated clock rates vary among the different tapes (Fig. 5 C). Notably, tapes 2 and 4, which were suspected to be edited early based on the insert count analysis, also exhibit elevated editing rates based on our phylogenetic analysis. Additionally, the editing rate for tape 6 is notably lower compared to other tapes, consistent with its lower insert count. We further estimated the insertion probabilities for all inserts and find a highly unbalanced pattern (Appendix Fig. 20). Specifically, there is approximately a 50% probability of inserting the sequences “CTT” or “CAC”. This suggests that independent insertions of the same edit to the same tape is likely, which Sciphy can account for.

To estimate the growth rate of the gastruloid over time, we use a birth-death skyline model with varying birth and death rates. One advantage of the mechanistic nature of this model is that it allows to directly set the rate changes to specific time points aligned with experimental steps. For instance, we allow the rates to change after day 4, when the cells start to grow within an aggregate. Additionally, we allow for another change in rates after CHIR treatment, at day 7.5, leading to 2 change points in total (Fig. 5 D). During the expansion from a single-cell to aggregate, we estimate a median growth rate of approximately 0.9 *d*^−^1 (95% HPD of [0.34, 1.50]), which corresponds to a doubling time of 18.5 hours. Between days 4 to 7.5, the median growth rate is estimated to be slightly elevated to 1 *d*^−^1 (95% HPD of [0.85, 1.2]). After the CHIR pulse, we infer a significantly lower growth rate compared to the previous interval, with a median of 0.5 *d*^−^1 (95% HPD of [0.36, 0.67]). A decrease in growth rate after day 8 is also recovered when allowing rate changes at days 7 and 8 (before and after CHIR pulse, see Appendix Fig. 18 “3 change times”), suggesting that this observation is robust to different timeline setups. This slowing population growth can be visually appreciated from the time-scaled phylogeny (Fig. 5, E) where the number of new branching points (cell division events) decreases after day 8. We further performed various sensitivity analyses and found that our results remained robust across all settings (Appendix Fig. 22). As empirical validation of this pattern, we compare our estimates to previously reported growth dynamics of gastruloids recovered through imaging approaches (Merle et al. (2024), Fig. 5, D). The magnitude of the decrease in growth rate is in close agreement across both methods. However, our estimates suggest the decrease may have occurred 12 (Fig. 5, D) to 24 hours (Appendix Fig. 18) “3 change times”) later than reported by Merle et al. (2024). One possible reason for this discrepancy is our assumption of a constant editing rate throughout the experiment. If the rate of DNA tape editing declines over time, as observed with another sequential prime editor Chen et al. (2024), some cell divisions may be erroneously placed at later time points, resulting in the delayed decline in growth rate. Another factor could be differences in experimental protocols, as our dataset derives from monoclonal expansion, whereas Merle et al.’s was generated from an initial aggregate of several hundred cells. Despite the uncertainty regarding the exact timing of the growth rate decline, both datasets contain a signal for such a decline, suggesting a conserved response to CHIR signaling, independent of growth protocol.

In summary, our analysis reveals a pattern of slowed cell divisions following CHIR treatment in this gastruloid, confirming previous observations Merle et al. (2024). Importantly, this example showcases that lineage tracing data alone was able to encode signal for time-varying growth dynamics previously obtained manually through a combination of imaging and segmentation.

## 3 DISCUSSION

In this study, we introduced an editing model, edit sequence simulator and likelihood calculation tailored for lineage recordings based on sequential genome editing. Packaged as ‘SciPhy’ (**S**equential **C**as-9 **I**nsertion based **Phy**logenetics), our model is integrated into the BEAST 2 Bayesian Phylogenetics framework. This integration empowers SciPhy users to specify complex editing dynamics, for example varying editing rates across DNA Tapes as done here or along tree branches (using e.g. relaxed clock models Drummond et al. (2006), Drummond and Suchard (2010)) which was not done here. Additionally, it supports more nuanced models of cell development, allowing for cell-type dependent division and differentiation rates using multi-type models Vaughan et al. (2014); Kühnert et al. (2016) or time-varying rates Drummond et al. (2005); Stadler et al. (2013). The BEAST 2 platform is increasingly attracting developers interested in single-cell analyses (Seidel and Stadler (2022); Chen et al. (2022); Lewinsohn et al. (2022); Zwaans et al. (2025)) and we foresee that support in this area will continue to grow in the future.

Our evaluations of SciPhy on experimental datasets demonstrate its ability to capture complex editing dynamics, such as the presence of preferential insertions and editing rate variation between tapes, patterns that are prevalent in both sequential lineage datasets we analysed and those from other lineage tracing technologies Chow et al. (2021); Seidel and Stadler (2022); McKenna et al. (2016); Feng et al. (2021). As a mechanistic model, SciPhy can directly account for these factors and even use the signal from tapes being edited at different rates to inform branch length estimation. This ability to estimate branch lengths, and from them, cell population dynamics is consistent with a growing body of related work assessing properties of sequential editing data Mulberry and Stadler (2025); Pilarski et al. (2025).

We compared SciPhy to existing phylogenetic methods on both simulated and real data. On the simulated benchmark we showed that SciPhy outperforms all existing methods in terms of tree reconstruction accuracy and branch length estimation. We further compare SciPhy to ordered UPGMA clustering, which was used in both previously published analyses, on real data. UPGMA does not account for preferential insertions nor different edit rates for different tapes. We find that some—but not all—aspects of the tree topology are captured in both SciPhy posterior trees and the UPGMA tree. Notably, the branch lengths differ significantly, which we showed impacts the downstream inference of the cell population growth rate and is likely to impact other parameters one may wish to estimate from cell phylogenies in the future, such as cell-type differentiation rates.

For both datasets analysed in our study, we used phylodynamic models to estimate cell population growth based on SciPhy’s posterior distribution of timed trees. In HEK293T cells, this approach showed that sequential lineage tracing data alone contains signal for the growth rate characteristic of this cell line. When analysing murine gastruloid development, we estimated dynamic growth rates over experimentally informed time intervals and observed a significant reduction in cell growth after CHIR stimulation, consistent with previously reported dynamics Merle et al. (2024). This is also aligned with recent research aiming to resolve mechanisms of axis elongation during gastruloid development de Jong et al. (2024), suggesting that elongation occurs not through increased, localized cell divisions, but rather through active cell movement and differential adhesion. In both analyses, SciPhy allowed us to extract relevant and previously inaccessible insights from the lineage tracing data.

In our analysis of cell culture growth, we observed that the inferred editing rates for the 4-site tapes were consistently higher than those for the 5-site tapes. We used a statistical test to show this pattern is unlikely to arise under a null model of equal rates across all tapes. However, a limitation of our approach is that all inferences were performed using SciPhy, which assumes equal editing rates across sites. We present simulations that provide support for SciPhy estimating an average edit rate across sites for each tape, essentially allowing us to constrain the space of within-tape rate variation to scenarios consistent with the estimated per-tape average. Nevertheless, a model that explicitly allows for site-specific editing rates would enable more direct testing of such hypotheses and remains an important direction for future work.

While SciPhy offers promising avenues, it has some limitations. Tree space grows exponentially with the number of cells and current state-of-the-art Bayesian phylogenetic inference methods may not converge for trees with more than around 1000 tips. Bayesian phylogenetic inference requires repeatedly evaluating the likelihood to reach convergence. Thus, despite the likelihood calculation scaling linear with the number of tips, further speed-ups of the likelihood calculation can bring large benefits. We have achieved considerable improvements on the runtime of individual computations of the likelihood with caching and multi-thread parallelization (and provide estimates of the resulting combined, overall runtime for convergence of typical analyses in Appendix Fig. 1 and 2). However, the empirical data analysis on 1000 cells still required ∼15 days, and would benefit from further reduction in runtime. For future improvements, we aim to accelerate the likelihood calculation using software optimizations or approximations, and expedite convergence on large datasets by confining tree space and adapting recent advancements based on mutation annotated trees Varilly et al. (2025) that have been demonstrated to scale to 100’000 tip trees. Improvements to other components of Bayesian inference, such as, e.g., MCMC sampling methods, may also provide additional runtime reduction (Sherri et al. (2017), Hoffman et al. (2014)) but their applicability beyond continuous/real parameter contexts, such as the one encountered in phylogenetic inference, remains an open research problem.

Another limitation shared by all CRISPR-based barcoding technologies is that resulting lineage tracing datasets are typically lossy as a result of the sparseness of single cell RNA-seq and/or silencing of the regulatory elements that drive transcription of recording substrates (e.g. DNA Tapes). Jointly, these mechanisms result in the partial recovery of DNA Tapes per cell. For example, in the HEK293T cell culture dataset used in our study, none of the tapes were recovered for all sequenced cells, and there was variability in the rate of recovery of individual tapes. The relative contribution of either mechanism to this loss is unknown and for ordered lineage tracing data, state-of-the art lineage reconstruction approaches have thus far neglected cells with incomplete DNA Tapes. In our simulations, we found that trees reconstructed from incomplete alignments of DNA tapes are less accurate, but that parameters inferred jointly with those trees remain overall unbiased (see Appendix Fig. 24, for the editing and growth rates). However, ignoring such sources of loss may still bias tree or population parameter inference, by violating a common assumption of birth-death models of uniform sampling of the population of interest, e.g. if DNA tapes are simultaneously lost for groups of related cells. Related work for single-cell diploid data and CRISPR-based unordered data has shown that mechanistic models of error sources improve features of inferred lineage trees (Chen et al. (2022); Mai et al. (2024), especially relating to branch lengths. Follow up work should integrate a dropout model accommodating tape absence such that data loss and the resulting bias is avoided, and thoroughly investigate more complex patterns of loss. For the datasets analyzed in this study, sequencing error is unlikely to be a major issue, as the tapes are recovered using UMIs (any retained error would thus need to arise multiple times independently). For datasets relying on other sequencing techniques, extending SciPhy with explicit models of such error may, however, become relevant follow up work.

Many challenges remain in reconstructing accurate lineage trees, including refining experimental designs to improve signal quality in lineage tracing data. Future studies should systematically compare sequential genome editing with previous unordered methods, assessing both tree topology and the time estimates. Such comparisons will be essential to guide experimental efforts.

Lineage recorders based on sequential genome editing present a notable advancement as they do not only include a number of edits but also the order in which they occur. With SciPhy, we present a model that directly captures and uses this detailed information which we expect to lead to improved tree reconstruction compared to commonly-used clustering tools. We see great promise that sequential lineage recorders which can co-track cellular signals Chen et al. (2024) will be developed in the future. In parallel, SciPhy can be expanded to inform about the timing of such recorded signalling events in addition to the single cell trees, leading to developmental insight in ever-greater detail.

## 4 METHODS AND MATERIALS

### 4.1 Editing model for barcodes with ordered inserts

We present a model that captures the editing dynamics underlying the generation of sequential lineage tracing data using DNA recording systems such as the ones recently developed by Choi et al. (2022); Loveless et al. (2021). In these systems, cells carry stably integrated barcodes or “tapes”, which are contiguous arrays of CRISPR-Cas9 target sites. Within these tapes, target sites are sequentially made amenable to prime editing, resulting in the ordered accumulation of inserts. We consider an experimental setup starting with a single cell carrying several of these unedited tapes. As this cell and its offspring undergo proliferation, their tapes experience editing. At the end of the experiment, single cells are sampled and their tapes sequenced. Aligning sequenced tapes to their corresponding, unaltered tape reveals genetic modifications accumulated during the experiment. In the following, we detail the notations and assumptions we use to model this process. Consider the following setup. Each cell carries *k* tapes that independently undergo editing. Each unedited tape is an array of N target sites, numbered from 1 to N. Each site can acquire a single insertion or edit, with *M* different inserts available.

Our model is defined as follows:

1. Initially, only a tape’s first site is susceptible to editing. Subsequent sites are edited sequentially: editing at a site *i*_1_*_<i_*_≤*N*_ is only possible if its preceding site *i* − 1 is already edited.
2. Editing is a two-step process :

a. Step 1: The Cas9 prime editor introduces cuts at the target sites in a time-dependent manner at rate *r*, called the editing rate or clock rate. This means editing follows a bounded Poisson process which stops once *N* edits have occurred.
b. Step 2: pegRNA mediated insertions following cut events are instantaneous. The set Γ of *M* possible insertions is predefined and denoted as:

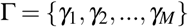 Given a cut event, the probability of adding insert *γ_i_* is *f_i_*(with *η* := ( *f*_1_*, f*_2_*, …, f_M_*)), allowing for some insertions to have higher probabilities than others. We additionally assume that insertions operate independently - an insertion at a previous site does not influence the insertion probability at the currently active site.
3. Editing of any target site is irreversible.

From the model definition, if follows that the number of edits introduced in each independent tape is Poisson distributed. However, the distribution of the total number of edits per cell is not guaranteed to show features of a classic, unimodal Poisson distribution, because it is also influenced by the shared ancestry of the sampled cell population and its growth patterns (see Appendix Fig. 13 and 14).

Formally, for an array of length N, this model of editing is a continuous time Markov chain on state space Ω of all possible insertions and parameters *r, η*. These states are all possible finite sequences of length *n* ≤ *N*, or n-tuples of ordered insertions, that we denote as:

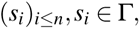

where subscript *i* denotes the insertion site in the tape, i.e. its order. Note that the empty tuple 0/ represents the unedited state or starting state for all k tapes sampled per cell and the fully edited tape is an absorbing state of the Markov chain.

Let *Y_t_* be the state of an array at time *t* after the start of the experiment. Given an array state *Y_t_* = *a* and its state after a time interval Δ*t*, *Y_t_*_+Δ*t*_ = *b*, we denote the tuple of inserts added to *a* to result in *b* as *c*. Specifically, *c* represents the elements in *b* that are not in *a*, which can be expressed as *c* = *b* \ *a*. The size of this tuple, denoted as |*c*|, corresponds to the number of inserts introduced between states a and b.

### 4.2 Transition probabilities

We denote the transition probability between states a and b under the model introduced above as *P_a,b_*(Δ*t*; *η, r*) = *P*(*Y_t_*_+Δ*t*_ = *b*;*Y_t_* = *a, η, r*). In the following, we derive this transition probability. In step 1, we calculate the probability that a number of |*c*| new edits have been inserted. In step 2, we calculate the probability that those edits correspond exactly to the tuple c.

**Step 1** To derive the probability of introducing |*c*| edits, we distinguish two cases. In case 1, the number of new inserts |*c*| does not lead to saturation in the array (|*a*| + |*c*| = |*b*| *< N*). Then, the probability of |*c*| editing events happening at rate *r*, in time Δ*t* is given by the Poisson distribution (by model definition 2a):

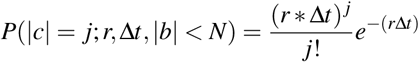

In case 2, introducing |*c*| inserts leads to saturation of the array (|*a*|+|*c*| = |*b*| = *N*), i.e reaching position N in the tape. We calculate the probability of this event by targeting its complement, namely the probability that less than |*c*| inserts were introduced (again, by assumption 2a):

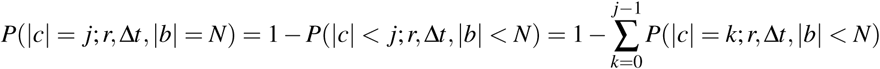

Combining both cases, the probability of introducing |*c*| edits in an array of length N given an editing rate *r* is:

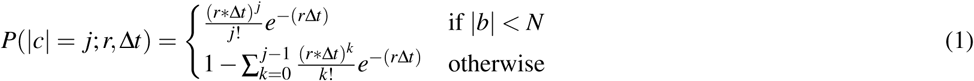

**Step 2** Let us assume that the |*c*| insertions that occurred from a to b are exactly the tuple *c* = (*s_i_*)*_i_*_=|_*_a_*_|+1_*_,…,_*_|_*_b_*_|_*, s_i_* ∈ Γ. Conditional on observing exactly |*c*| inserts, the probability of realising the specific tuple c equals the product over the insertion probabilities *f_l_* for each element of c, (by independence, see Assumption 2b):

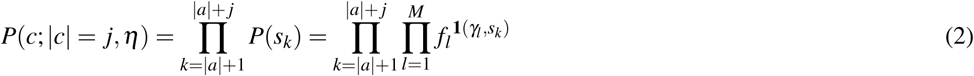

Note that the indicator function **1**(*γ_l_, s_k_*) serves as a one-hot selector, keeping the insertion probability *f_l_* only when the edit *γ_l_* matches the observed symbol *s_k_* in the tuple c. This construction collapses the inner product to the single insertion probability corresponding to the realised edit at position *k*.

**Step 3** Combining the steps above, the transition probability between any two states a and b in time interval Δ*t* given model parameters *r* and *η*, is obtained as follows (and as exemplified/described in Fig. 6):

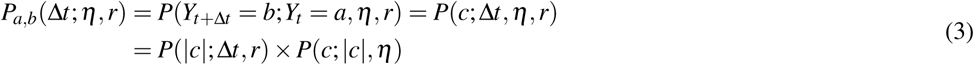

where *P*(|*c*|; Δ*t, r*) was derived in Step 1 and *P*(*c*; |*c*|*, η*) in Step 2.

**Figure 6.**
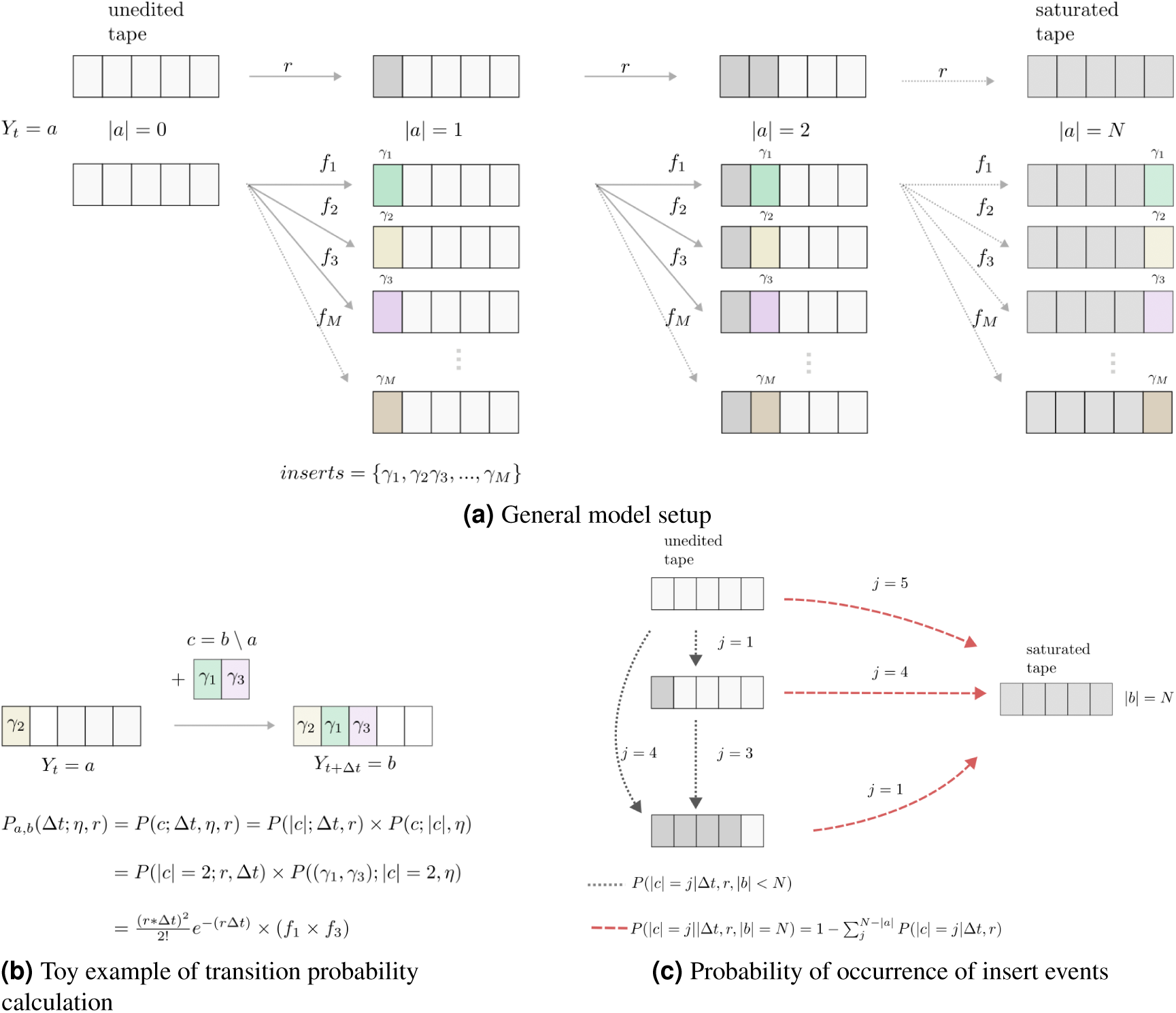
General description of model components used to calculate transition probabilities in SciPhy. An overview of SciPhy’s editing model for a tape with N=5 target sites and its parameters is shown in (**a**). Target sites within tapes (rectangles in the tape) acquire inserts (colored boxes, inserts *γ*_1_ to *γ_M_*) with probability *f*_1_ to *f_M_* following Cas9 mediated nicks at rate *r*. Upon editing these sites are inactivated (grey boxes). (**b**) is a toy example for the calculation of the transition probability for a tape acquiring 2 edits (*γ*_1_ and *γ*_3_). In (**c**) we showcase different scenarios for the calculation of the probability of a given number of inserts to be added into a tape, distinguishing between cases where a tape gets saturated with inserts (red arrows pointing to the right) or not (grey arrows pointing down).

### 4.3 Likelihood calculation

The editing model defined above can be used to calculate the likelihood of the model parameters (*r, η*) and the cell phylogeny *T* given the tape data *D* at the tips, *Lik*(*r, η, tree*; *D*) = *P*(*D*; *r, η, tree*). Let *T* be a tree with *n* tips and *n* − 1 internal nodes. Tho oldest of these nodes is the root node, being the most recent common ancestor of all sampled cells. Additionally, we introduce a stem branch preceding the root, connecting the root node to the start of the experiment, or origin node. We number the leaf nodes from 1*, …, n* arbitrarily, and the internal nodes from *n* + 1*, …,* 2*n* with increasing distance from the present. Each cell may have *k* tapes. Since we assume that tapes are edited independently from each other, we calculate the probability density of tape *i* given the tree and *r, η* separately for *i* = 1*, …, k*. The likelihood is then the product over the *k* probability densities. In what follows, we thus consider one particular tape.

We first obtain the sets of possible tape states at each internal node given the data at the tip nodes. The likelihood of the tree is then calculated by summing over all possible state transitions between the nodes. This summation is performed through dynamic programming, going from the leaf nodes toward the origin of the tree; the employed particular algorithm is Felsenstein’s pruning algorithm Felsenstein (1973).

Hence, we first determine all possible states at internal nodes. We restate that editing is irreversible and ordered (model definitions 1, 3). Therefore, for any sequence at a leaf node, ancestors can only harbor either the same or fewer edits. For any leaf node *m*, let *v^m^* be its tape with elements 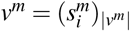 where |*v^m^*| is the number of edits in *v^m^* as defined in the editing model section. The set of its possible ancestor tape states *A_m_* is the set of all subsets of edits from *v^m^*, akin to a power set on ordered tuples:

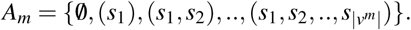

For an internal node *i* of tree *T*, it follows that the set of possible states *A_i_* is the intersection of possible states of its two children nodes *k* and *j*, *A_k_* and *A_j_* (note that for the leaves, we initialize with all possible ancestor states). This is a result of editing being irreversible (assumption 3): an edit present in one child node but not the other cannot have been present in their parent node. Conversely, an edit present in both children nodes, can either have been present in the parent node or have been acquired by both children independently, therefore it should be included in the set of possible states. Consequently, the possible state set for node *i*, *A_i_* is obtained as the intersection of the possible states of its children’s *k, j*, *A_k_* and *A_j_*, recursively up the tree:

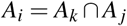

noting that the leaves are initialize with all possible ancestor states.

Given the possible states at the internal nodes, we calculate the probability density *P*(*D*|*T, r, η, A*_2*n*_) of the tape data given the tree and model parameters following notations in (Seidel and Stadler (2022)). To match assumption 1, we assume the unedited state at the origin node *A*_2*n*_ = {0/)}.

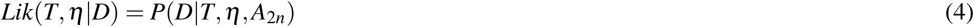

The transition probabilities from every parent node *π_i_* to its child node *i* along their connecting branch with length *τ_i_* are calculated using Eq. (3). Remember that *A_i_* is the set of all possible states at node *i*. Any specific state within *A_i_* is denoted as *a_i_*. To account for all possible transitions, we sum the probabilities over all possible states at each internal node of the tree.

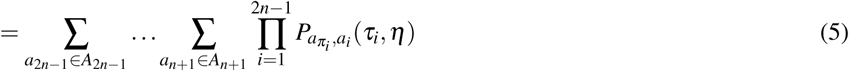

Using Felsenstein’s pruning algorithm, we ‘move the summations down the tree as far as possible’ (Felsenstein (2004)), meaning we perform the summations recursively up the tree, saving computations:

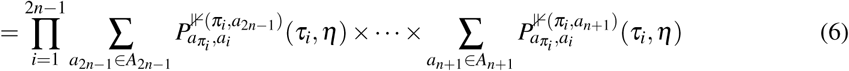

As stated above, this calculation of *P*(*D*|*T, r, η, A*_2*n*_) is performed for each tape. Multiplying the resulting probability densities for each tape leads the targeted likelihood function.

For a set of *k* tapes, where every tape is of length N on a tree with *n* tips, we perform n-1 operations to calculate the set of ancestral states at every internal node. Performing the intersection at an internal node scales with the size of the set, N. Hence, computing the ancestral states scales in O(knN).

Additionally, for each of the k tapes, we calculate the transition probabilities at every one of the n-1 internal nodes. At each node, we consider a maximum of (N+1)x(N+1) potential state transitions. Hence, our likelihood computations without optimization scales in O(knN^2^) (In current technologies N is relatively small, *N* ≈ 5). Note that the computation over the different k tapes is conveniently an ‘embarrassingly parallel’ problem.

Overall, the calculation is polynomial in the size of the input, *O*(*knN* + *knN*^2^) = *O*(*knN*^2^). For a fixed *N* and *k*, the runtime scales linearly with the number of tips in the tree, meaning the number of cells in the dataset.

### 4.4 Implementation

We implement the above editing model alongside an edit sequence simulator based on the feast package Vaughan (2024) and a likelihood calculation as a new package –SciPhy– for the phylogenetic inference platform BEAST 2. Specifically, we introduce a new substitution model class, SciPhySubstitutionModel, which replaces standard models such as Jukes–Cantor to compute transition probabilities between edit states. We also extend the Generic TreeLikelihood class in BEAST 2 with the SciPhyTreeLikelihood class tailored to sequential editing. This implementation leverages domain-specific properties of lineage tracing data—for example, that the set of possible ancestral sequences is known. Upon installing the SciPhy package, these components are directly available to BEAST 2 users via both the command-line interface and the graphical interface BEAUti.

BEAST 2 performs phylogenetic inference by using Markov Chain Monte Carlo (MCMC) to quantify the posterior distribution of the lineage tree and model parameters given a set of tape sequence alignments, where each tape may be specified to have independent properties (such as tape length/truncation, and editing features). During the MCMC, tree operators modify the tree to explore all possible topologies that could explain the data. As some of these changes are local, the likelihood for subtrees that are not modified remains unchanged. We make use of that property by implementing caching, where the likelihoods for subtrees are saved at each internal node to further speed up calculations (3.5-fold speed up on a personal computer running OSX on Intel(R) Core(TM) i5 Dual-Core @2.30Ghz). Additional speed-up is achieved by computing the likelihood for each tape alignment in parallel, which is easily achieved with existing components in BEAST 2 that support threading. These software optimizations lead to an overall 8-fold speed up relative to the naive implementation for an analysis involving 1000 sequences using 6 computing cores on the “Euler” scientific computing cluster. Using this setup, we provide realistic estimates for the average runtime required for convergence on smaller datasets from the validation study, and when varying the number of editing sites in Appendix Fig. 1 and 2.

### 4.5 Validation

We employed well-calibrated simulations (Dawid (1982)) to ensure the correct implementation of our Bayesian method SciPhy. We further used this simulation setup to assess the amount of information in the simulated data regarding parameters of our model and key features of the lineage tree.

We simulated 100 phylogenetic trees under a birth-death sampling model (Stadler et al. (2013)) with parameters provided in Tab. 2.

**Table 2.**
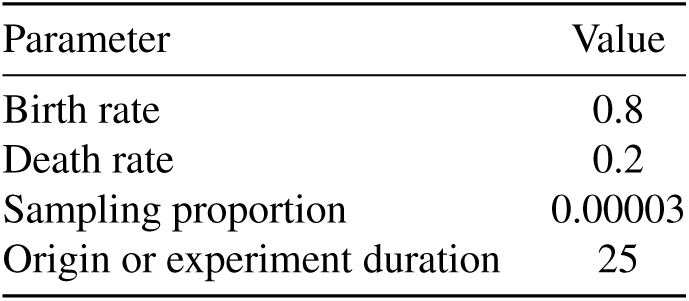
Parameters used for simulating a phylogenetic tree under a birth-death sampling model.

| Parameter | Value |
| --- | --- |
| Birth rate | 0.8 |
| Death rate | 0.2 |
| Sampling proportion | 0.00003 |
| Origin or experiment duration | 25 |

We simulated alignments of tapes along each of the 100 trees. Firstly, we drew the rate of edit introduction (or clock rate) and edit probabilities to generate 13 distinct edits from a defined distribution (Tab. 3). Then we simulated the editing process of 10 independent tapes along each phylogenetic tree under these parameters. The resulting alignments of tapes were then used as input data for SciPhy to infer the phylogenetic tree as well as the rate of edit introduction and edit probabilities, using the same distributions (i.e. Tab. 3) as a prior. For all parameters, we used an Effective Sample Size (ESS) value of 200 as threshold to ensure convergence of the MCMC chains. Using these outputs, we first compared the 100 simulated (true) parameters and trees with their estimated counterparts. Specifically, we chose tree height, tree length and tree balance as summary statistics for comparison of the lineage trees. Tree balance was calculated using the B1 index, as implemented in package TreeStat2 Drummond (2025). For each dataset, we additionally constructed a point estimate of the lineage tree from the sampled posterior set of trees using the conditional clade distribution (CCD) algorithm (with the CCD0 parameterization, Berling et al. (2025)) and assessed accuracy in lineage reconstruction by reporting the Phylogenetic Information (PI) distance to the true tree Smith (2020), compared to a randomly reconstructed tree obtained by simulation under the birth-death sampling model. We test the difference between distances obtained with SciPhy and the random reconstruction using a paired t-test.

**Table 3.**
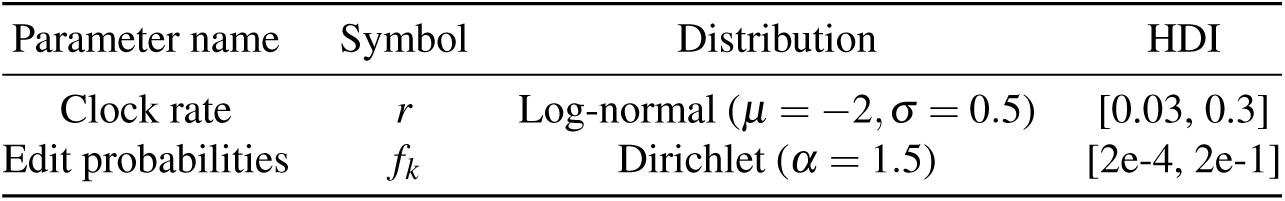
Distributions for the editing model parameters used in the validation.

We chose the distributions over the editing model parameters to be representative for a broad range of editing dynamics. The clock rate distribution was selected to result in between 1 to 8 insertions for the experiment duration. We used a Dirichlet distribution with a concentration parameter of 1.5 to draw the edit probabilities. This allows most edits to occur at similar, low probabilities while a few edits could occur more frequently.

For each MCMC inference, SciPhy was run for at most 10^9^ iterations to estimate the clock rate, edit probabilities and lineage tree, keeping all other parameters fixed to the truth (i.e. the value of the simulation). We used an ESS value of 200 for all parameters as threshold to ensure convergence of the MCMC chains. We then discarded the initial 10% of the iterations as burn-in. Lastly, we calculated the coverage as the proportion of times the simulated (true) value fell within the estimated 95% HPD (Highest Posterior Density) interval. For each of these datasets, we estimate the total runtime necessary for convergence by weighting the total empirical runtime by the ratio between the final obtained ESS and the desired minimum ESS of 200, as reported in Appendix figure 1. For the first simulated lineage tree, we repeat this estimation for tapes harboring an increasing number of editing sites, as reported in Appendix figure 2.

We assess the effect of alignment sparsity on the accuracy of phylogenetic reconstruction using SciPhy by generating incomplete alignments as input data for the same inference setup as used in the validation study. We consider two main mechanisms leading to tape loss: heritable loss of tapes at a constant rate through time, capturing e.g. transgene silencing, and tape dropout at sequencing, assuming that tapes will fail to be captured by the sequencing pipeline with a given probability. We simulated the editing process with tape loss for 20 independent tapes along each phylogenetic tree, and chose wide distributions for the rate of heritable loss and dropout probability, such that we would expect between 9.7 and 88% of sequenced tapes to be missing per simulated alignment (see Appendix Tab. 7). This led to an average of 11 tapes recovered per cell over all simulations (see Appendix Fig. 26). For each of these lossy tape alignments, we generate input alignments by constructing the largest complete set of tapes without missing data that include at least a third of all cells (“Filtered lossy data”). We used these alignments to infer the phylogenetic tree for the subset of cells, as well as the editing and growth parameters, and assessed the accuracy in lineage reconstruction as described above.

### 4.6 Methods benchmark

We reuse nine datasets from the validation study, sampling evenly across tree sizes by categorizing datasets into three groups: small (0–150 tips), medium (150–450 tips), and large (450–650 tips). From each category, we select three datasets to ensure balanced representation across tree sizes in our benchmark. For SciPhy, we reuse the previously inferred results from the validation study. We run the ordering-aware UPGMA as done before Choi et al. (2022), as well as standard UPGMA using hierarchical clustering with average linkage. TiDeTree (version 1.0.2) Seidel and Stadler (2022) is run using the same prior distributions as used for SciPhy. Additionally, for each dataset, we simulate a random tree under a birth–death sampling model using TreeSim (version 2.4) Stadler (2015), conditioning on the number of tips to match that of the dataset. We report the normalized phylogenetic information distance using TreeDist (version 2.9) Smith (2022).

### 4.7 Phylodynamic analysis of HEK293T expansion

We use SciPhy for the analysis of the publicly available dataset (i.e. taken from https://github.com/shendurelab/DNATickerTape, Supplementary File 2 DataTableMOI19.csv) generated as described in (Choi et al. (2022)). This study uses genetic tapes that contain 5 target sites for insertion, where each insertion step consists of the introduction of a trinucleotide character. We process the tape sequences as follows. We first retain all 3257 cells containing all 13 most frequent tapes for subsequent analysis, as done before. Out of these 13 tapes, 3 (barcoded with ‘TTCCGTCA’,’TGGTTTTG’ and ‘TTCACGTA’) are found to possess less than the 5 original editable target sites, respectively 4 (’TTCCGTCA’,’TGGTTTTG’) and 2 (’TTCACGTA’) sites. We refer to these tapes as “truncated”. This procedure yields a dataset containing 19 different possible trinucleotide insertions. We create 3 random subsets of this data, each consisting of 1000 cells and perform phylodynamic analysis on each subset.

We use SciPhy to model the editing dynamics in this filtered dataset. We assume that each of the 13 tapes can accumulate edits at an independent rate, while insertion probabilities are shared across all tapes. We place prior distributions on the parameters of the editing model as described in table 4. As no quantitative information on the editing dynamics is available for this experimental setup, we use broad priors that capture a variety of experimentally plausible values (as in the validation part). For example, the clock rate prior spans between 1 and 10 edits per tape over 25 days. As each tape has a maximum of 5 sites, 10 edits correspond to the scenario of early saturation. The prior is centered on 5, the desired edit count. The prior on the edit probabilities is a Dirichlet prior with concentration 1.5, chosen to reflect expected asymmetries in editing outcomes observed in similar non-sequential recorders Gong et al. (2021).

**Table 4.**
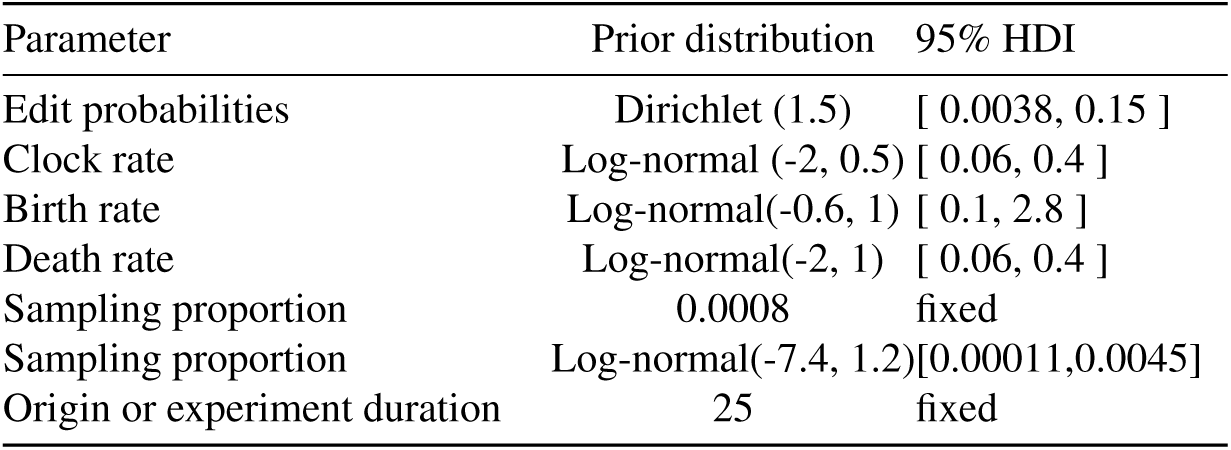
Priors used for analysis of the HEK293T dataset using SciPhy and the birth-death sampling model.

| Parameter | Prior distribution | 95% HDI |
| --- | --- | --- |
| Edit probabilities | Dirichlet (1.5) | [ 0.0038, 0.15 ] |
| Clock rate | Log-normal (-2, 0.5) | [ 0.06, 0.4 ] |
| Birth rate | Log-normal(-0.6, 1) | [ 0.1, 2.8 ] |
| Death rate | Log-normal(-2, 1) | [ 0.06, 0.4 ] |
| Sampling proportion | 0.0008 | fixed |
| Sampling proportion | Log-normal(-7.4, 1.2) | [0.00011,0.0045] |
| Origin or experiment duration | 25 | fixed |

Being sampled from a monoclonal culture, cells are expected to undergo exponential growth and to not acquire any specific population structure. Consequently, we choose to infer the dynamics underlying the growth of this cell population using the birth-death sampling model for constant and piecewise constant rates (Stadler et al. (2013)). We set priors on the birth and death rate (Tab. 4) that are very broad; leading for instance to between 12 and 2.5 ∗ 10^30^ cells at the time of sequencing, which broadly contains the reported number of cell (1.2 ∗ 10^6^). We additionally input known experimental parameters by setting the origin time to the experiment duration of 25 days and fixing the sampling proportion to 1000*/*1.2*e*6 = 0.0008. In the piecewise constant analysis, rates were allowed to vary every 2 days, yielding a timeline of 13 epochs where an Ornstein-Uhlenbeck (OU) prior is applied to the birth and death rates, and where we apply a Log-normal(0.5, 0) hyperprior on parameters *σ* and *ν* of the OU process.

For each subset of the data, we run 3 MCMC chains for at most 10^9^ iterations. We assess convergence by confirming that the three chains are sampling the same stationary distribution, with an ESS value per chain *>* 200 for all parameters of interest and visually in Tracer (Bouckaert et al. (2014)). We discard 10% burn-in and combine all 3 chains using LogCombiner (Bouckaert et al. (2014)). The combined ESS and *r*^ values for the posterior probability of all analyses of this dataset are reported in Appendix table 8.

To test whether the ordering of 4x and 5x tapes is unexpected under the assumption of equal editing rates for all tapes, we compute the probability of the observed (i.e. that k of the three tapes with the fastest editing rates are 4x tapes, one-sided test) using the hypergeometric distribution. We model the setup as follows: We have a total of 12 tapes (N=12), of which 3 tapes (K=3) have 4 sites and 9 tapes have 5 sites. If we draw the three tapes with the highest rates (n=3), we want to calculate *p*(*k*), the probability that k of these tapes are of type 4x. We do this using the standard hypergeometric distribution:

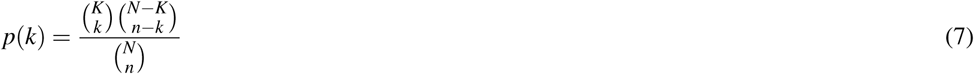

To compare the UPGMA tree to SciPhy’s estimated trees on real data, we use the R package TreeDist (Smith (2022)) to provide mappings of tree sets for four dimensions, pairwise. For this, we extract a random subsample of 1641 trees inferred under SciPhy to calculate distances to the UPGMA topology. We do so using three metrics for topological comparisons, namely the Robinson Foulds distance, the Phylogenetc Information distance, and the Clustering Information distance. Further we employ one metric that also incorporates differences in branch lengths, namely the weighted Robinson Foulds metric. For each metric, we perform multidimensional scaling to map these distances in lower dimensions using principal coordinate analysis (Gower (1966)). Out of all 12 dimensional mappings obtained, For each of these mappings we choose the number of dimensions required to faithfully visualize tree distances based on the product of the trustworthiness and continuity metrics as described by Smith (2022). For all mappings we find that 4 or more dimensions provide acceptable levels of distortion (trustworthiness x continuity ¿ 8.5). These first 4 dimensions, which we label 1 to 4 in our plots, are used for reference but do not represent interpretable features of analyzed trees.

### 4.8 Gastruloid experiment

#### Culturing and genetic engineering of the mouse embryonic stem cells (mESCs)

We adopted a standard protocol for culturing mESCs (Beccari et al. (2018)), which is described in detail in Regalado, Qiu et al. (Regalado et al. (2025)). Briefly, mESCs were cultured on a gelatin-coated plate using the Serum-based medium, which is GMEM (Gibco) supplemented with 8% Fetal Bovine Serum (FBS, Biowest), 8% KnockOut Serum Replacement (KSR, Gibco), 1X Glutamax (Gibco), 1X MEM non-essential amino acids (Gibco), 1 mM Sodium Pyruvate (Gibco), and 0.1 mM beta-mercaptoethanol (BME, Gibco). Gelatin-coated plates were prepared by making 0.2% (w/v) gelatin solution (Sigma, G1393) and autoclaving the bottle, which was applied to each culturing well (1 mL per well within a 6-well culturing plate) and incubated in a tissue culture incubator set to 37 °C with 5% CO2 for at least 30 minutes. After the coating, the gelatin solution was aspirated from the plate immediately before depositing mESCs. Cells are grown in the stem-cell-designated incubator (set to 37 °C with 5% CO2) and biosafety cabinet to avoid cross-contamination.

To allow DNA Typewriter-based lineage recording, mESCs were genetically engineered with piggyBac transposition system that integrated: 1) PB-TAPE, a transposon library including TAPE-targeting epegR-NAs (random NNNGGA insertions) expressed from U6 promoter and GFP transcript including a DNA Tape sequence in its 3’-UTR, 2) PB-PEmax, a transposon including Prime Editor (PEmax) expression cassette with constitutively expressed Puromycin resistance gene (PuroR), and 3) PB-mCherry, a transposon that expresses mCherry as a high-integration selection carrier. DNA Tape is structured to include the T7 promoter sequence, a 12-bp Tape-specific degenerate barcode (TapeBC, NNNNNAANNNNN), 6xTAPE capable of six sequential insertions, and 10X Genomics Capture Sequence 1 (CS1) to aid recovery of DNA Tape during scRNA-seq.

These transposon vectors and transposase plasmids were mixed with the 73:20:2:5 mass ratio (PB-TAPE:PB-iPEmax:PB-mCherry:Super piggyBAC transposase expression vector, SBI) then transfected using Lipofectamine 2000 (ThermoFisher) protocol (4 µg DNA per 1 million cells in 6-well plate). Transfected cells were cultured for 5 days and selected under 2 ug/mL Puromycin for another 5 days to ensure integration of PB-iPEmax. Selected cells were dissociated into single cells and flow-sorted by mCherry expression (from PB-mCherry), then plated at low seeding density for monoclonal gastruloid induction.

#### Monoclonal gastruloid induction and single-cell RNA-seq profiling

We used the monoclonal gastruloid induction protocol, as described in Regalado et al. (2025). Briefly, we cultured genetically engineered mESCs on a layer of Mouse Embryonic Fibroblast (MEF) cells at a low seeding density, resulting in the formation of monoclonal colonies across the culturing plate after five days. On Day 4, the mESC colonies were selectively removed from the MEF layer using Collagenase IV treatment and transferred to a non-adherent 10-cm plate in the differentiation medium (NDiff227, Takara), spontaneously forming 3D spheroids. After 24 hours, 3D spheroids or aggregates were manually picked into 96-well plates (non-adherent, round bottom) and subsequently used for the downstream gastruloid induction protocol Moris et al. (2020), including pulsing with 3 µM CHIR-99021, subsequently described as CHIR pulse, for 24 hours, followed by three days of culturing in NDiff227 medium until the harvesting of gastruloids for single-cell profiling (dissociated and processed following the conventional protocols: Cell Preparation guide, CG00053 Rev C, and User Guide for Chromium Next GEM Single Cell 3’ HT Reagent Kits v3.1, Rev D). Resulting libraries are sequenced on NextSeq2000 (Illumina).

Next, we processed the sequencing reads of DNA Tape captured in the scRNA-seq experiment. We extracted cell-specific barcode (CellBC), UMI added during cDNA synthesis, TapeBC, and 6 InsertBC from each DNA Tape sequencing read, and removed reads with less than 2 UMI associated with particular CellBC-TapeBC-6xInsertBC combinations. In the resulting reads, if there were multiple 6xInsertBC read patterns recovered per CellBC-TapeBC combinations, 6xInsertBC with the highest read counts were retained, resulting in a table of CellBC, TapeBC, and 6xInsertBC sequences. This table was used for the lineage tree reconstruction.

### 4.9 Gastruloid analysis

We use SciPhy to analyse DNA Typewriter tapes derived from the experiment described above. Similarly to the analysis of the HEK293T dataset, the full set of sequenced cells was filtered for cells where 8 of the most frequent tapes could be recovered by sequencing. This process yielded a set of 780 cells retained for our phylodynamic analysis, where 42 different trinucleotide characters were found to have been inserted into the tapes. We use SciPhy to model the prime editing process throughout the experiment for the DNA Typewriter tapes, where, as previously, each of the 8 tapes acquires edits at an independent rate, but where insertion probabilities for the 42 trinucleotide characters are shared across the tapes. We use the same prior distribution on the insertion probabilities as in the cell culture analysis, based on the expectation that editing outcomes are not uniformly distributed. For the clock rate, we adjust the prior to reflect the shorter 11-day time frame of the gastruloid experiment. The 95% highest density interval again corresponds to an expected range of 1 to 10 edits per tape, with the median aligned with the desired outcome of 5 edits (see Table 5 for prior distributions).

**Table 5.**
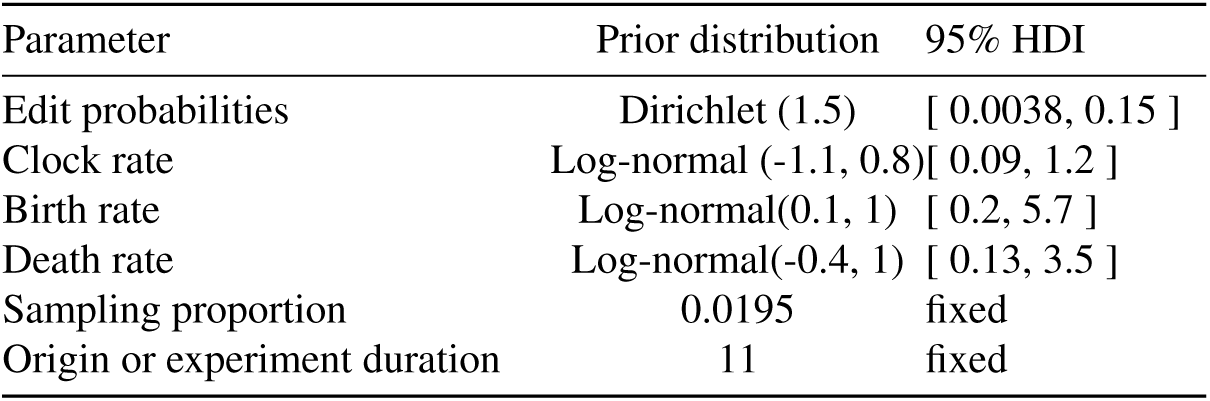
Priors used for the analysis of the gastruloid dataset using SciPhy and the birth-death sampling model.

| Parameter | Prior distribution | 95% HDI |
| --- | --- | --- |
| Edit probabilities | Dirichlet (1.5) | [ 0.0038, 0.15 ] |
| Clock rate | Log-normal (-1.1, 0.8) | [ 0.09, 1.2 ] |
| Birth rate | Log-normal(0.1, 1) | [ 0.2, 5.7 ] |
| Death rate | Log-normal(-0.4, 1) | [ 0.13, 3.5 ] |
| Sampling proportion | 0.0195 | fixed |
| Origin or experiment duration | 11 | fixed |

We study the growth dynamics that drive the formation of this gastruloid by fitting a birth-death sampling model with piecewise constant rates. We hypothesized that dynamics that underlie gastruloid formation are predominantly driven by experimental steps that lead to (1) aggregate formation, (2) aggregate growth, (3) Chiron-pulse driven (4) gastruloid elongation. Therefore, we allow birth and death rates to vary between these experimental steps, leading to two variants of the analysis:

1. Variant 1: considers birth and death rate changes between the steps of aggregate formation and gastruloid elongation only, leading to 2 change points, at days 4 and 7.5, shown in main text, Fig. 5.
2. Variant 2: considers birth and death rate changes between all steps of gastruloid growth, including the CHIR pulse, leading to 3 change points at days 4, 7 and 8, shown in Appendix Fig. 18.

We set broad priors on the birth and death rate, consistent with a growth leading to up to ∼ 10^7^ cells, and additionally fix the sampling proportion and origin time as known experimental parameters. Rate variation between different epochs of the experimental timeline in both variants of the analysis was modelled using an Ornstein-Uhlenbeck (OU) with a Log-normal(0.5, 0) hyperprior on parameters *σ* and *ν* of the OU process. All detailed priors are listed in Table 5.

The inference was performed as follows: we ran 3 MCMC chains on the dataset for at least ∼ 10^9^ iterations or until convergence. We assessed convergence, and combined all 3 chains as previously described (see above). The combined ESS and *r̂* values for the posterior probability for our analysis of this dataset are reported in Appendix table 8.

To evaluate our estimates of the growth through time against empirical measurements, we use a time series of the total cell count for gastruloids reported in Merle et al. (2024)(Extended Data Fig 2b). We calibrated their experimental timeline to match ours by aligning the day of CHIR application to day 7. Assuming exponential growth through time (as previously done in Merle et al. (2024)), we calculate the growth rate in each time interval *t_d_* (in days) as follows. Given a cell count *N*_0_ at time *t*_0_ and *N*_1_ at time *t*_1_ = *t*_0_ + *t_d_*, the growth rate *g_d_* on interval *t_d_* is expressed as:

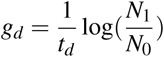

For comparison to our estimates, we calculate the growth rate over time periods that most closely match our chosen intervals for inference. For example, in Fig. 5 D, the rates are calculated between day 7 and day 10 to match our last time period, between 7.5 and 11 days.

### 4.10 Data availability

Raw sequencing data and associated processed data for the monoclonal gastruloid analysis have been uploaded to the Gene Expression Omnibus (GEO) and are available under accession number GSE315827.

### 4.11 Code availability

All code to reproduce the analyses is publicly available from https://github.com/seidels/ `sciphy-materials`. The SciPhy codebase is openly accessible at https://github.com/ `azwaans/SciPhy`.

## Supporting information

sciphy-supplement

## 5 DECLARATION OF GENERATIVE AI AND AI-ASSISTED TECHNOLOGIES IN THE WRITING PROCESS

During the preparation of this work the authors used the ChatGPT language model by OpenAI to refine the manuscript’s language. After using this tool, the authors reviewed and edited the content as needed and take full responsibility for the content of the publication.

## 6 ACKNOWLEDGMENTS

This project has received funding from the European Research Council (ERC) under the European Union’s Horizon 2020 research and innovation programme grant agreement no. 101001077 (to T.S.) and the Paul G. Allen Family Foundation (to J.S.). The authors would like to thank François Bienvenu, Timothy G. Vaughan, Louis A. Du Plessis, Cecilia Valenzuela Agüi, Marcus Overwater, Nicola Mulberry, Laura Tomás and Ugnė Stolz for helpful discussions and feedback, as well as Léah Friedman and Thomas Gregor for providing their raw data on raw cell counts throughout gastruloid development.

## 7 AUTHORS’ CONTRIBUTIONS

S.S.: conceptualization, methodology, software, validation, formal analysis, investigation, data curation, writing—original draft, writing—review and editing, visualization, project administration. A.Z.: conceptualization, methodology, software, validation, formal analysis, investigation, data curation, writing—original draft, writing—review and editing, visualization S.R.: data curation, writing—review and editing. J.C.: data curation, writing—review and editing. J.S.: writing—review and editing, supervision, funding acquisition. T.S.: conceptualization, methodology, resources, writing—review and editing, supervision, funding acquisition.

## A APPENDIX

### A.1 Validation

**Table 6.**
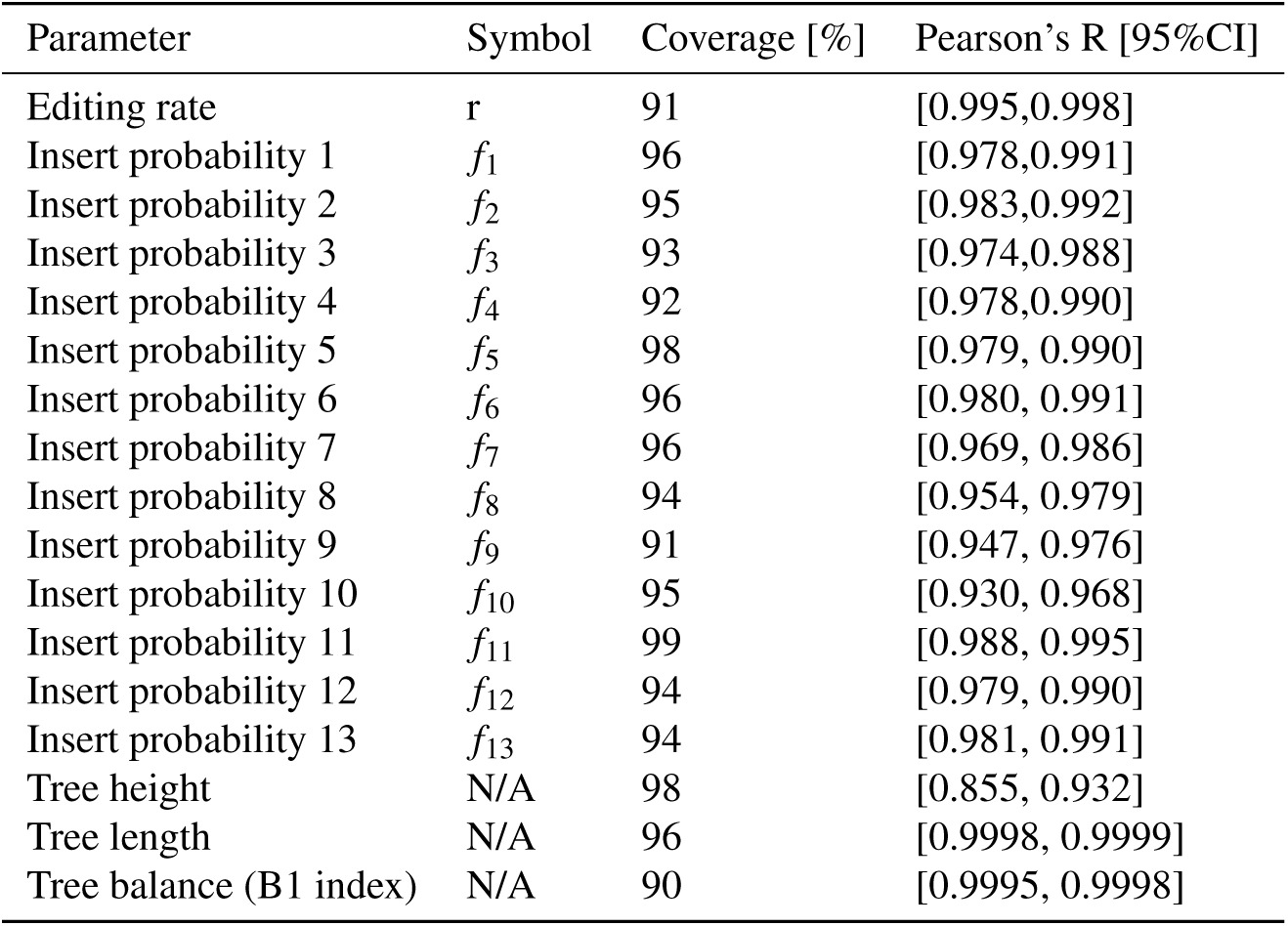
Coverages and correlations to true value in validation study.

**Figure 1.**
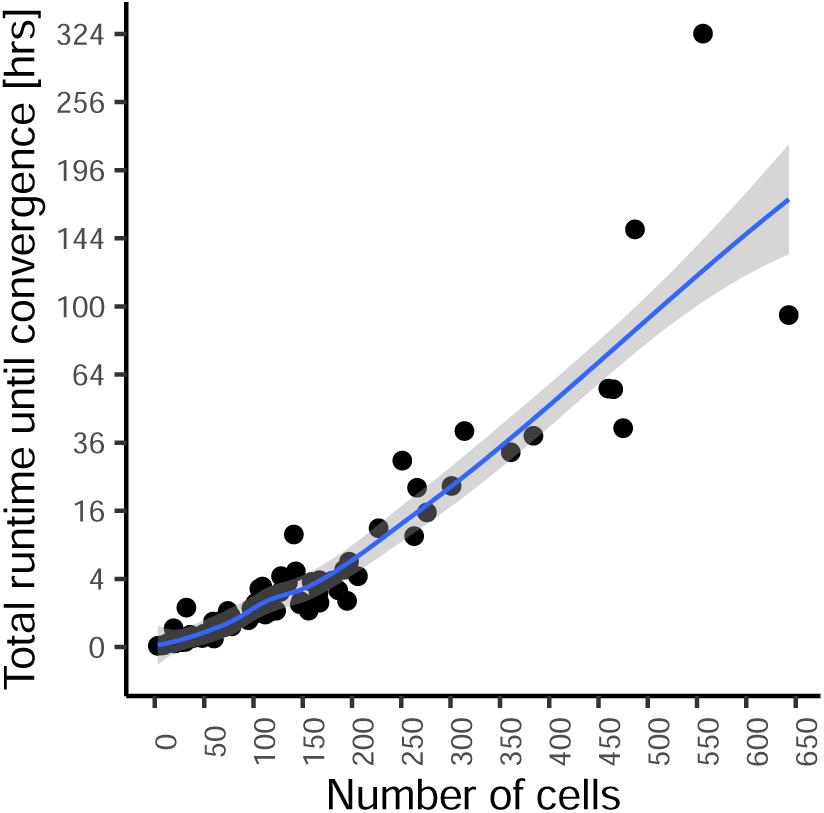
Approximate runtime needed for convergence of a SciPhy analysis. We report the runtime required for convergence of the lineage tree reconstruction and parameter estimation for all datasets in the validation study. We display here the empirical total runtime rescaled to reach a minimum ESS value of 200 for the SciPhy likelihood.

**Figure 2.**
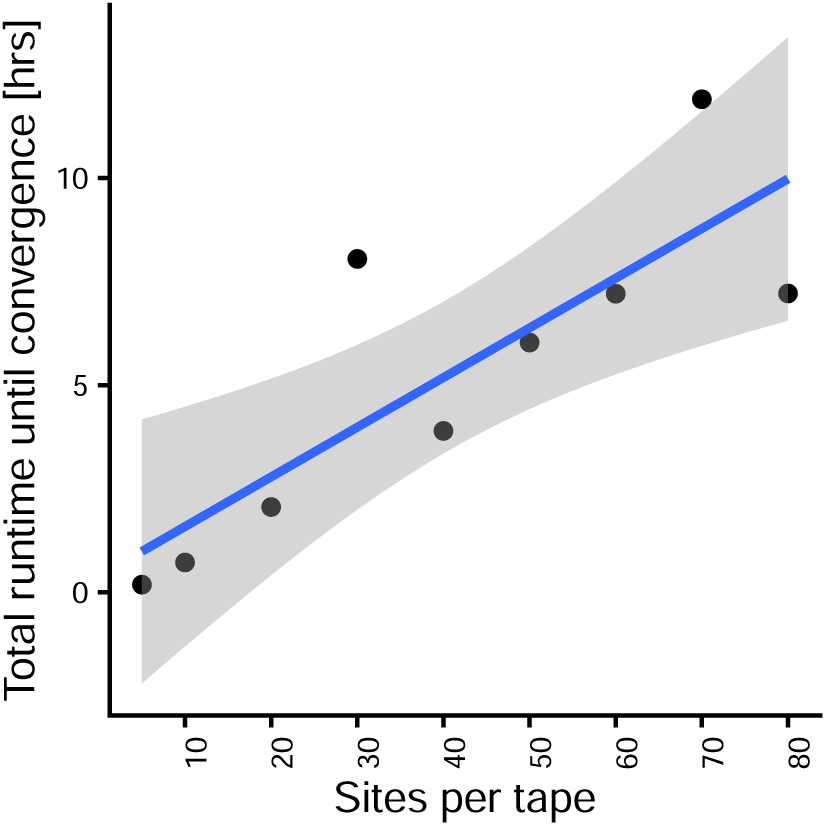
Approximate runtime needed for convergence of a SciPhy analysis w.r.t. tape length. We report the runtime required for convergence of the lineage tree reconstruction and parameter estimation based on 123 cells harboring a single tape, varying the number of sites per tape. We display here the empirical total runtime rescaled to reach a minimum ESS value of 200 for the SciPhy likelihood.

**Figure 3.**
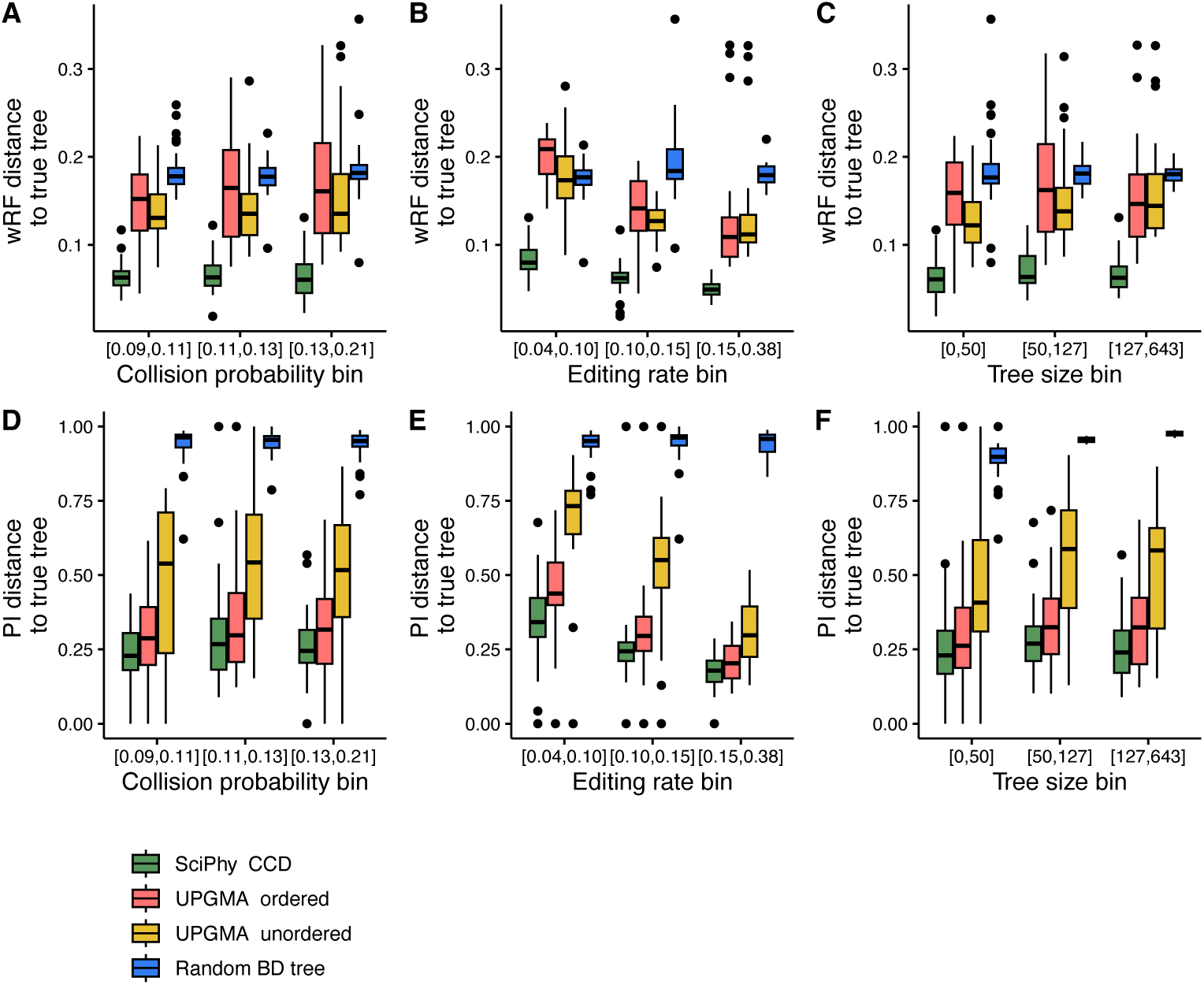
Accuracy of SciPhy reconstructed trees w.r.t. different simulated parameter regimes, benchmarked to UPGMA on all validation datasets. For all datasets simulated in the validation study, we showcase the distance from the true tree to trees reconstructed with SciPhy (summarised using the Conditional Clade Distributions, denoted as SciPhy CCD), an order-aware UPGMA method (UPGMA ordered), the standard UPGMA method, ignoring the order between edits (UPGMA unordered), and randomly generated through simulation under the birth-death sampling model (Random BD tree), where UPGMA trees are all scaled to 25 days for comparison. These distances are shown for binned simulation parameters values and calculated using the weighted Robinson Foulds (wRF, panels A-C) metric and Phylogenetic Information metric (PI, panels D-F). The “collision probability” presented here is the sum of squared insert probabilities, and represents how skewed towards certain inserts the editing process is.

**Figure 4.**
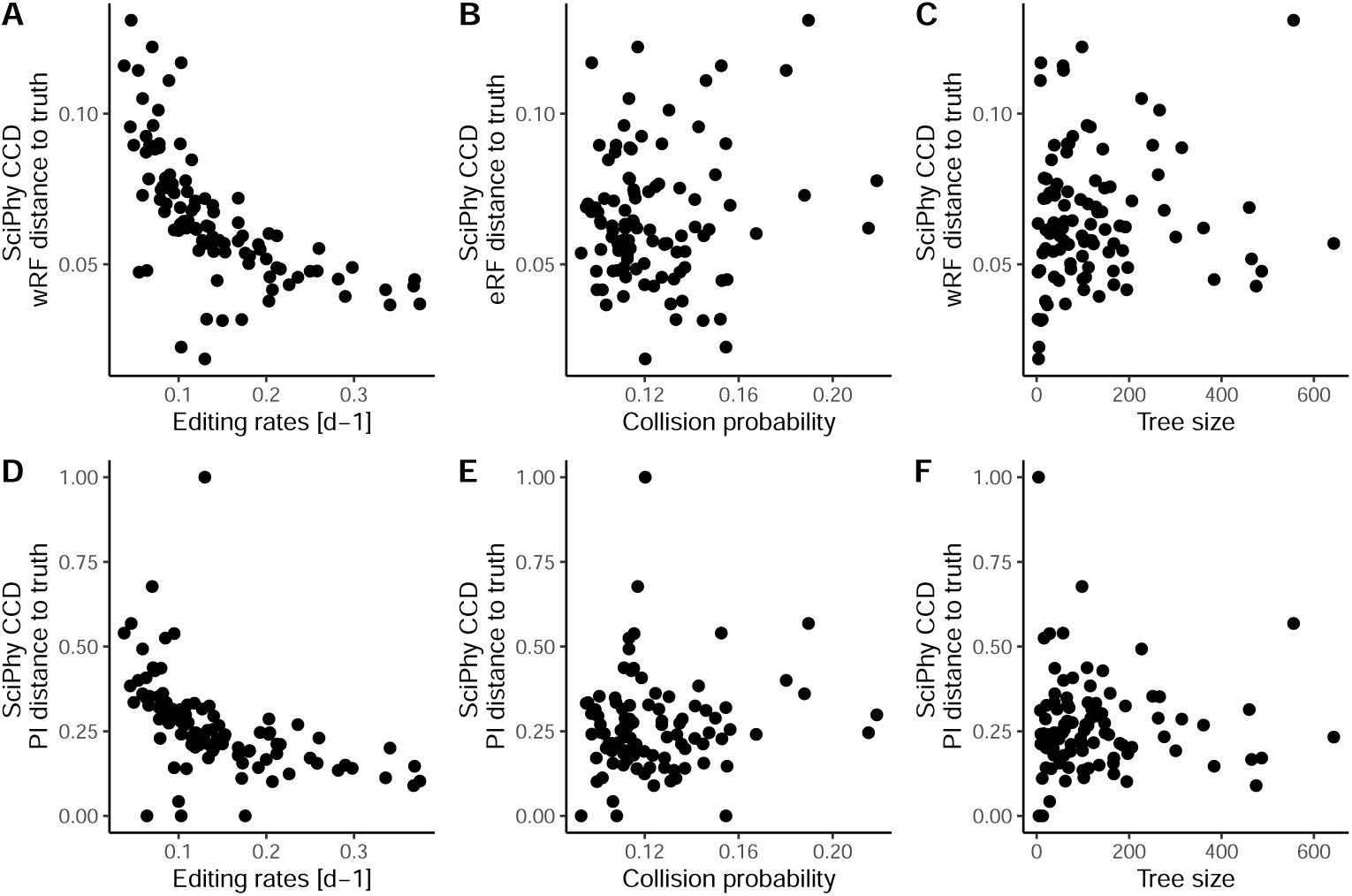
Accuracy of SciPhy reconstructed trees w.r.t. simulated parameter values on all validation datasets. For all datasets simulated in the validation study, we showcase the distance from the true tree to trees reconstructed with SciPhy (summarised using the Conditional Clade Distributions, SciPhy CCD). These distances are shown against simulation parameters values and calculated using the weighted Robinson Foulds (wRF, panels A-C) metric and Phylogenetic Information metric (PI, panels D-F). The “collision probability” presented here is the sum of squared insert probabilities, and represents how skewed towards certain inserts the editing process is.

### A.2 Cell culture

**Figure 5.**
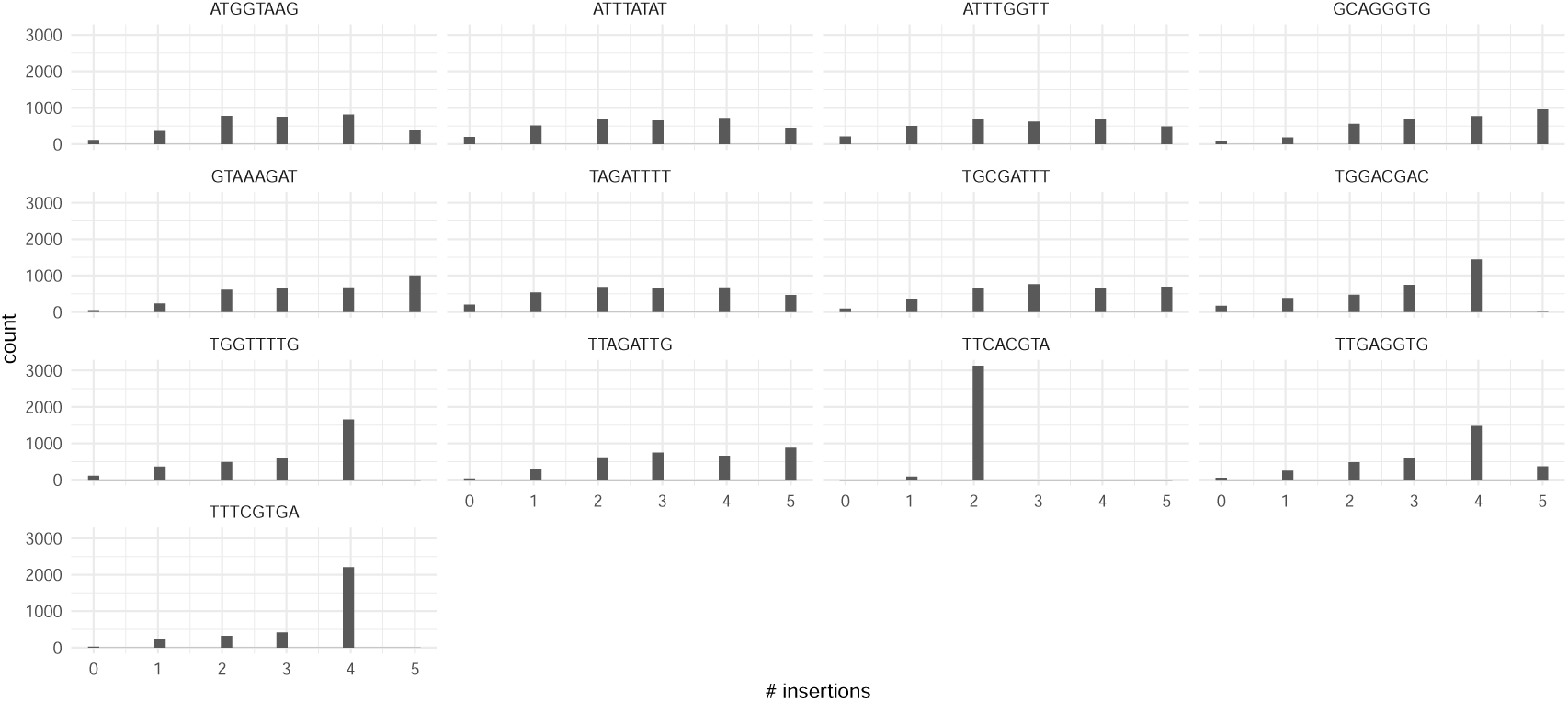
Number of insertions per tape in the HEK293T dataset. The majority of non truncated barcodes accumulate between 2 and 5 insertions throughout the experiment.

**Figure 6.**
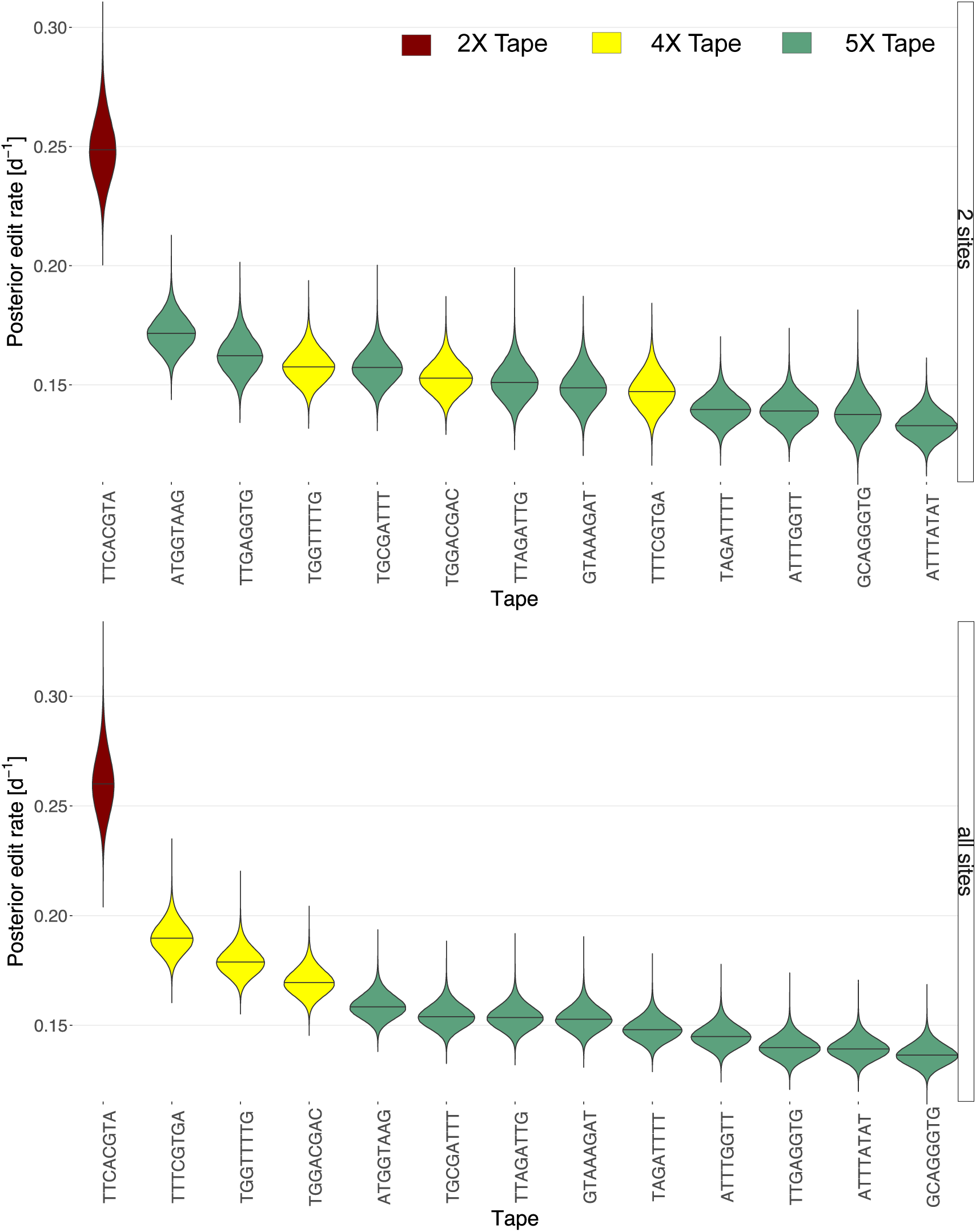
Comparing estimated editing rates. We show the estimated rate of editing for each tape. In the upper row, we show the results when estimating the editing rates from a dataset where all tapes are truncated to 2 sites. The lower row shows the analysis from the main text Fig. 3 where we include all available sites per tape. The edit rates are ordered by their median and are colored according to original tape length, where 2X means 2 sites etc.

**Figure 7.**
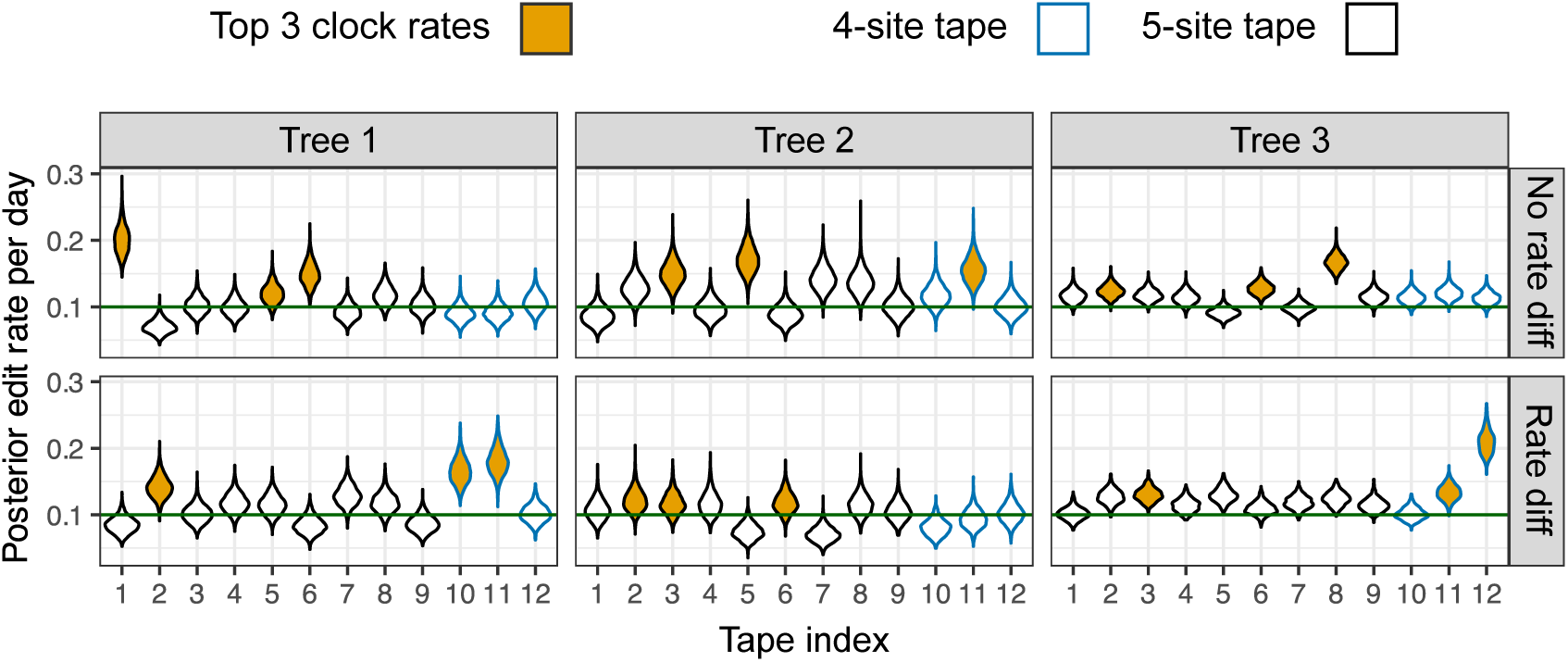
Effect of within-tape rate variation on inferred editing rates. Posterior estimates of edit rates per tape under SciPhy for three simulated trees. In each panel, tapes are shown along the x-axis (tape indices 1–12), with violin plots representing the posterior distribution of the per-tape edit rate. Tapes 10–12 (blue outlines) are 4-site tapes; others are 5-site tapes. Orange-filled violins indicate the three tapes with the highest median inferred edit rates. Top row: scenario with no rate differences between sites. Bottom row: scenario where the 5th site is edited at 20% of the base rate. In the “Rate diff” condition, 4-site tapes more frequently appear among the fastest edited rates, as they lack the slower 5th site.

**Figure 8.**
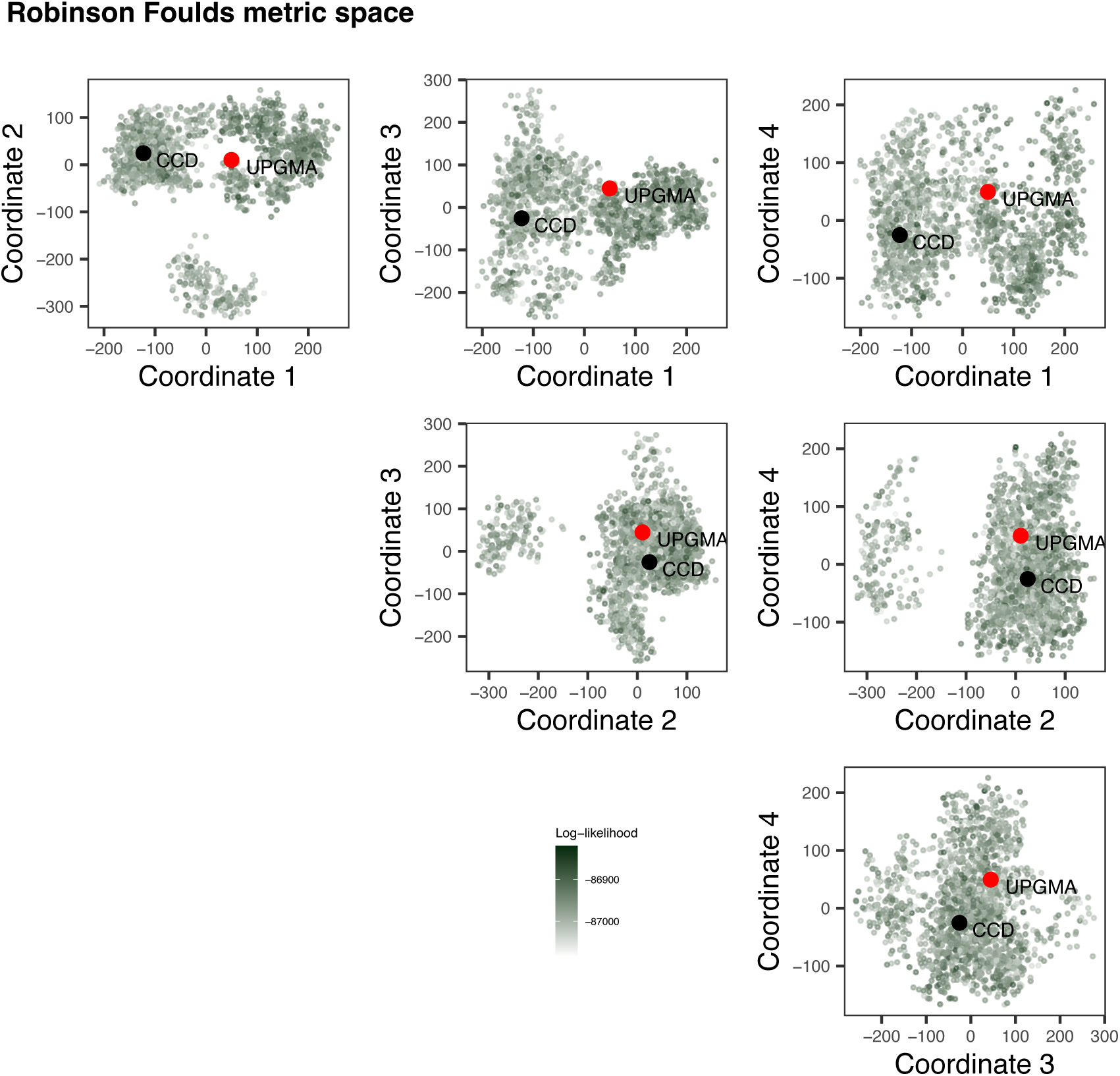
Comparison of the SciPhy posterior tree set to the UPGMA tree obtained from the HEK293T data, with respect to topology. Pairwise Robinson Fould (RF) distances between SciPhy posterior trees, the UPGMA tree and the CCD tree estimated for the HEK293T dataset are mapped in 4 coordinates pairwise to visualize these tree sets.

**Figure 9.**
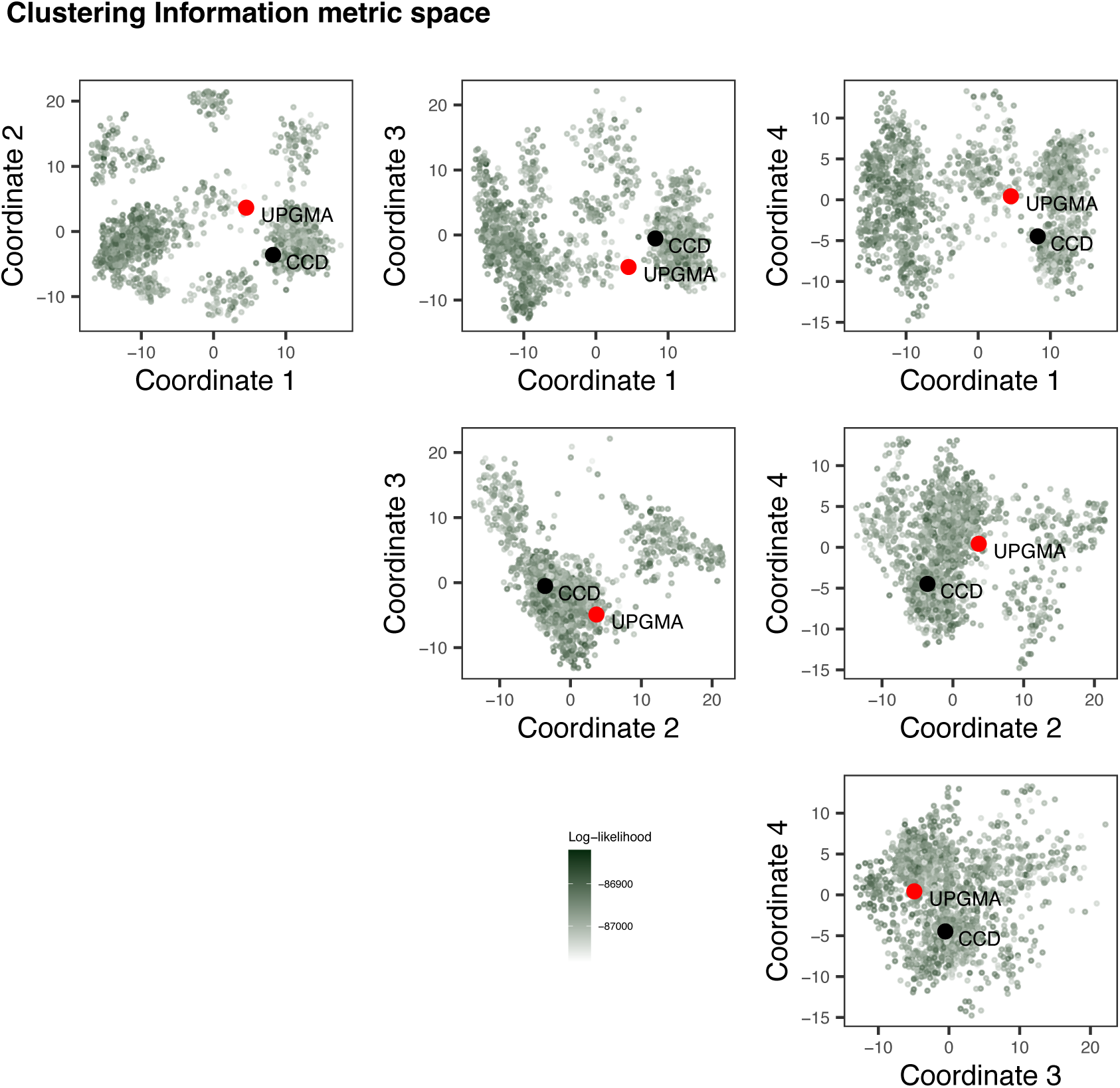
Comparison of the SciPhy posterior tree set to the UPGMA tree obtained from the HEK293T data, with respect to topology. Pairwise Clustering Information (CI) distances between SciPhy posterior trees, the UPGMA tree and the CCD tree estimated for the HEK293T dataset are mapped in 4 coordinates pairwise to visualize these tree sets.

**Figure 10.**
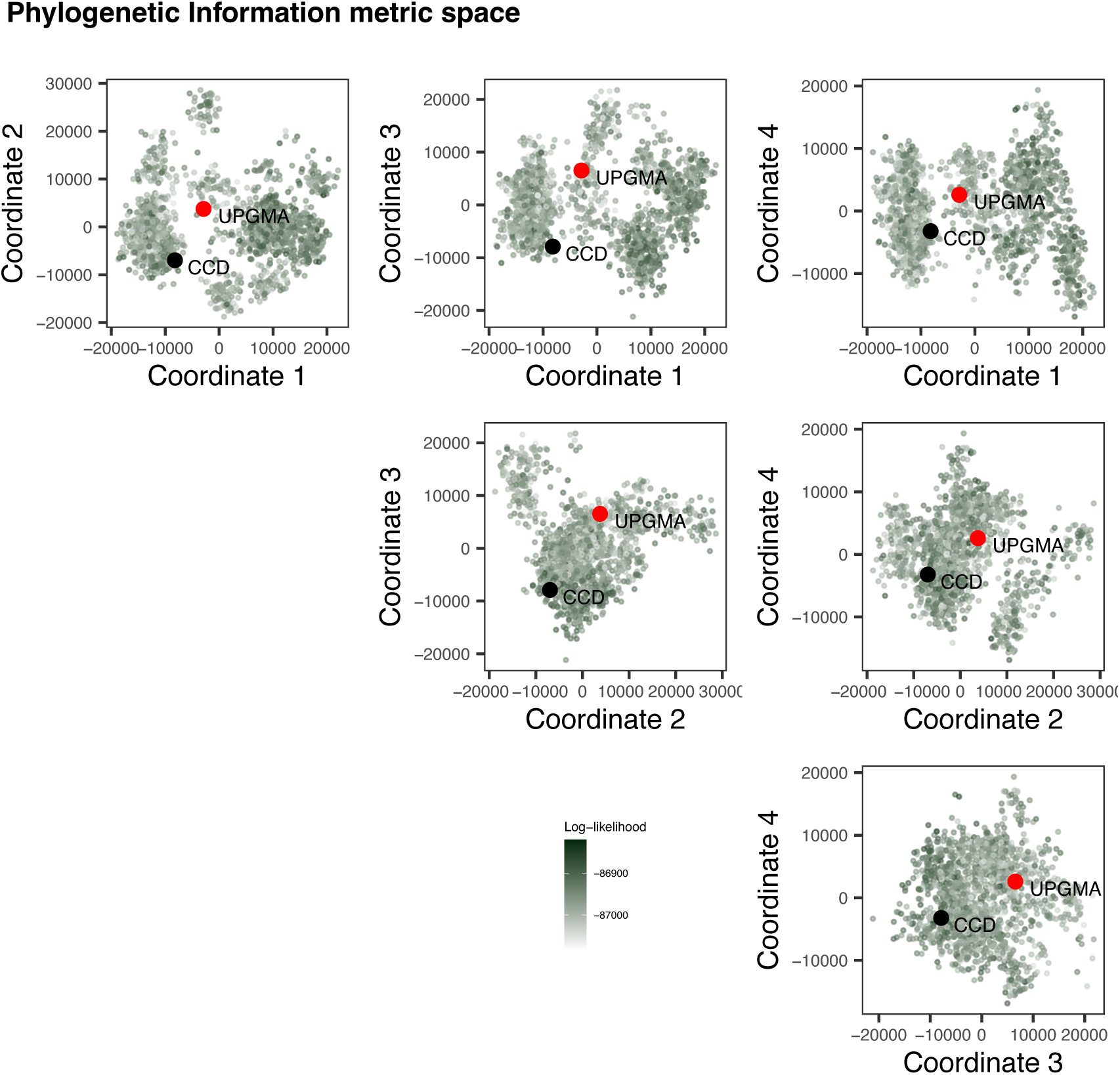
Comparison of the SciPhy posterior tree set to the UPGMA tree obtained from the HEK293T data, with respect to topology. Pairwise Phylogenetic Information (PI) distances between SciPhy posterior trees, the UPGMA tree and the CCD tree estimated for the HEK293T dataset are mapped in 4 coordinates pairwise to visualize these tree sets.

**Figure 11.**
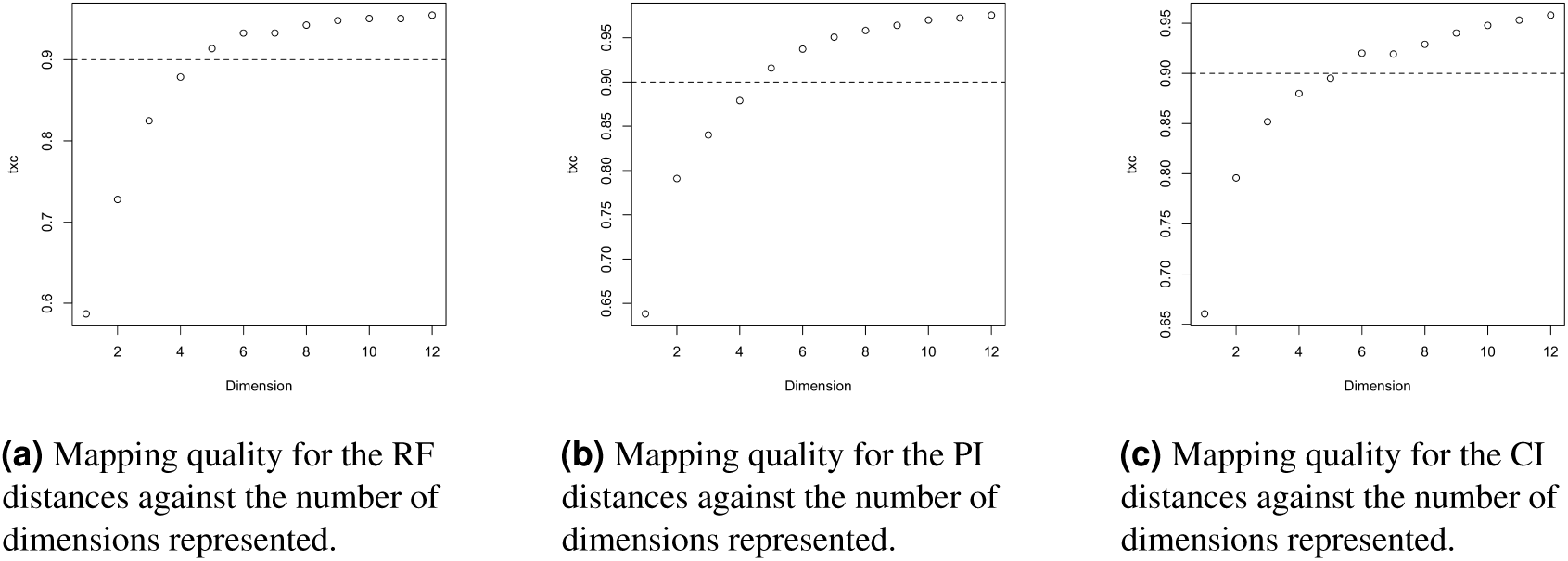
2D mapping quality per number of dimensions plotted for the RF, PI and CI distances. For each of the 2D mappings of pairwise topological distances presented in our study, we plot here the trustworthiness × continuity score (txc) against the number of dimensions from the mapping used the represent the sets of trees shown. A mapping is considered sufficiently representative for a txc score of about 0.9.

**Figure 12.**
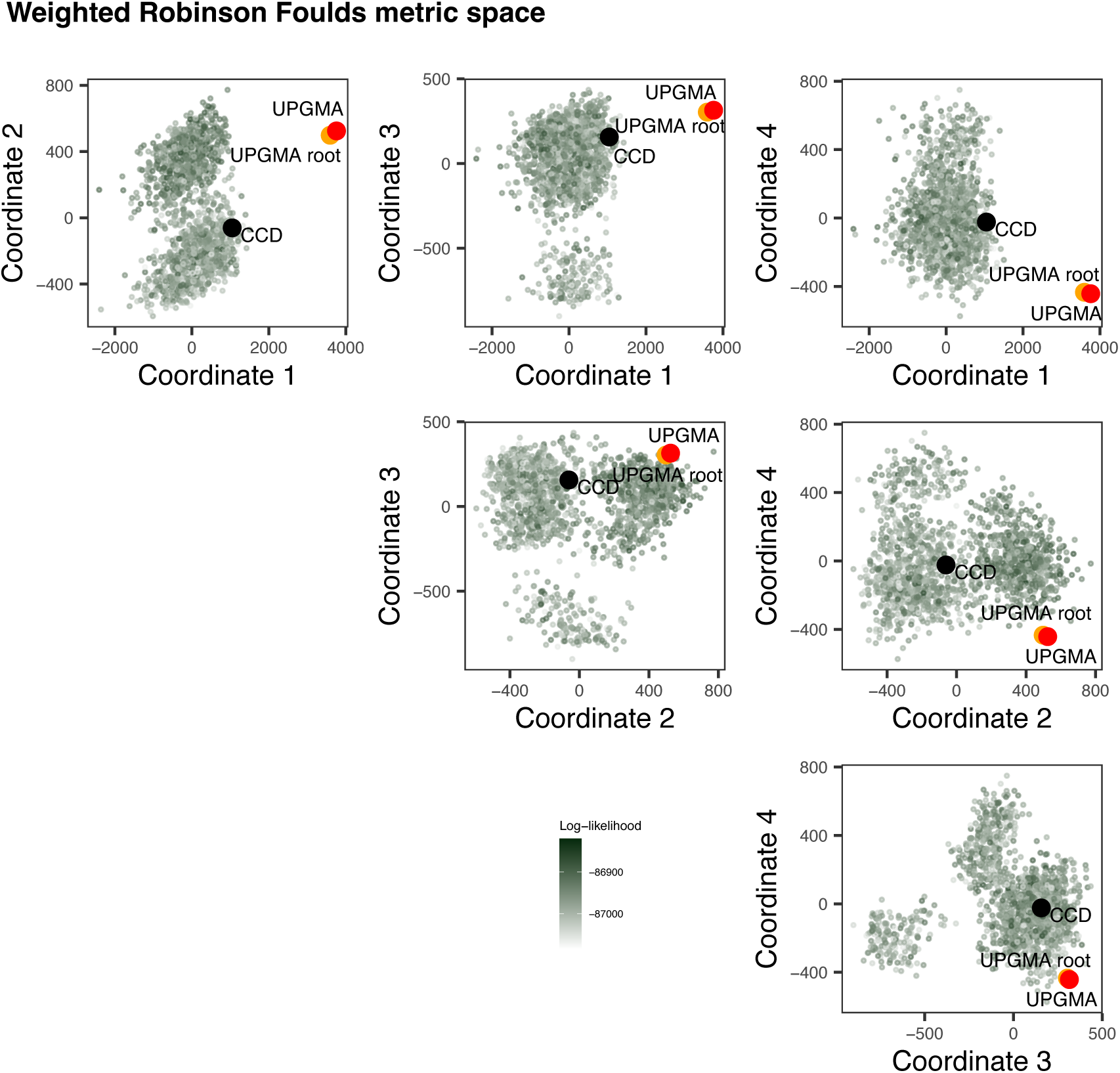
Comparison of the SciPhy posterior tree set to the UPGMA tree obtained from the HEK293T data, with respect to topology and branch lengths. Pairwise weighted Robinson Fould (wRF)distances between SciPhy posterior trees, the UPGMA tree (labelled ‘UPGMA’), the UPGMA tree scaled to the median estimated tree height estimated by SciPhy (labelled ‘UPGMA root’) and the CCD tree estimated for the HEK293T dataset are mapped in 4 coordinates pairwise to visualize these tree sets.

**Figure 13.**
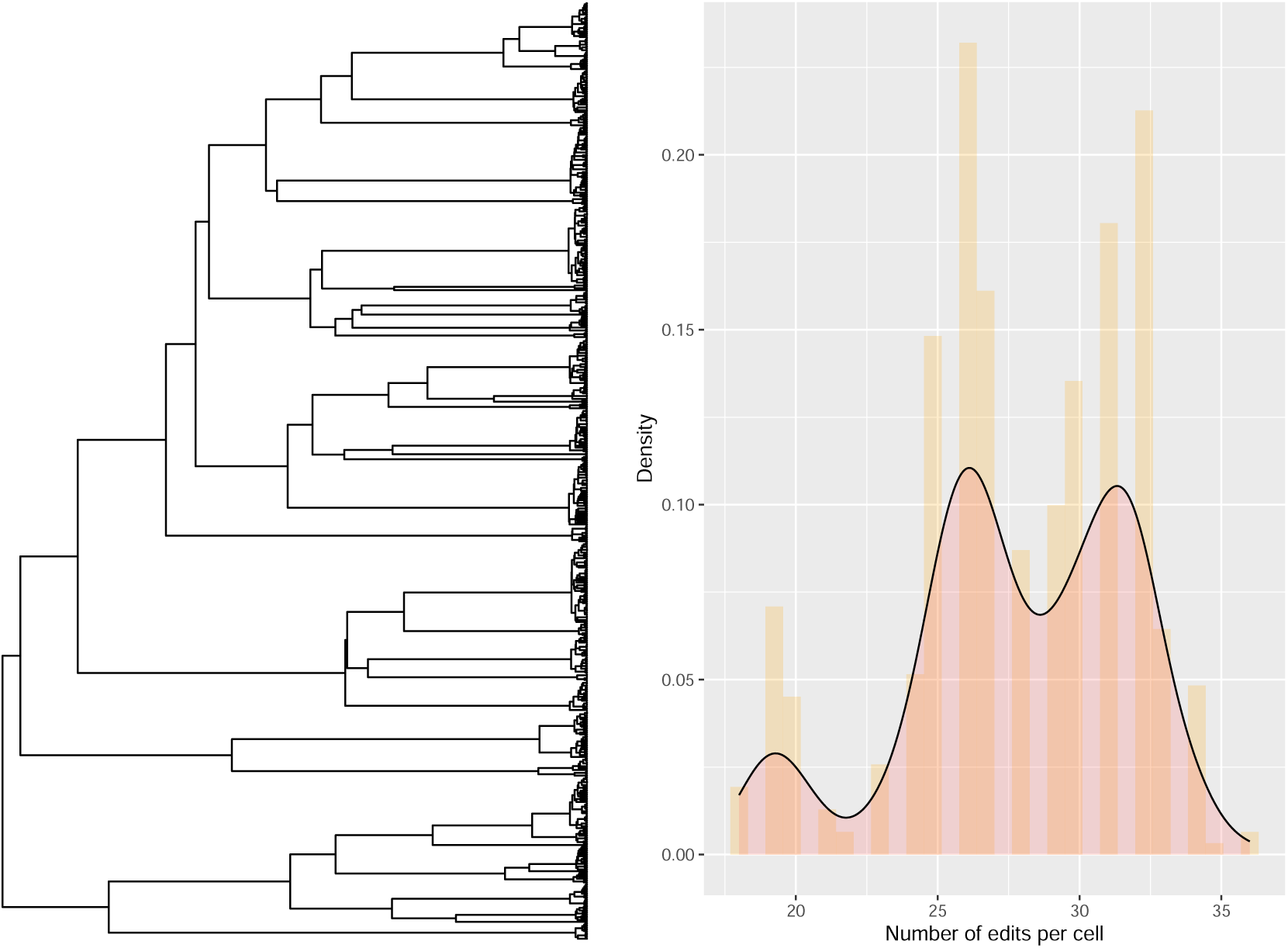
Simulated distribution of the total number of edits (right) under SciPhy along a tree (left) with early divisions. Here, SciPhy data (13 tapes of length 5) was simulated along a tree where the majority of branching events happen toward the present. This corresponds to a scenario where the numbers of edits per cell are not independent for the majority of the history of the cell population, and where the structure of the lineage tree leaves an imprint on the resulting distribution of the total edits per cell at the tips.

**Figure 14.**
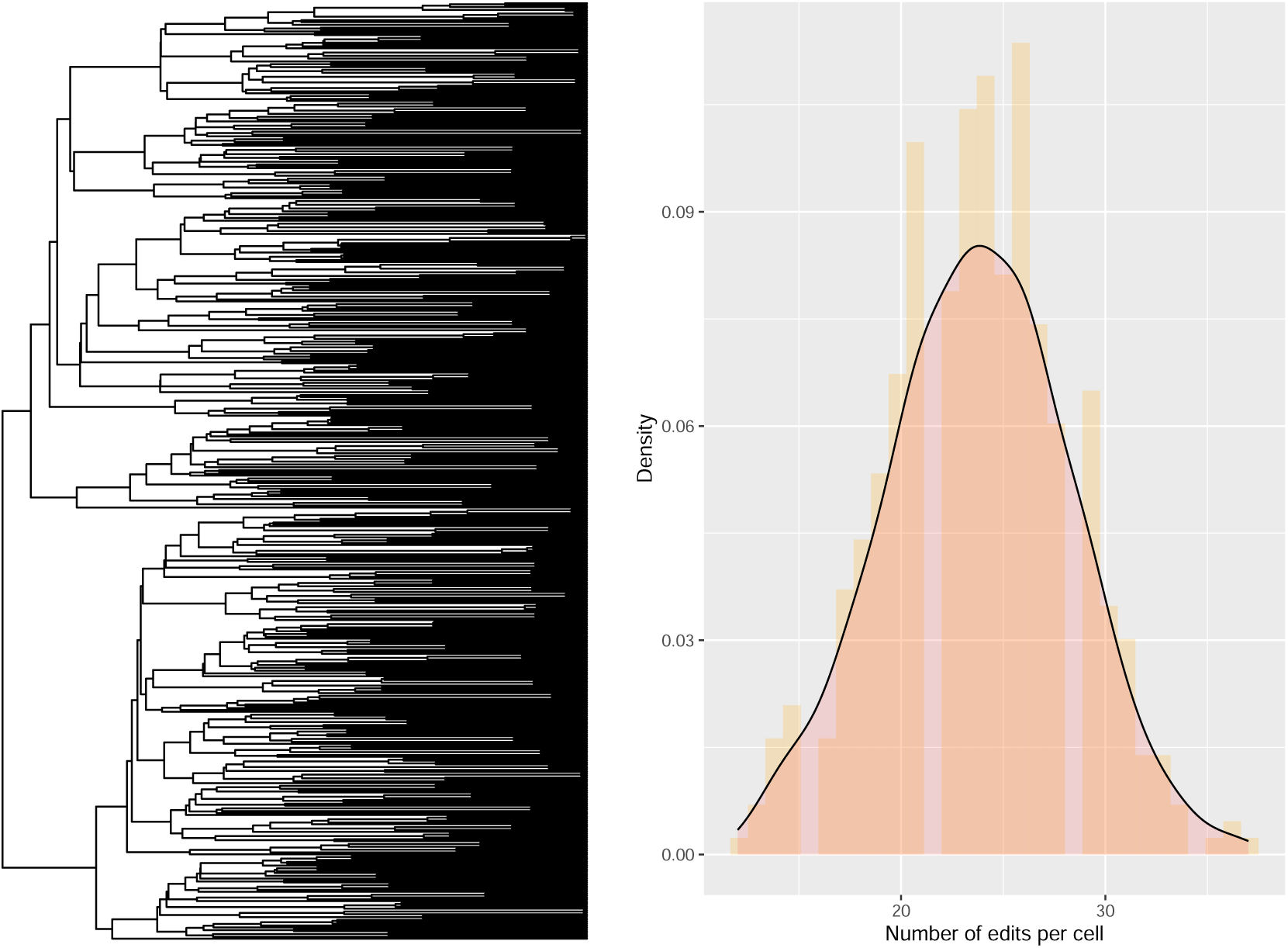
Simulated distribution of the total number of edits (right) under SciPhy along a tree (left) with late divisions. Here, SciPhy data (13 tapes of length 5) was simulated along a tree where the majority of branching events happen toward the past. This corresponds to a scenario where the numbers of edits per cell are independent for the majority of the history of the cell population, resulting in a unimodal distribution of the total number of edits.

**Figure 15.**
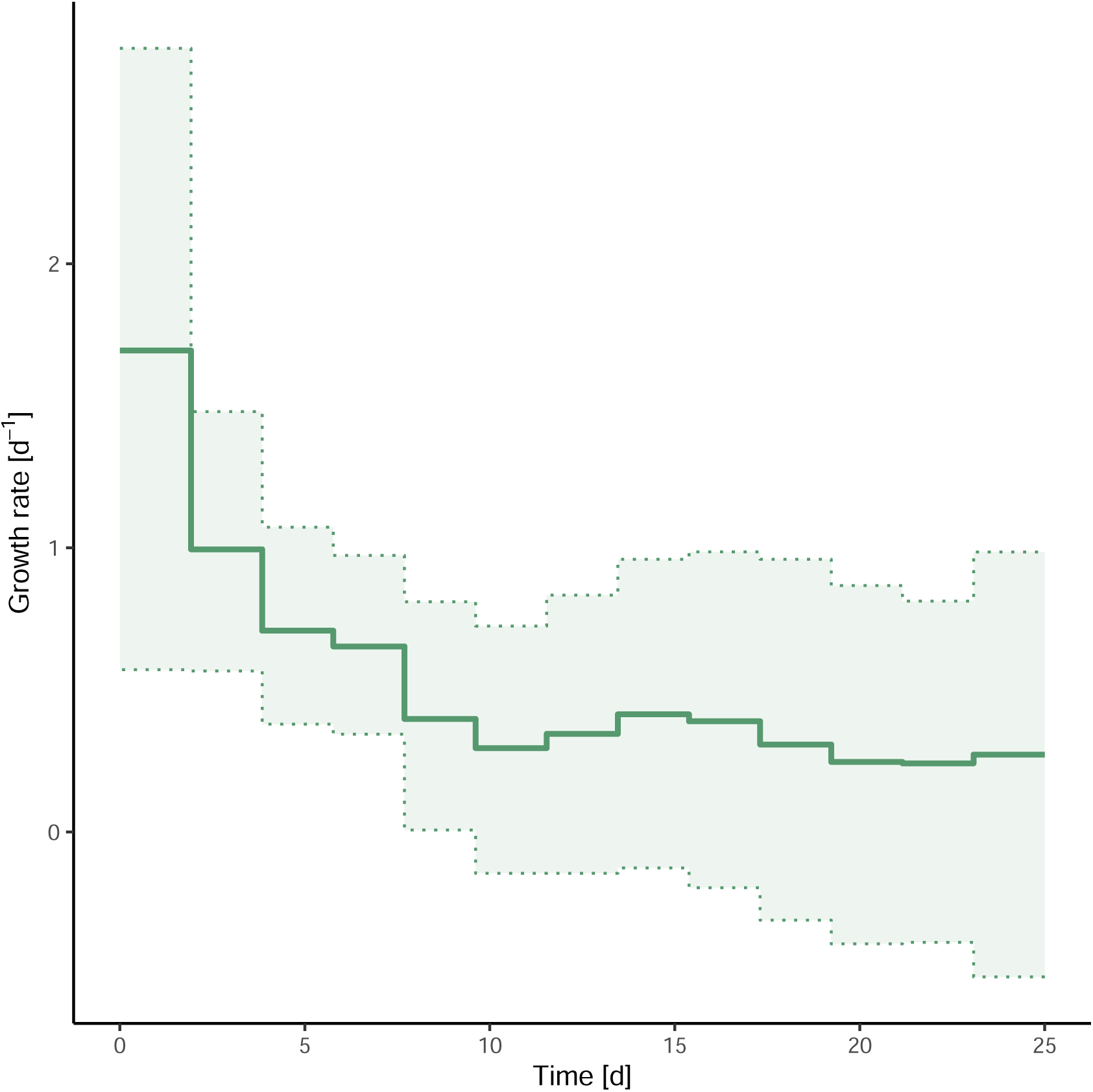
Time-varying (dynamic) growth rate estimated for the HEK293T cell population, with a sampling proportion fixed to 0.0008. We report the time-varying growth rate estimated using SciPhy, where the growth rate is allowed to vary every 2 days over the duration of the experiment, using an Ornstein-Ulhenbeck (OU) smoothing prior.

**Figure 16.**
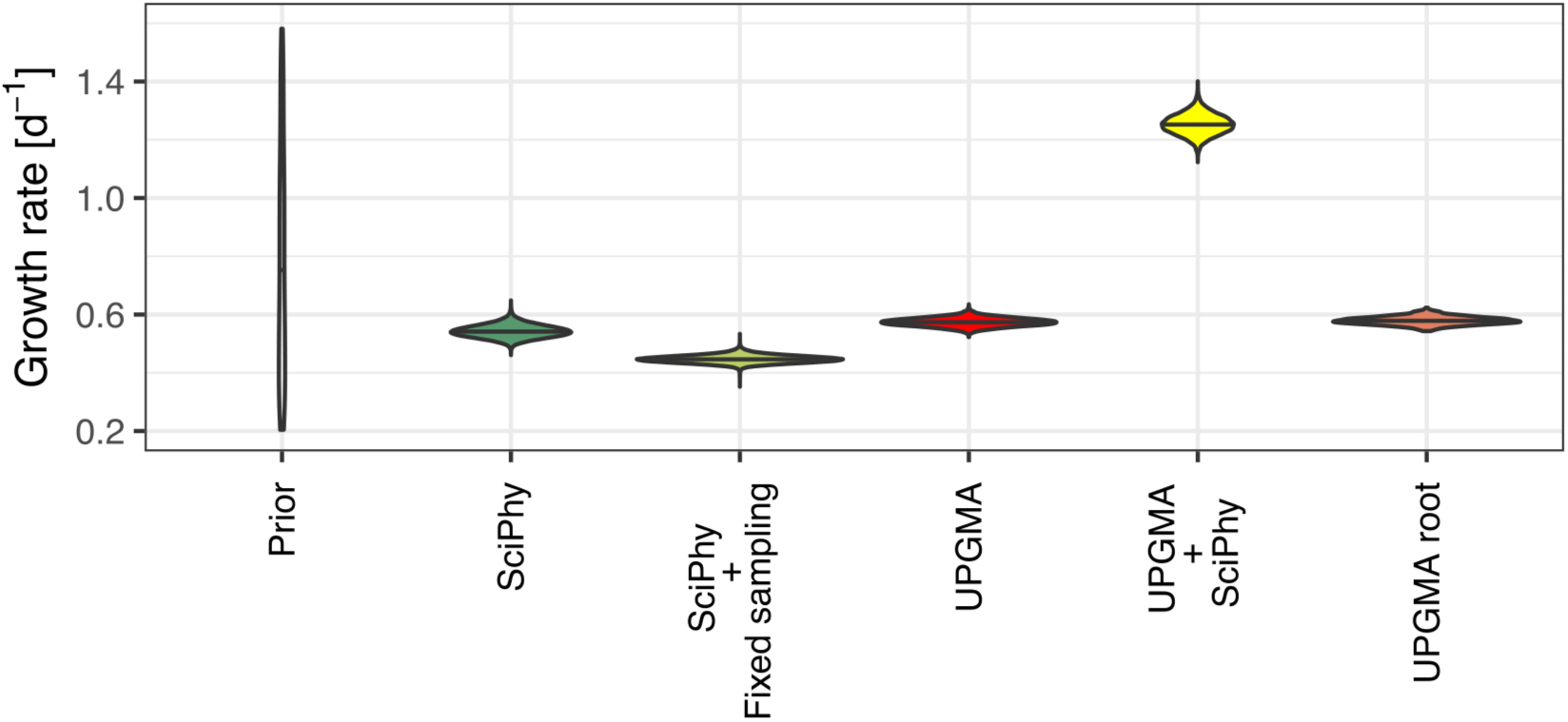
Comparing growth rates estimated with SciPhy and on fixed phylogenies. We report the constant growth rate estimated using SciPhy, with the sampling proportion fixed to 0.0008 (”SciPhy + Fixed sampling”), with a co-estimated sampling proportion (”SciPhy”), contrasted to inferences based on a fixed UPGMA tree topology. For comparison, we look at the UPGMA tree topology with the root height scaled to 25 days (”UPGMA”), with its root scaled to the median SciPhy tree MRCAs and the UPGMA tree with branch lengths scaled using SciPhy (”UPGMA - SciPhy”).

#### Sampling from prior

To investigate whether the growth rate is informed mainly by the molecular sequence data (the tapes in our study), or the number of sequenced cells at the end of the experiment and the prior distributions, we re-ran the inference devoid of the sequence data (Appendix Fig. 17). This approach yielded a median growth rate of 0.57 per day (95% HPD interval: [0.48, 0.68] d−1), notably different from the estimate with sequence data and aligning with the expected growth rate under a deterministic exponential growth model. This suggests that the slower cell division rate we initially estimated is largely influenced by the sequence data.

**Figure 17.**
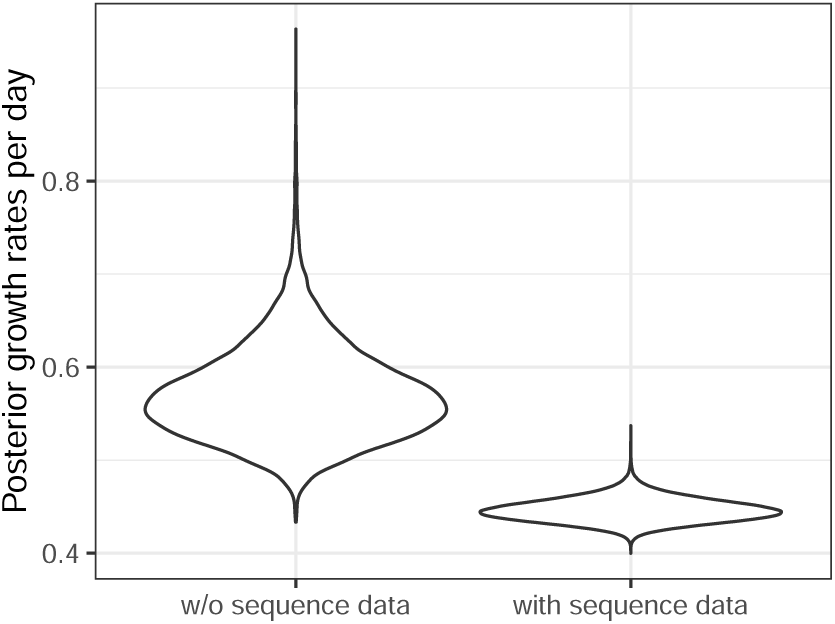
Comparing growth rate estimates with and without sequence data in the HEK293T analysis. We report the growth rates estimated using SciPhy as in the main text (with sequence data, right) and compare it to the same analysis just without the sequence data (left).

**Figure 18.**
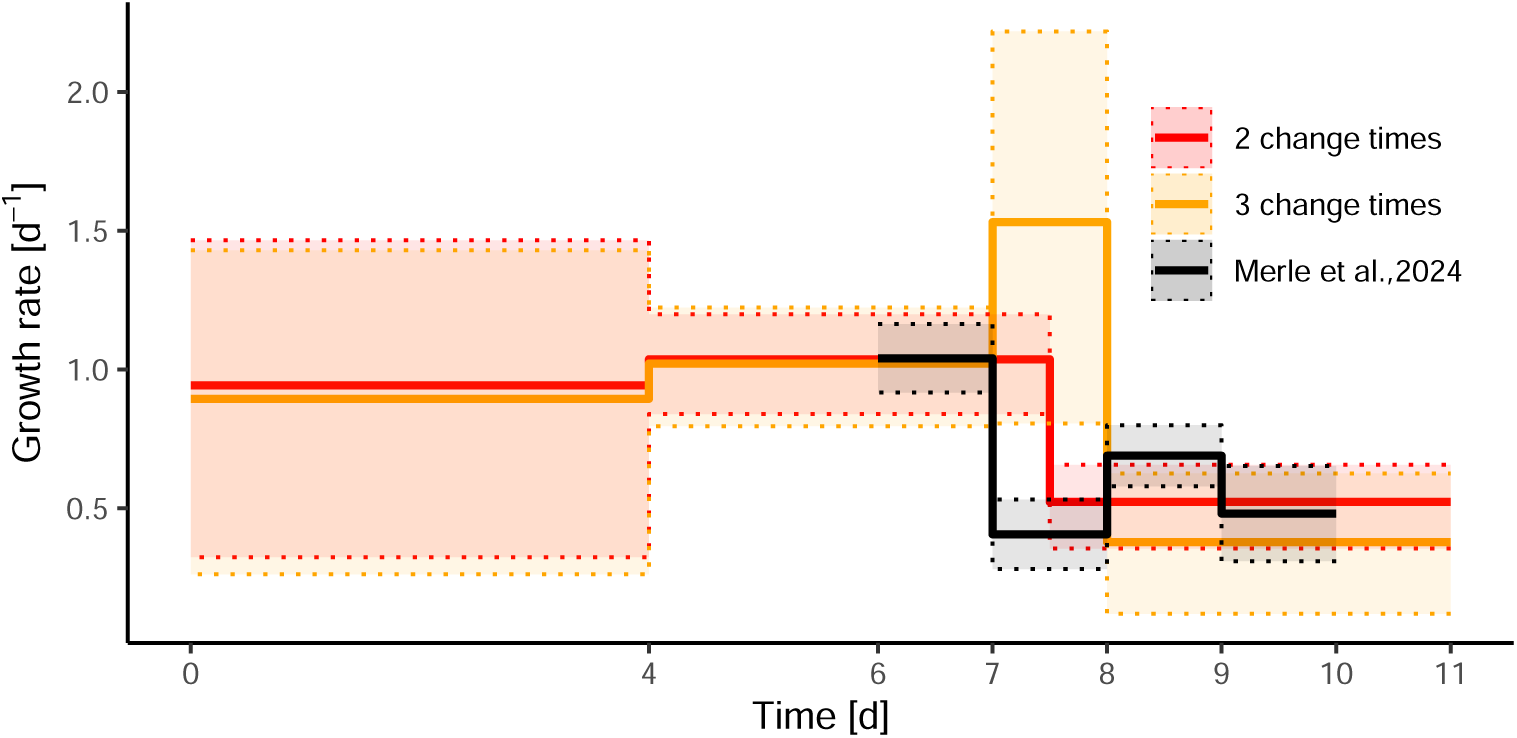
Comparing estimates of the time-varying growth rate for gastruloid data under different time divisions of the experimental timeline and against observed dynamics. We report the growth rates (median and 95% HPD) estimated using SciPhy under 2 change times (also in main text), and (3 change times) and the growth rates (mean ±*σ*) reported in Merle et al. (2024) for comparison.

**Figure 19.**
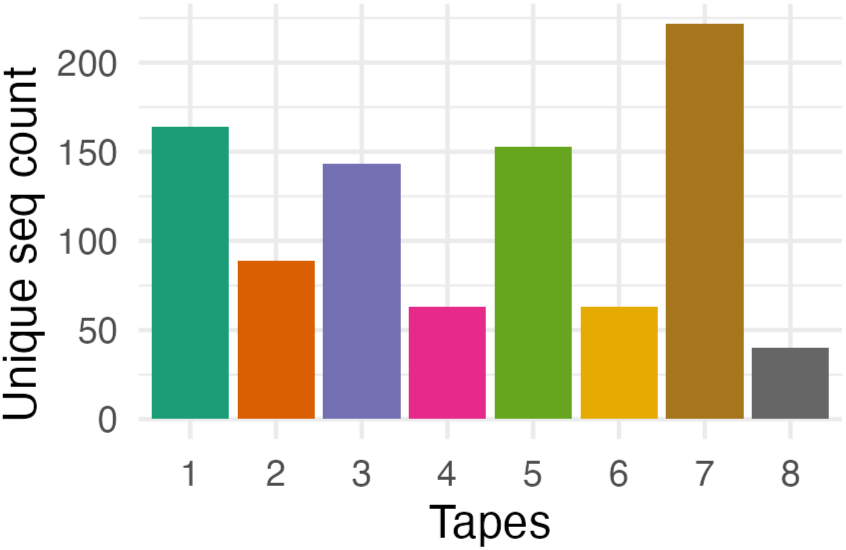
We show the number of unique insert sequences for each tape.

**Figure 20.**
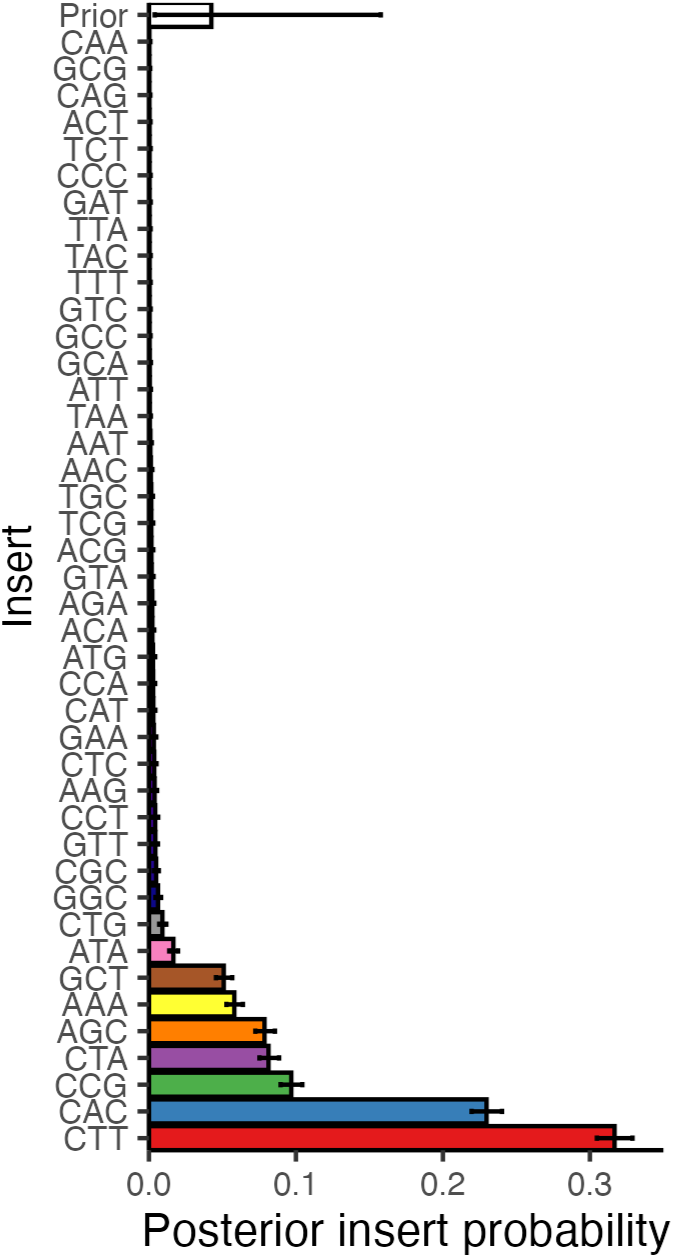
We show the posterior insertion probability for every possible insert. Note that we use a highly distinguishable color scheme for the first 9 inserts and a less distinguishable color scheme for the remaining 33 inserts, that have very low insertion probability.

**Figure 21.**
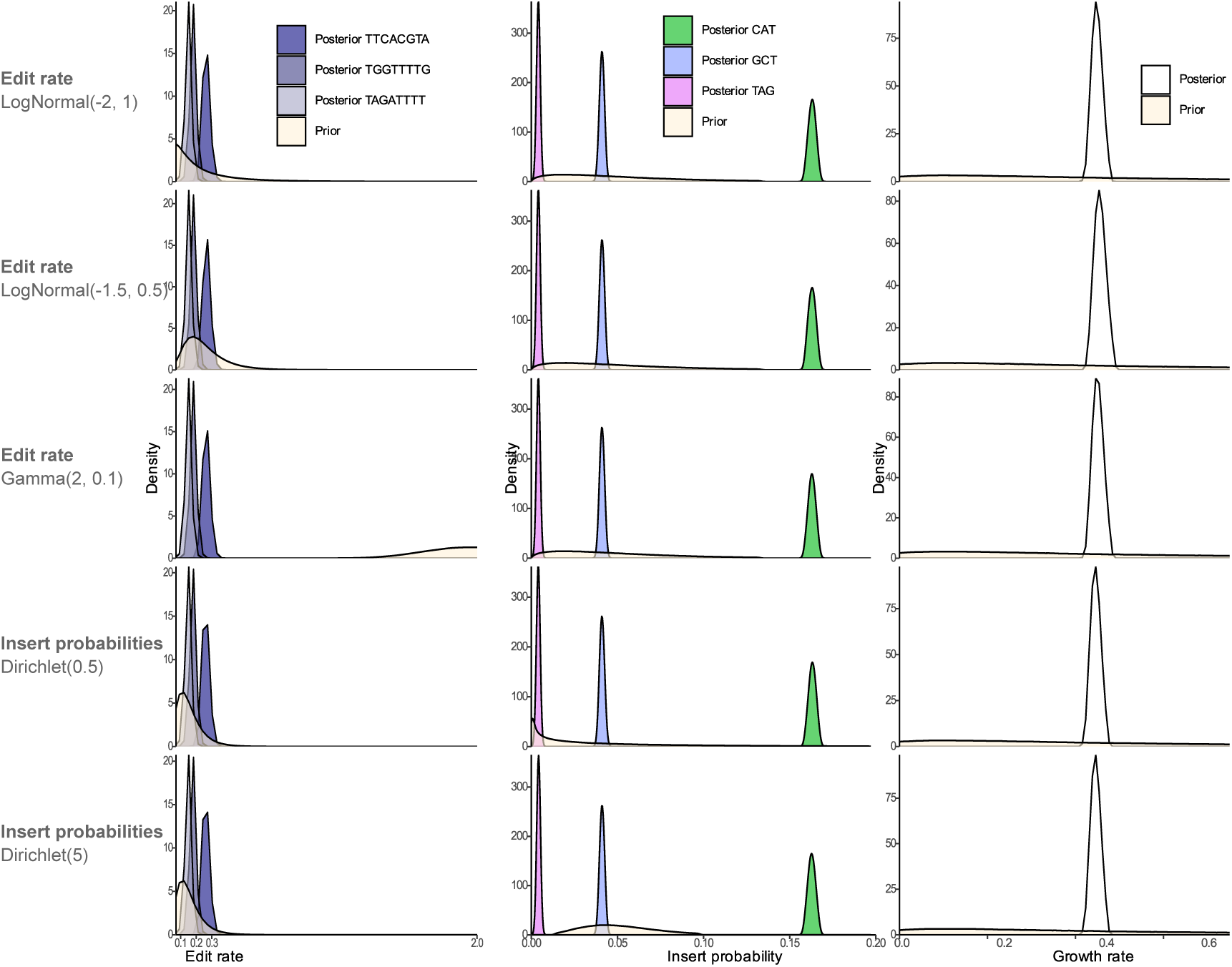
Results of the phylodynamic analysis of cell culture growth are robust to different choices of prior distributions.

**Figure 22.**
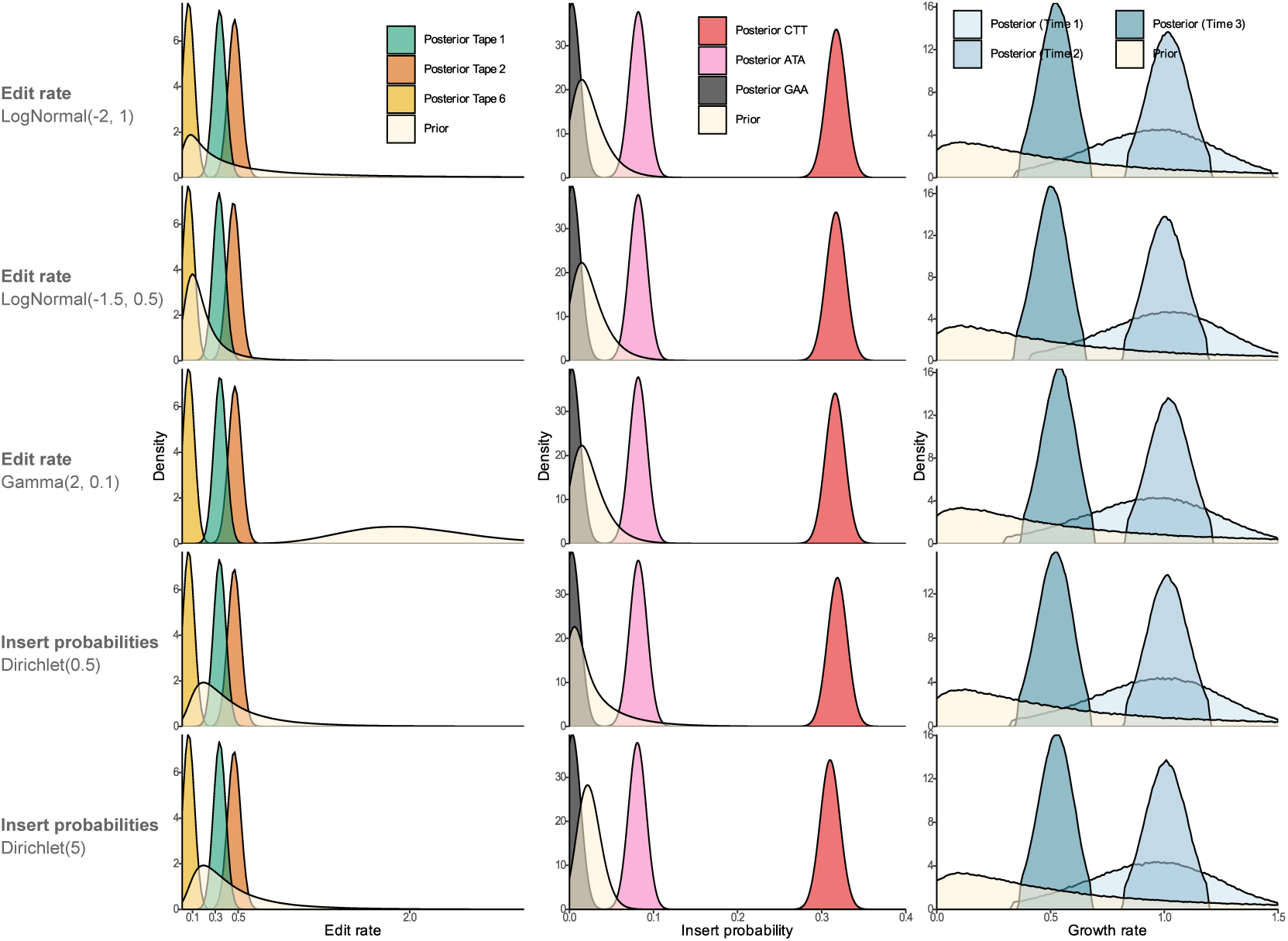
Results of the phylodynamic analysis of gastruloid growth are robust to different choices of prior distributions.

#### A.2.2 Simulation of incomplete tape alignments

**Table 7.**
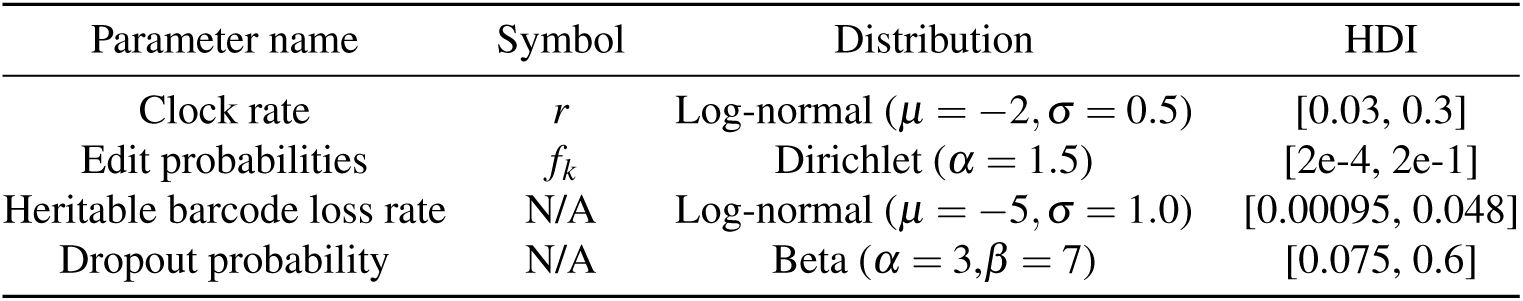
Distributions for the editing model parameters used to simulate incomplete barcode alignments.

**Figure 23.**
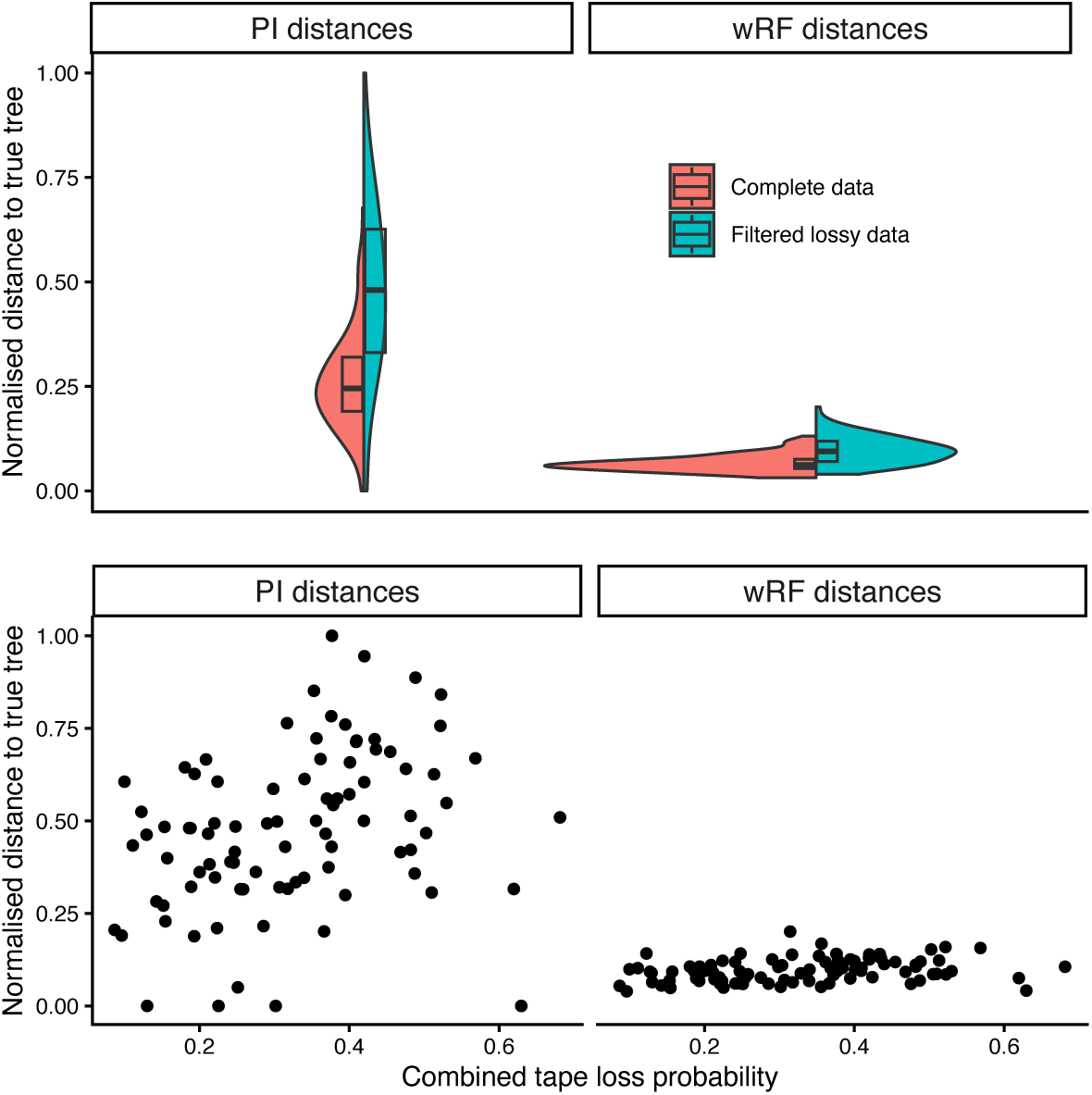
Robustness of tree inference using SciPhy to sparsity in tape alignments. In the top panel, for all trees simulated in the validation study, we showcase the distance from the true tree to trees reconstructed with SciPhy (summarized as point estimates with the conditional clade distribution (CCD) algorithm), using complete alignments of 10 tapes (”Complete data”) or lossy alignments (”Filtered lossy data”) of 20 tapes (leading to an average of 11 tapes after filtering) as input. In the bottom panel, we additionally show these distances against the combined tape loss (resulting of both heritable probability used for simulation of lossy alignments.

**Figure 24.**
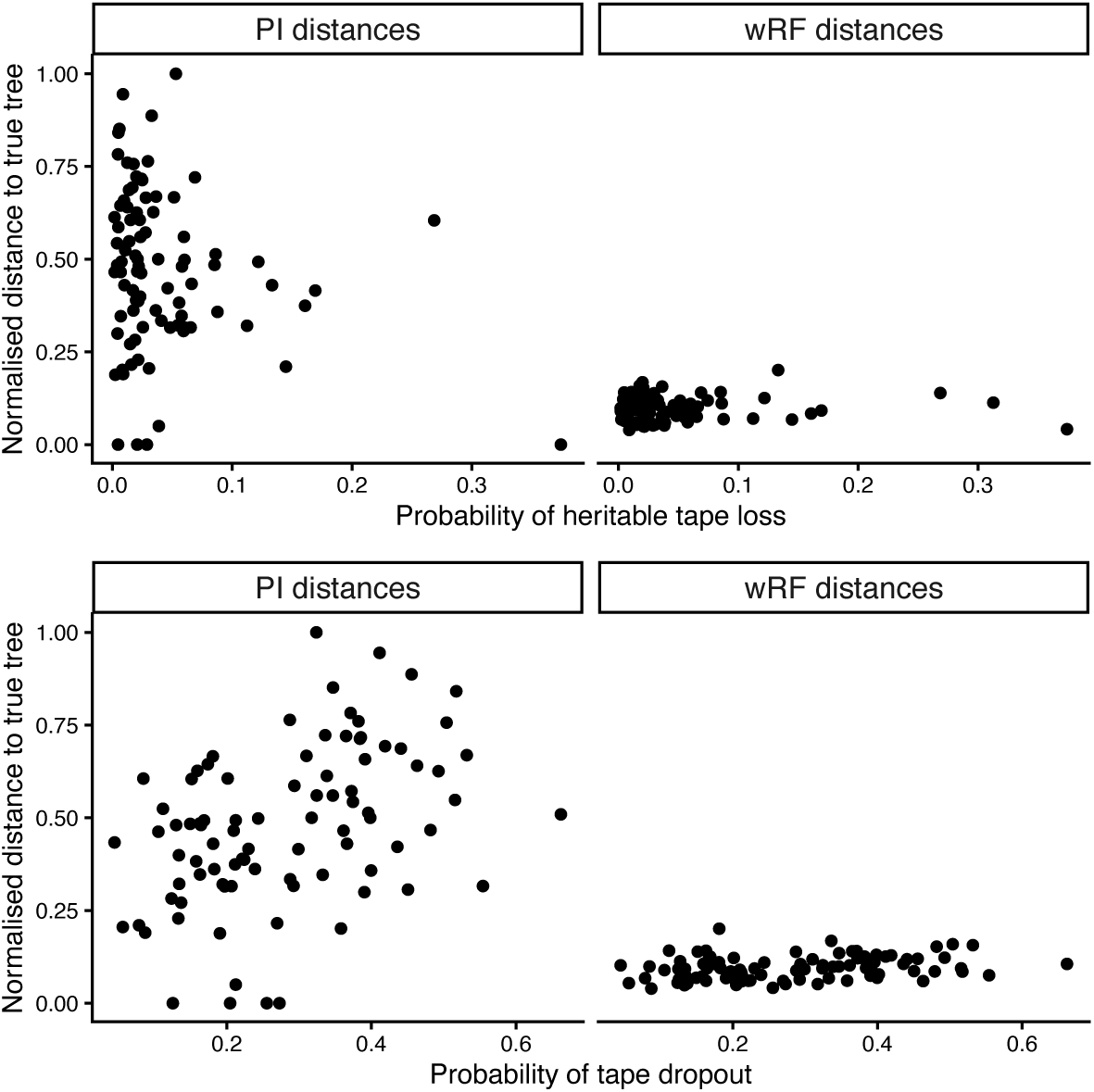
Robustness of tree inference using SciPhy to sparsity in tape alignments. For all trees simulated in the validation study, we showcase the distance from the true tree to trees reconstructed with SciPhy (summarized as point estimates with the CCD algorithm) using lossy alignments (”Filtered lossy data”) of 20 tapes (leading to an average of 11 tapes after filtering) as input. In the top panel, we show these distances against the heritable tape loss probability (or transgene silencing probability) used for simulation, while in the bottom panel we show these distances against the tape dropout probabilities (or probability of loss upon sampling).

**Figure 25.**
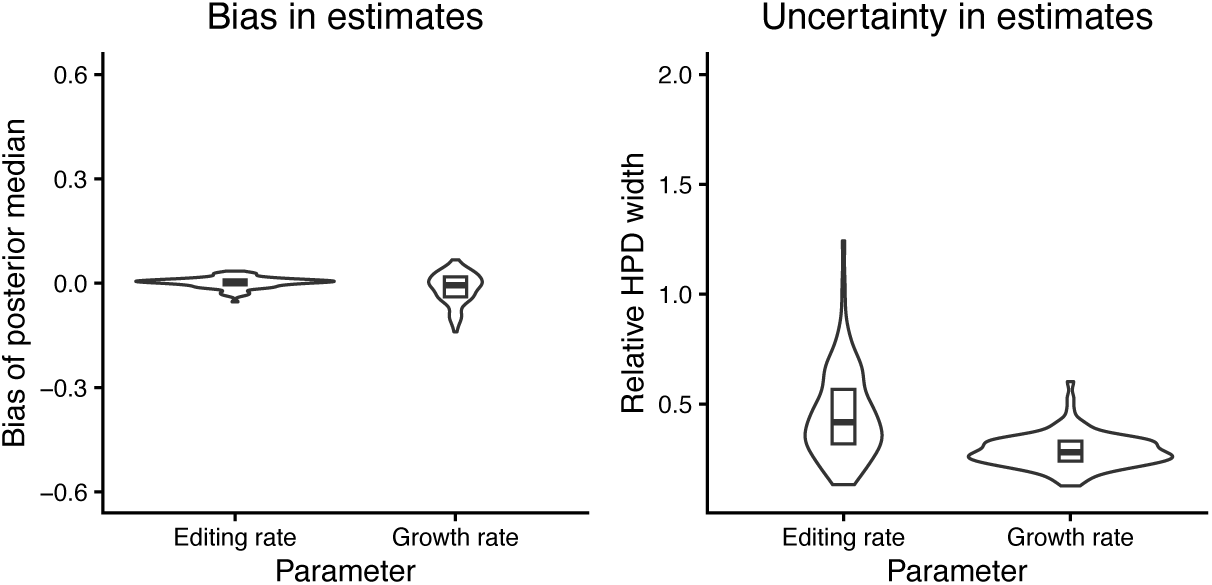
Robustness of parameter inference using SciPhy to sparsity in tape alignments. For all datasets simulated in the validation study, we showcase the bias (left) and uncertainty (right) in editing and growth estimates obtained using incomplete alignments (”Filtered lossy data” in Figure 23) of 20 tapes as input.

**Figure 26.**
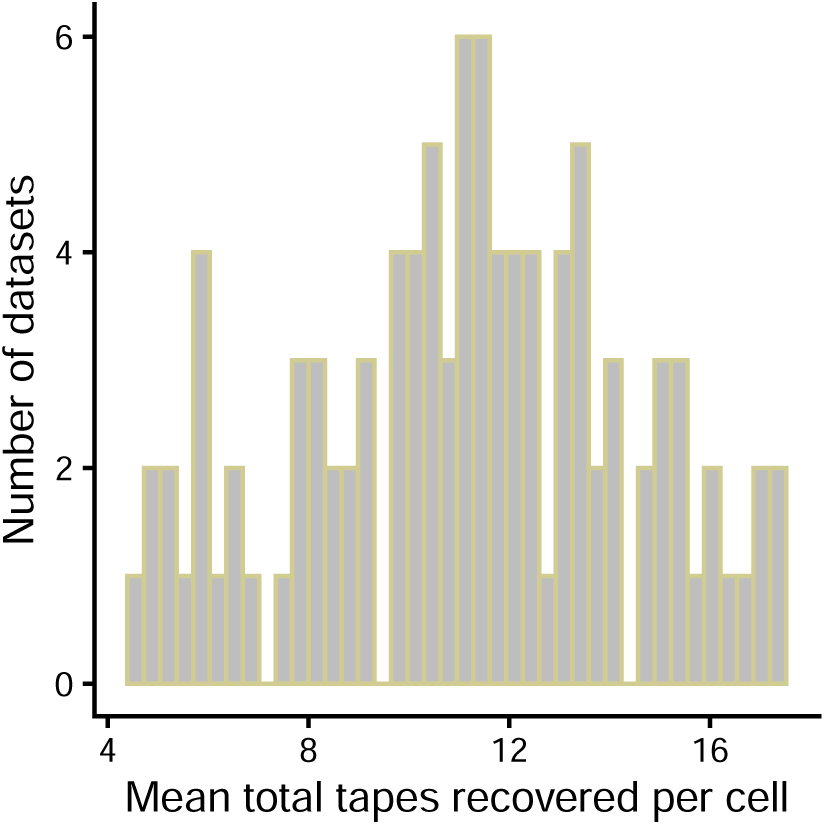
Simulations of incomplete tape alignments. Mean number of tapes recovered per cell per dataset for the simulations of incomplete alignments, out of the original 20 tapes.

#### A.2.2 Convergence metrics

**Table 8.**
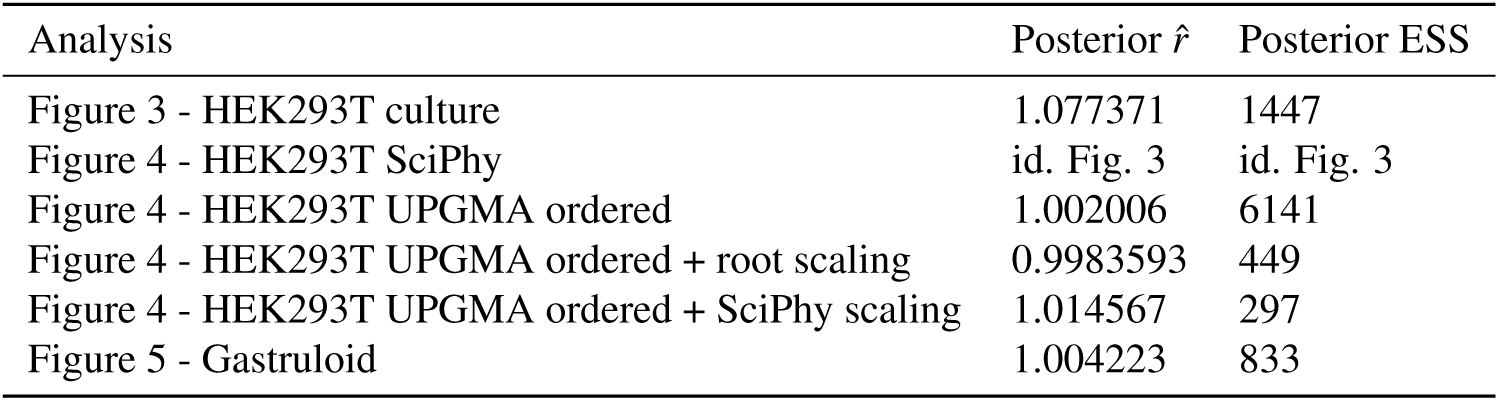
*r̂* and ESS convergence metrics for all analyses reported.

## REFERENCES

Alemany, A., Florescu, M., Baron, C. S., Peterson-Maduro, J., and Van Oudenaarden, A. (2018). Whole-organism clone tracing using single-cell sequencing. Nature, 556(7699):108–112.

Anzalone, A. V., Randolph, P. B., Davis, J. R., Sousa, A. A., Koblan, L. W., Levy, J. M., Chen, P. J., Wilson, C., Newby, G. A., Raguram, A., and Liu, D. R. (2019). Search-and-replace genome editing without double-strand breaks or donor DNA. Nature 2019 576:7785, 576(7785):149–157.

Beccari, L., Girgin, M., Turner, D. A., Baillie-Johnson, P., Cossy, A.-C., Beccari, L., Moris, N., Lutolf, M., Duboule, D., and Arias, A. M. (2018). Generating gastruloids from mouse embryonic stem cells. Protocol Exchange.

Berling, L., Klawitter, J., Bouckaert, R., Xie, D., Gavryushkin, A., and Drummond, A. J. (2025). Accurate bayesian phylogenetic point estimation using a tree distribution parameterized by clade probabilities. PLOS Computational Biology, 21(2):e1012789.

Biodata, S. (2024). Hek293t. https://www.cellosaurus.org/CVCL_0063.

Bouckaert, R., Heled, J., Kühnert, D., Vaughan, T., Wu, C.-H., Xie, D., Suchard, M. A., Rambaut, A., and Drummond, A. J. (2014). Beast 2: a software platform for bayesian evolutionary analysis. PLoS computational biology, 10(4):e1003537.

Bowling, S., Sritharan, D., Osorio, F. G., Nguyen, M., Cheung, P., Rodriguez-Fraticelli, A., Patel, S., Yuan, W.-C., Fujiwara, Y., Li, B. E., et al. (2020). An engineered crispr-cas9 mouse line for simultaneous readout of lineage histories and gene expression profiles in single cells. Cell, 181(6):1410–1422.

Chan, M. M., Smith, Z. D., Grosswendt, S., Kretzmer, H., Norman, T. M., Adamson, B., Jost, M., Quinn, J. J., Yang, D., Jones, M. G., et al. (2019). Molecular recording of mammalian embryogenesis. Nature, 570(7759):77–82.

Chen, K., Moravec, J. C., Gavryushkin, A., Welch, D., and Drummond, A. J. (2022). Accounting for errors in data improves timing in single-cell cancer evolution. bioRxiv, page 2021.03.17.435906.

Chen, W., Choi, J., Li, X., Nathans, J. F., Martin, B., Yang, W., Hamazaki, N., Qiu, C., Lalanne, J.-B., Regalado, S., et al. (2024). Symbolic recording of signalling and cis-regulatory element activity to dna. Nature, pages 1–9.

Choi, J., Chen, W., Minkina, A., Chardon, F. M., Suiter, C. C., Regalado, S. G., Domcke, S., Hamazaki, N., Lee, C., Martin, B., et al. (2022). A time-resolved, multi-symbol molecular recorder via sequential genome editing. Nature, 608(7921):98–107.

Chow, K.-H. K., Budde, M. W., Granados, A. A., Cabrera, M., Yoon, S., Cho, S., Huang, T.-H., Koulena, N., Frieda, K. L., Cai, L., et al. (2021). Imaging cell lineage with a synthetic digital recording system. Science, 372(6538):eabb3099.

Dawid, A. P. (1982). The well-calibrated bayesian. Journal of the American Statistical Association, 77(379):605–610.

de Jong, M. A., Adegeest, E., Bérenger-Currias, N. M., Mircea, M., Merks, R. M., and Semrau, S. (2024). The shapes of elongating gastruloids are consistent with convergent extension driven by a combination of active cell crawling and differential adhesion. PLOS Computational Biology, 20(2):e1011825.

Drummond, A. (2025). Treestat2 package. https://github.com/alexeid/TreeStat2.

Drummond, A. J., Ho, S. Y. W., Phillips, M. J., and Rambaut, A. (2006). Relaxed phylogenetics and dating with confidence. PLoS biology, 4(5):e88.

Drummond, A. J., Rambaut, A., Shapiro, B., and Pybus, O. G. (2005). Bayesian Coalescent Inference of Past Population Dynamics from Molecular Sequences. Molecular Biology and Evolution, 22(5):1185–1192.

Drummond, A. J. and Suchard, M. A. (2010). Bayesian random local clocks, or one rate to rule them all. BMC biology, 8(1):1–12.

Felsenstein, J. (1973). Maximum Likelihood and Minimum-Steps Methods for Estimating Evolutionary Trees from Data on Discrete Characters. Systematic Biology, 22(3):240–249.

Felsenstein, J. (2004). Inferring phylogenies. page 664.

Feng, J., Dewitt III, W. S., McKenna, A., Simon, N., Willis, A. D., and Matsen IV, F. A. (2021). Estimation of cell lineage trees by maximum-likelihood phylogenetics. The annals of applied statistics, 15(1):343.

Gong, W., Granados, A. A., Hu, J., Jones, M. G., Raz, O., Salvador-Martínez, I., Zhang, H., Chow, K. H. K., Kwak, I. Y., Retkute, R., Prusokas, A., Prusokas, A., Khodaverdian, A., Zhang, R., Rao, S., Wang, R., Rennert, P., Saipradeep, V. G., Sivadasan, N., Rao, A., Joseph, T., Srinivasan, R., Peng, J., Han, L., Shang, X., Garry, D. J., Yu, T., Chung, V., Mason, M., Liu, Z., Guan, Y., Yosef, N., Shendure, J., Telford, M. J., Shapiro, E., Elowitz, M. B., and Meyer, P. (2021). Benchmarked approaches for reconstruction of in vitro cell lineages and in silico models of c. elegans and m. musculus developmental trees. Cell Systems, 12:810–826.e4.

Gower, J. C. (1966). Some distance properties of latent root and vector methods used in multivariate analysis. Biometrika, 53(3-4):325–338.

He, Z., Maynard, A., Jain, A., Gerber, T., Petri, R., Lin, H.-C., Santel, M., Ly, K., Dupré, J.-S., Sidow, L., et al. (2022). Lineage recording in human cerebral organoids. Nature methods, 19(1):90–99.

Hoffman, M. D., Gelman, A., et al. (2014). The no-u-turn sampler: adaptively setting path lengths in hamiltonian monte carlo. J. Mach. Learn. Res., 15(1):1593–1623.

Kühnert, D., Stadler, T., Vaughan, T. G., and Drummond, A. J. (2016). Phylodynamics with Migration: A Computational Framework to Quantify Population Structure from Genomic Data. Molecular Biology and Evolution, 33(8):2102–2116.

Lewinsohn, M. A., Bedford, T., Müller, N. F., and Feder, A. F. (2022). State-dependent evolutionary models reveal modes of solid tumor growth. bioRxiv, page 2022.08.05.502978.

Loveless, T. B., Carlson, C. K., Hu, V. J., Dentzel Helmy, C. A., Liang, G., Ficht, M., Singhai, A., and Liu, C. C. (2021). Molecular recording of sequential cellular events into dna. bioRxiv, pages 2021–11.

Mai, U., Chu, G., and Raphael, B. J. (2024). Maximum likelihood inference of time-scaled cell lineage trees with mixed-type missing data. In International Conference on Research in Computational Molecular Biology, pages 360–363. Springer.

McKenna, A., Findlay, G. M., Gagnon, J. A., Horwitz, M. S., Schier, A. F., and Shendure, J. (2016). Whole-organism lineage tracing by combinatorial and cumulative genome editing. Science, 353(6298):aaf7907.

McKenna, A. and Gagnon, J. A. (2019). Recording development with single cell dynamic lineage tracing. Development (Cambridge), 146(12).

Mendes, F. K., Bouckaert, R., Carvalho, L. M., and Drummond, A. J. (2024). How to validate a bayesian evolutionary model. bioRxiv, page 2024.02.11.579856.

Merle, M., Friedman, L., Chureau, C., Shoushtarizadeh, A., and Gregor, T. (2024). Precise and scalable self-organization in mammalian pseudo-embryos. Nature Structural & Molecular Biology, pages 1–7.

Moris, N., Anlas, K., van den Brink, S. C., Alemany, A., Schröder, J., Ghimire, S., Balayo, T., van Oudenaarden, A., and Martinez Arias, A. (2020). An in vitro model of early anteroposterior organization during human development. Nature, 582(7812):410–415.

Mulberry, N. and Stadler, T. (2025). Strategies for resolving cellular phylogenies from sequential lineage tracing data. bioRxiv, pages 2025–01.

Pilarski, J., Stadler, T., and Seidel, S. (2025). Assessing the inference of single-cell phylogenies and population dynamics from genetic lineage tracing data. bioRxiv, pages 2025–01.

Regalado, S. G., Qiu, C., Kottapalli, S., Martin, B. K., Chen, W., Liao, H., Kim, H., Li, X., Lalanne, J.-B., Hamazaki, N., Domcke, S., Choi, J., and Shendure, J. (2025). Lineage recording in monoclonal gastruloids reveals heritable modes of early development. bioRxiv, page 2025.05.23.655664.

Seidel, S. and Stadler, T. (2022). Tidetree: a bayesian phylogenetic framework to estimate single-cell trees and population dynamic parameters from genetic lineage tracing data. Proceedings of the Royal Society B, 289(1986):20221844.

Sherri, M., Boulkaibet, I., Marwala, T., and Friswell, M. I. (2017). A differential evaluation markov chain monte carlo algorithm for bayesian model updating. arXiv preprint arXiv:1710.09486.

Smith, M. R. (2020). Information theoretic generalized robinson–foulds metrics for comparing phylogenetic trees. Bioinformatics, 36(20):5007–5013.

Smith, M. R. (2022). Robust analysis of phylogenetic tree space. Systematic Biology, 71(5):1255–1270.

Sokal, R. R., Sneath, P. H., et al. (1963). Principles of numerical taxonomy. Principles of numerical taxonomy.

Spanjaard, B., Hu, B., Mitic, N., Olivares-Chauvet, P., Janjuha, S., Ninov, N., and Junker, J. P. (2018). Simultaneous lineage tracing and cell-type identification using crispr–cas9-induced genetic scars. Nature biotechnology, 36(5):469–473.

Stadler, T. (2015). TreeSim: Simulating Phylogenetic Trees. R package version 2.2.

Stadler, T., Kühnert, D., Bonhoeffer, S., and Drummond, A. J. (2013). Birth–death skyline plot reveals temporal changes of epidemic spread in hiv and hepatitis c virus (hcv). Proceedings of the National Academy of Sciences, 110(1):228–233.

VanHorn, S. and Morris, S. A. (2021). Next-Generation Lineage Tracing and Fate Mapping to Interrogate Development. Developmental Cell, 56(1):7–21.

Varilly, P., Schifferli, M., Yang, K., Burcham, T., Cronan, P., Glennon, O., Jacks, O., Laning, E., Marrs, L., Oba, K., Yeung, S., Parker, E., Omah, I., Pekar, J. E., Luebbert, L., Andersen, K. G., Park, D. J., Schaffner, S. F., MacInnis, B. L., Happi, C., Lemieux, J. E., Ozonoff, A., Mitzenmacher, M. D., Fry, B., and Sabeti, P. C. (2025). Delphy: scalable, near-real-time bayesian phylogenetics for outbreaks. bioRxiv, page 2025.03.25.645253.

Vaughan, T. (2024). Feast package. https://github.com/tgvaughan/feast.git.

Vaughan, T. G., Kühnert, D., Popinga, A., Welch, D., and Drummond, A. J. (2014). Efficient Bayesian inference under the structured coalescent. Bioinformatics, 30(16):2272–2279.

Zwaans, A., Seidel, S., Manceau, M., and Stadler, T. (2025). A bayesian phylodynamic inference framework for single-cell crispr/cas9 lineage tracing barcode data with dependent target sites. Philosophical Transactions of the Royal Society B: Biological Sciences, 380.

