## Supplementary material for "SciPhy: A Bayesian phylogenetic framework using sequential genetic lineage tracing data": sciphy-supplement

### A APPENDIX

#### A.1 Validation

| Parameter | Symbol | Coverage [%] | Pearson's R [95%CI] |
|---|---|---|---|
| Editing rate | r | 91 | [0.995,0.998] |
| Insert probability 1 | $f_1$ | 96 | [0.978,0.991] |
| Insert probability 2 | $f_2$ | 95 | [0.983,0.992] |
| Insert probability 3 | $f_3$ | 93 | [0.974,0.988] |
| Insert probability 4 | $f_4$ | 92 | [0.978,0.990] |
| Insert probability 5 | $f_5$ | 98 | [0.979, 0.990] |
| Insert probability 6 | $f_6$ | 96 | [0.980, 0.991] |
| Insert probability 7 | $f_7$ | 96 | [0.969, 0.986] |
| Insert probability 8 | $f_8$ | 94 | [0.954, 0.979] |
| Insert probability 9 | $f_9$ | 91 | [0.947, 0.976] |
| Insert probability 10 | $f_{10}$ | 95 | [0.930, 0.968] |
| Insert probability 11 | $f_{11}$ | 99 | [0.988, 0.995] |
| Insert probability 12 | $f_{12}$ | 94 | [0.979, 0.990] |
| Insert probability 13 | $f_{13}$ | 94 | [0.981, 0.991] |
| Tree height | N/A | 98 | [0.855, 0.932] |
| Tree length | N/A | 96 | [0.9998, 0.9999] |
| Tree balance (B1 index) | N/A | 90 | [0.9995, 0.9998] |

**Table 6.** Coverages and correlations to true value in validation study

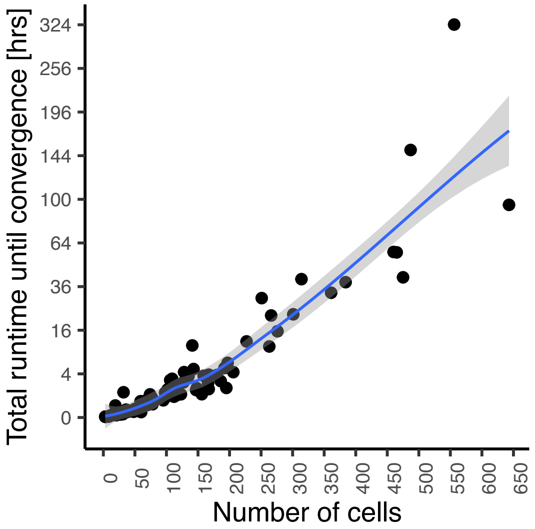


**Figure 1. Approximate runtime needed for convergence of a SciPhy analysis** We report the runtime required for convergence of the lineage tree reconstruction and parameter estimation for all datasets in the validation study. We display here the empirical total runtime rescaled to reach a minimum ESS value of 200 for the SciPhy likelihood.

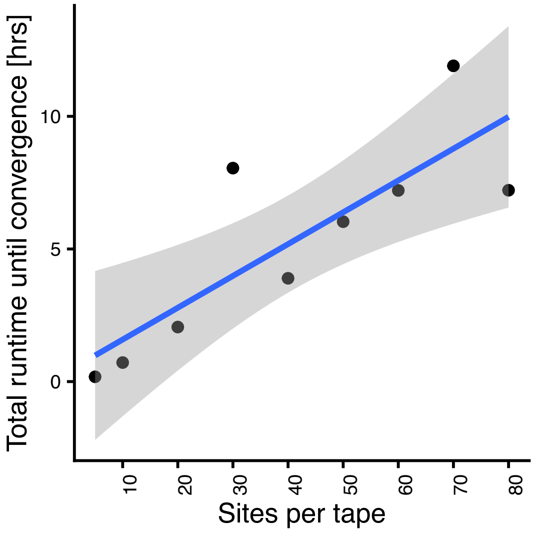


**Figure 2. Approximate runtime needed for convergence of a SciPhy analysis w.r.t. tape length.** We report the runtime required for convergence of the lineage tree reconstruction and parameter estimation based on 123 cells harboring a single tape, varying the number of sites per tape. We display here the empirical total runtime rescaled to reach a minimum ESS value of 200 for the SciPhy likelihood.

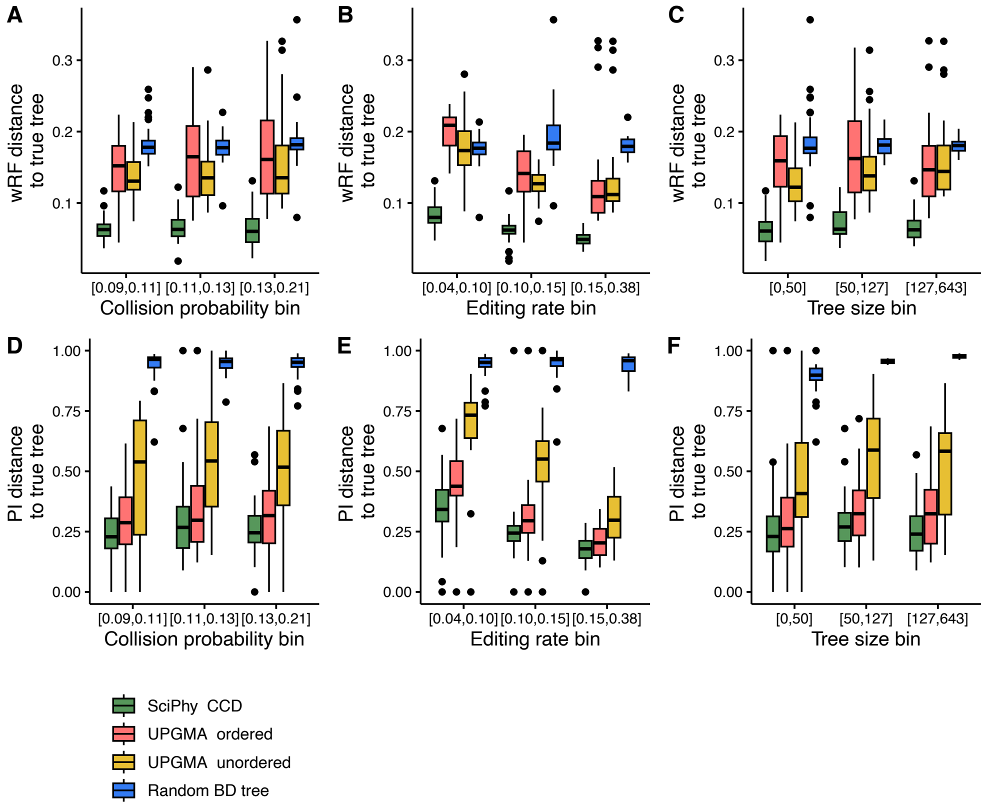


**Figure 3. Accuracy of SciPhy reconstructed trees w.r.t. different simulated parameter regimes, benchmarked to UPGMA on all validation datasets** For all datasets simulated in the validation study, we showcase the distance from the true tree to trees reconstructed with SciPhy (summarised using the Conditional Clade Distributions, denoted as SciPhy CCD), an order-aware UPGMA method (UPGMA ordered), the standard UPGMA method, ignoring the order between edits (UPGMA unordered), and randomly generated through simulation under the birth-death sampling model (Random BD tree), where UPGMA trees are all scaled to 25 days for comparison. These distances are shown for binned simulation parameters values and calculated using the weighted Robinson Foulds (wRF, panels A-C) metric and Phylogenetic Information metric (PI, panels D-F). The "collision probability" presented here is the sum of squared insert probabilities, and represents how skewed towards certain inserts the editing process is.

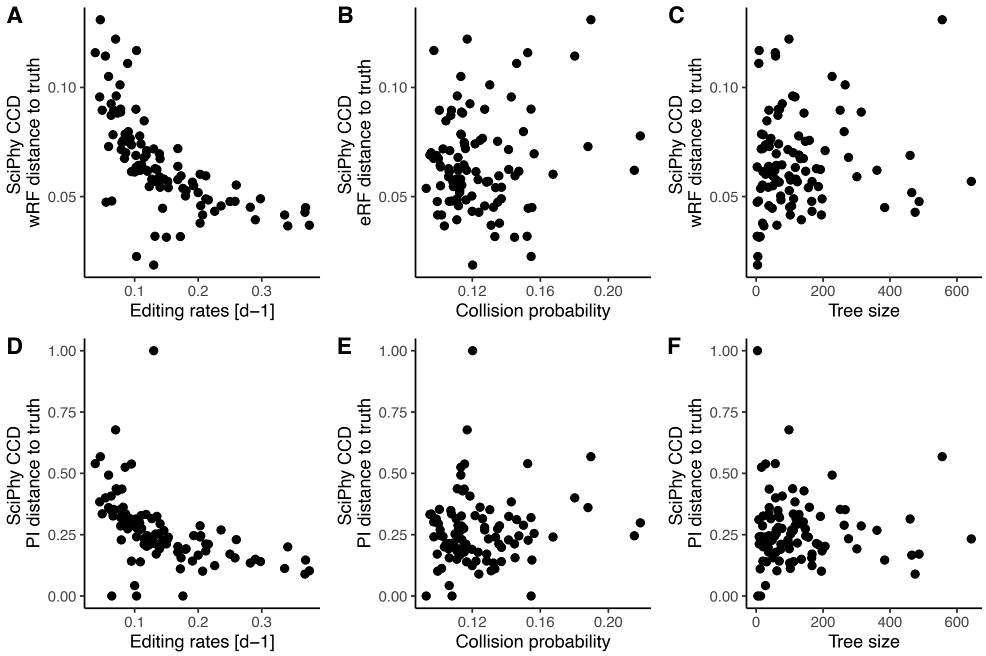


**Figure 4.** **Accuracy of SciPhy reconstructed trees w.r.t. simulated parameter values on all validation datasets** For all datasets simulated in the validation study, we showcase the distance from the true tree to trees reconstructed with SciPhy (summarised using the Conditional Clade Distributions, SciPhy CCD). These distances are shown against simulation parameters values and calculated using the weighted Robinson Foulds (wRF, panels A-C) metric and Phylogenetic Information metric (PI, panels D-F). The "collision probability" presented here is the sum of squared insert probabilities, and represents how skewed towards certain inserts the editing process is.

#### A.2 Cell culture

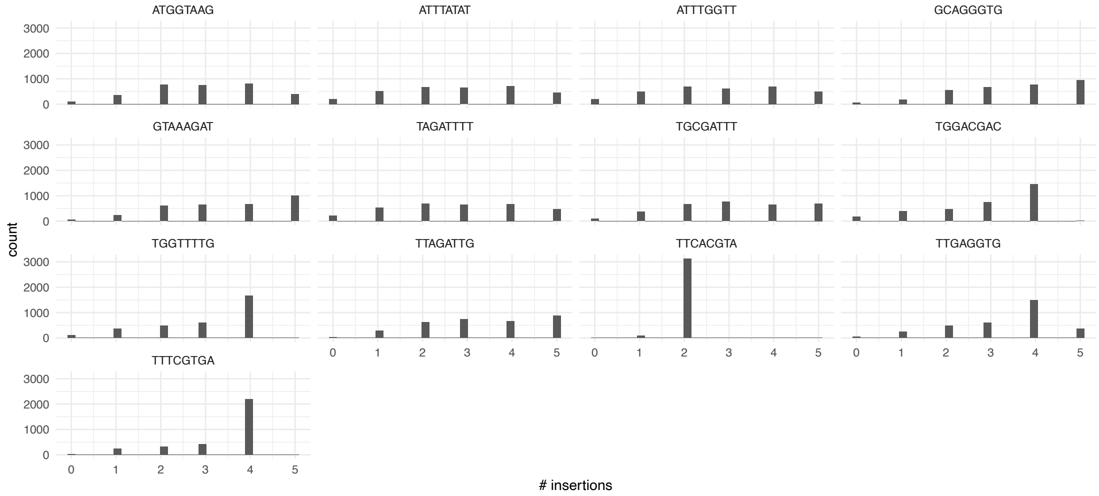


**Figure 5. Number of insertions per tape in the HEK293T dataset.** The majority of non truncated barcodes accumulate between 2 and 5 insertions throughout the experiment.

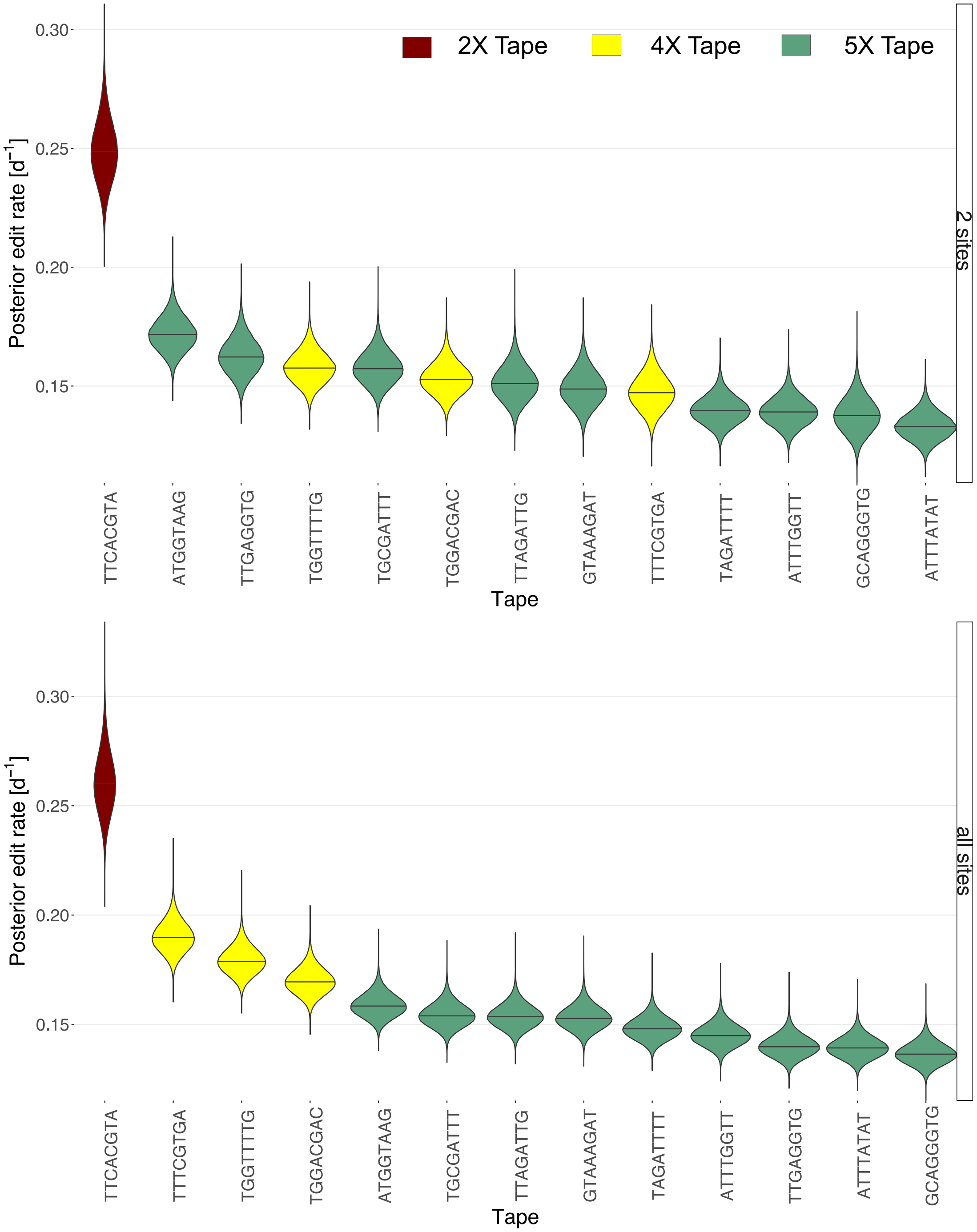


**Figure 6.** **Comparing estimated editing rates.** We show the estimated rate of editing for each tape. In the upper row, we show the results when estimating the editing rates from a dataset where all tapes are truncated to 2 sites. The lower row shows the analysis from the main text Fig. 3 where we include all available sites per tape. The edit rates are ordered by their median and are colored according to original tape length, where 2X means 2 sites etc.

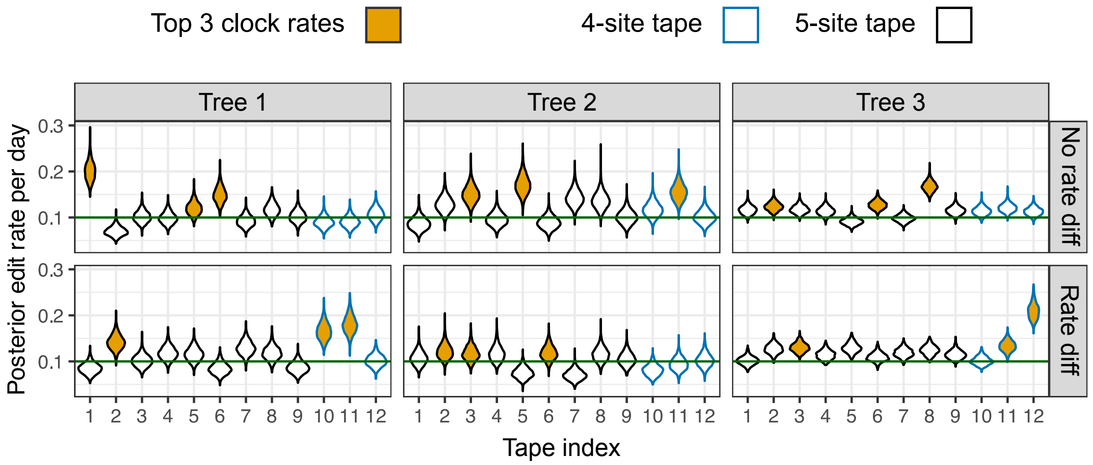


**Figure 7. Effect of within-tape rate variation on inferred editing rates.** Posterior estimates of edit rates per tape under SciPhy for three simulated trees. In each panel, tapes are shown along the x-axis (tape indices 1–12), with violin plots representing the posterior distribution of the per-tape edit rate. Tapes 10–12 (blue outlines) are 4-site tapes; others are 5-site tapes. Orange-filled violins indicate the three tapes with the highest median inferred edit rates. Top row: scenario with no rate differences between sites. Bottom row: scenario where the 5th site is edited at 20% of the base rate. In the "Rate diff" condition, 4-site tapes more frequently appear among the fastest edited rates, as they lack the slower 5th site.

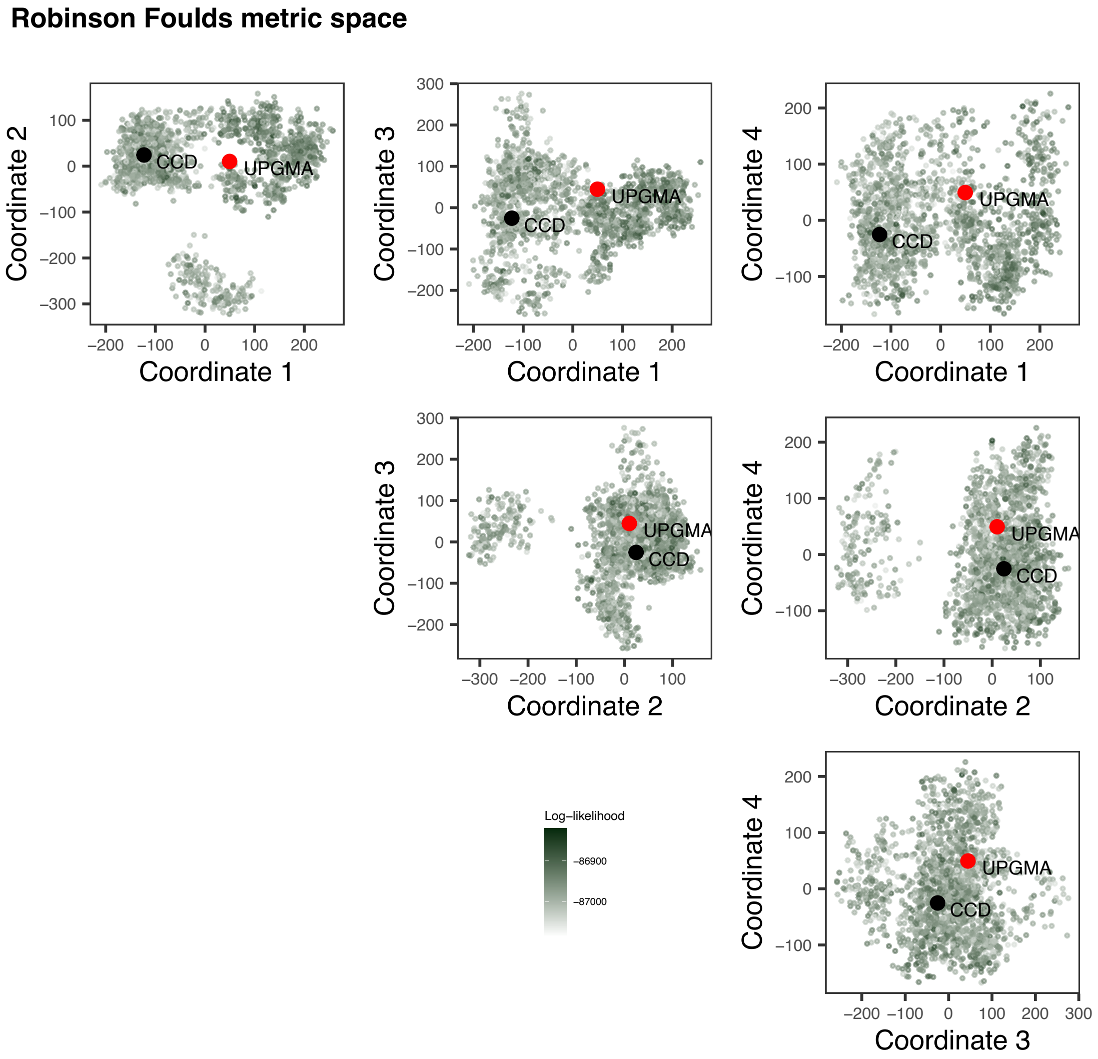


**Figure 8. Comparison of the SciPhy posterior tree set to the UPGMA tree obtained from the HEK293T data, with respect to topology.** Pairwise Robinson Fould (RF) distances between SciPhy posterior trees, the UPGMA tree and the CCD tree estimated for the HEK293T dataset are mapped in 4 coordinates pairwise to visualize these tree sets.

**Clustering Information metric space**

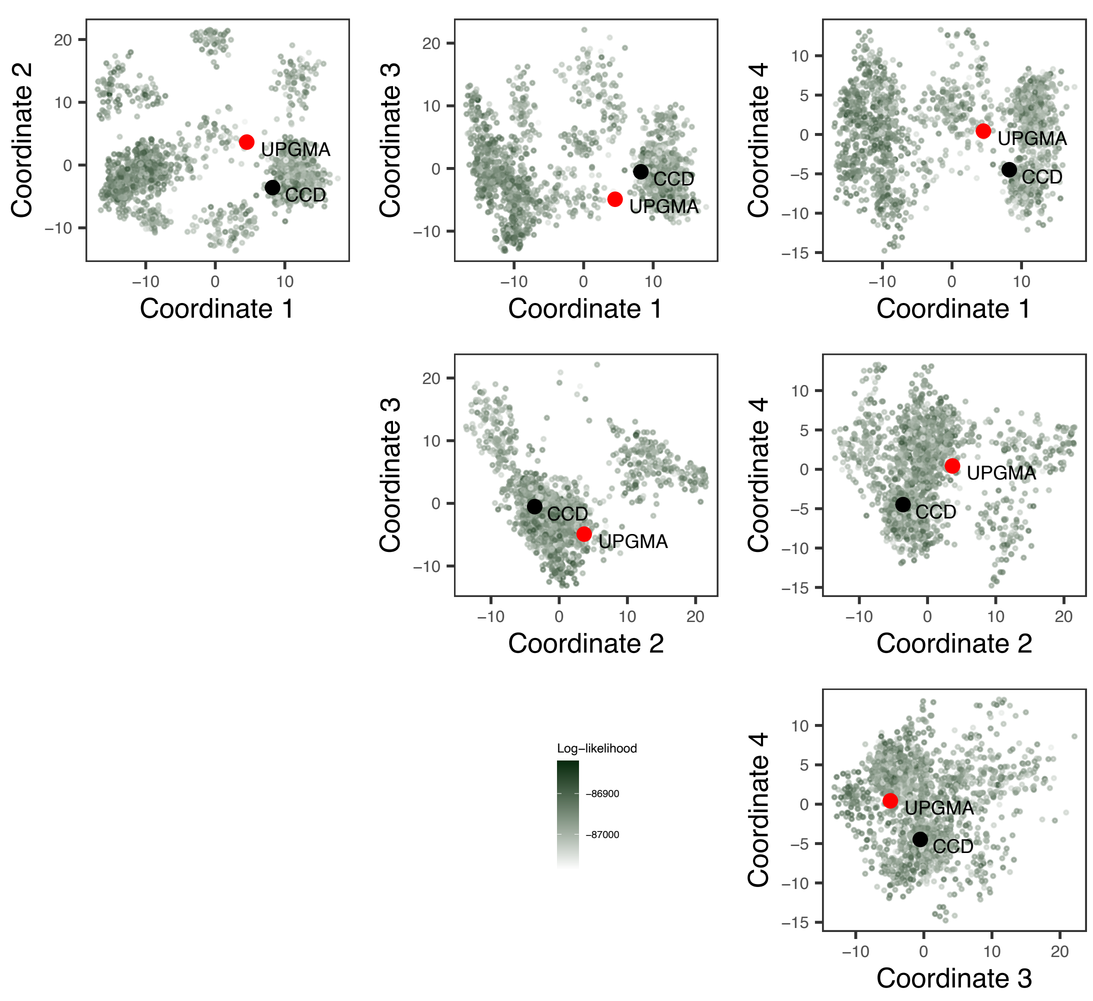


**Figure 9.** **Comparison of the SciPhy posterior tree set to the UPGMA tree obtained from the HEK293T data, with respect to topology.** Pairwise Clustering Information (CI) distances between SciPhy posterior trees, the UPGMA tree and the CCD tree estimated for the HEK293T dataset are mapped in 4 coordinates pairwise to visualize these tree sets.

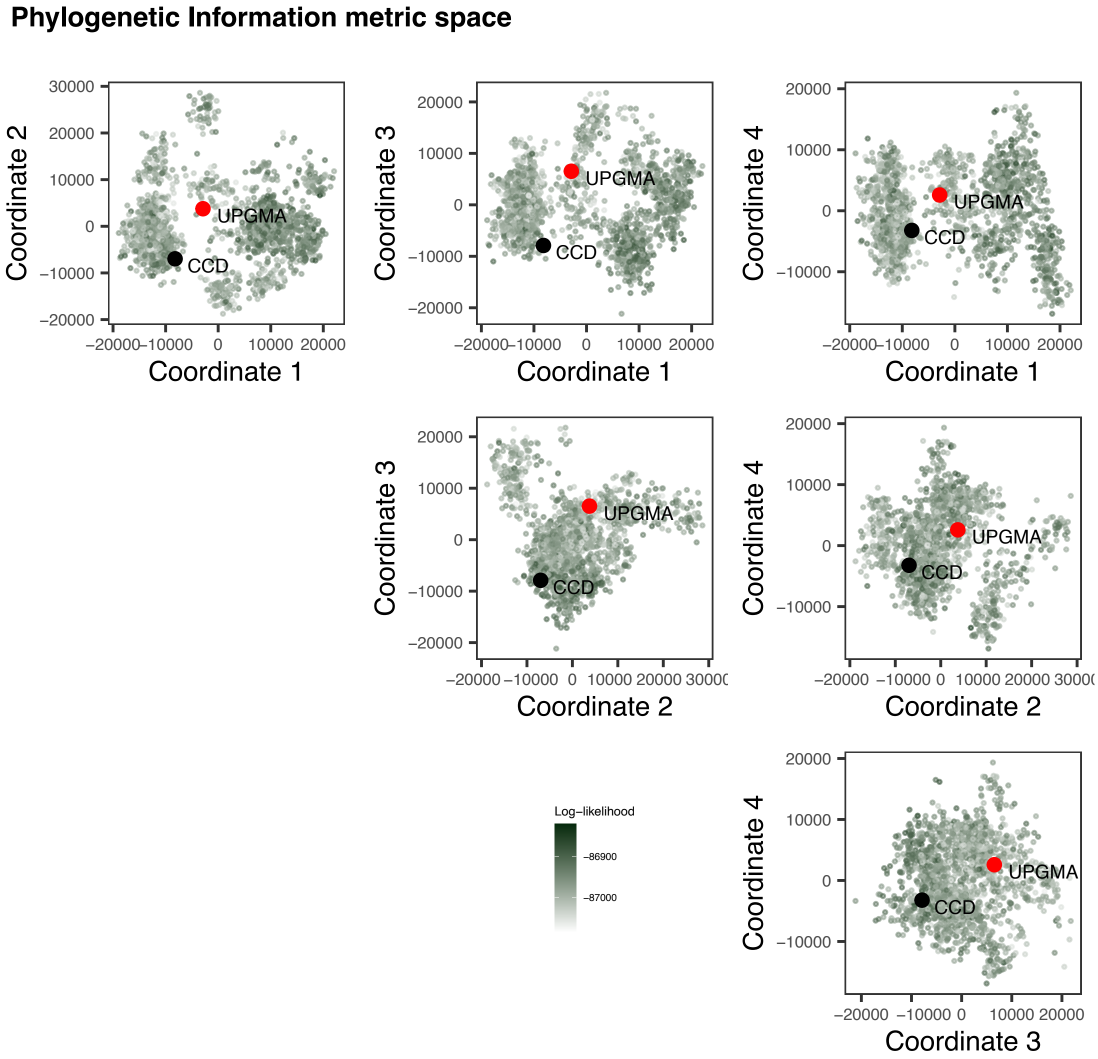


**Figure 10.** **Comparison of the SciPhy posterior tree set to the UPGMA tree obtained from the HEK293T data, with respect to topology.** Pairwise Phylogenetic Information (PI) distances between SciPhy posterior trees, the UPGMA tree and the CCD tree estimated for the HEK293T dataset are mapped in 4 coordinates pairwise to visualize these tree sets.

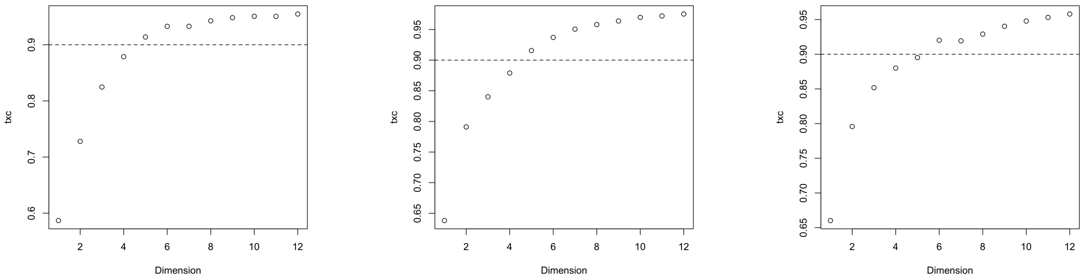


**(a)** Mapping quality for the RF distances against the number of dimensions represented.

**(b)** Mapping quality for the PI distances against the number of dimensions represented.

**(c)** Mapping quality for the CI distances against the number of dimensions represented.

**Figure 11. 2D mapping quality per number of dimensions plotted for the RF, PI and CI distances.** For each of the 2D mappings of pairwise topological distances presented in our study, we plot here the trustworthiness × continuity score (txc) against the number of dimensions from the mapping used the represent the sets of trees shown. A mapping is considered sufficiently representative for a txc score of about 0.9.

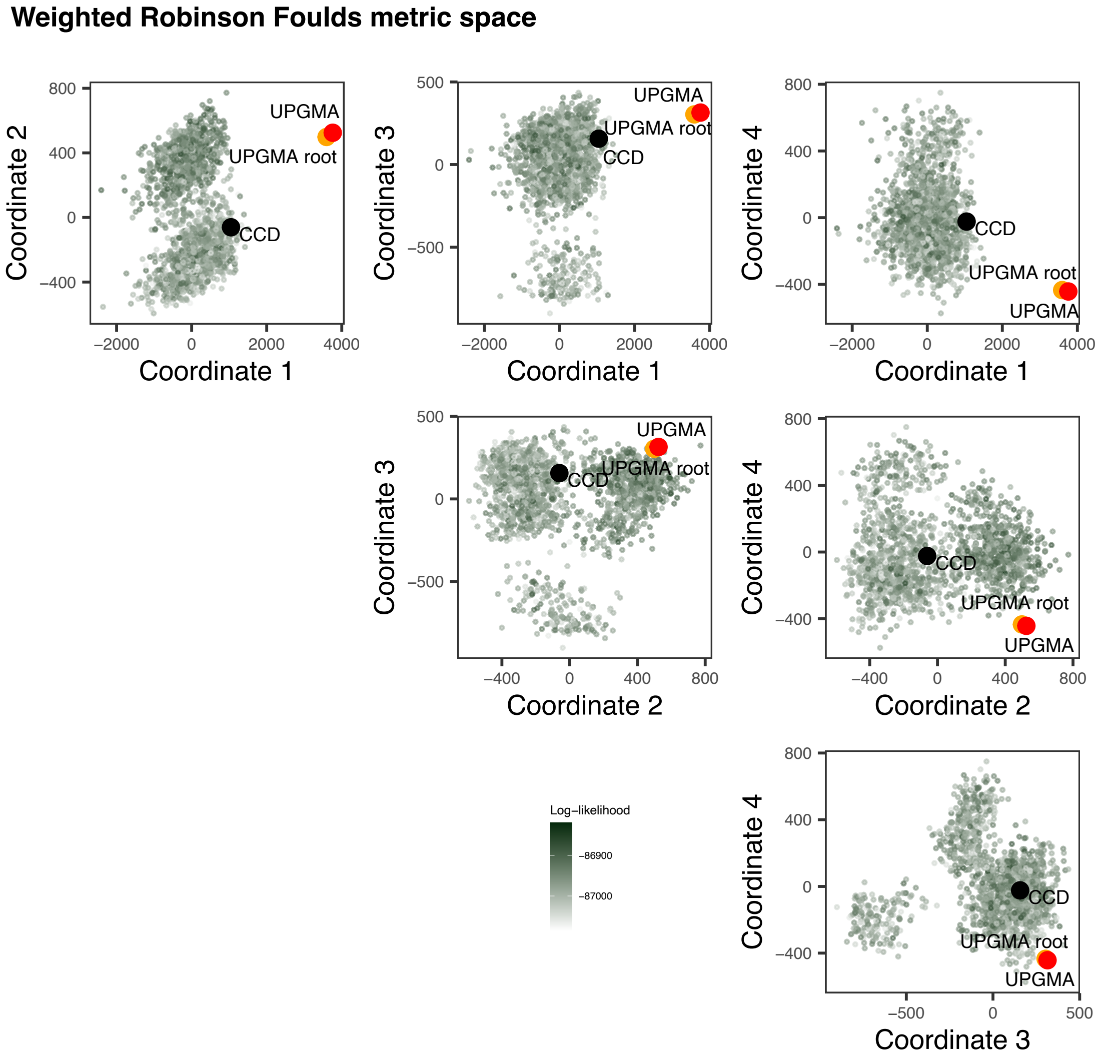


**Figure 12.** **Comparison of the SciPhy posterior tree set to the UPGMA tree obtained from the HEK293T data, with respect to topology and branch lengths.** Pairwise weighted Robinson Fould (wRF)distances between SciPhy posterior trees, the UPGMA tree (labelled 'UPGMA'), the UPGMA tree scaled to the median estimated tree height estimated by SciPhy (labelled 'UPGMA root') and the CCD tree estimated for the HEK293T dataset are mapped in 4 coordinates pairwise to visualize these tree sets.

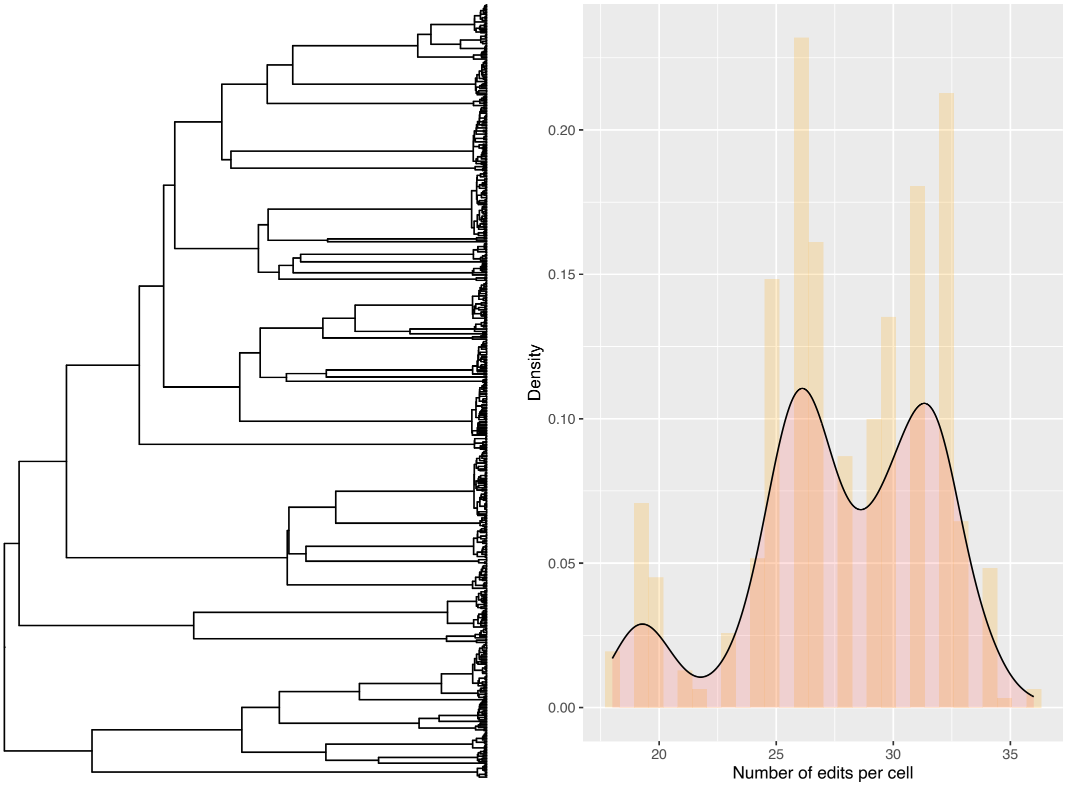


**Figure 13. Simulated distribution of the total number of edits (right) under SciPhy along a tree (left) with early divisions.** Here, SciPhy data (13 tapes of length 5) was simulated along a tree where the majority of branching events happen toward the present. This corresponds to a scenario where the numbers of edits per cell are not independent for the majority of the history of the cell population, and where the structure of the lineage tree leaves an imprint on the resulting distribution of the total edits per cell at the tips.

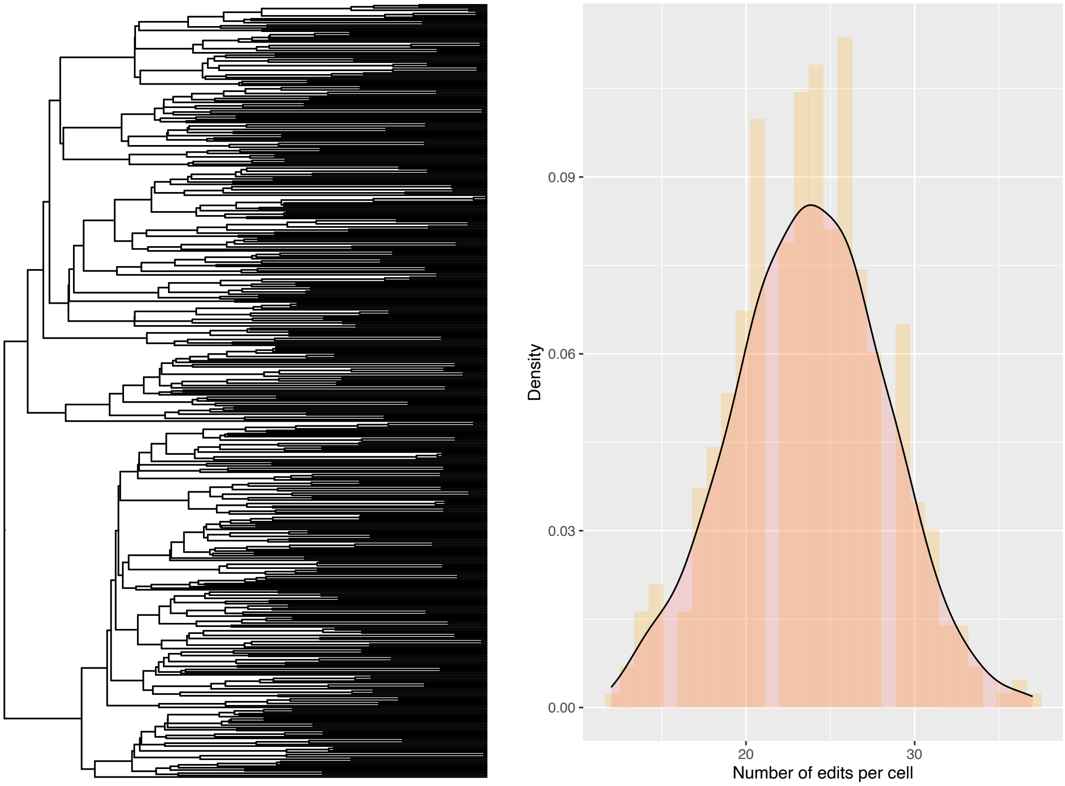


**Figure 14.** **Simulated distribution of the total number of edits (right) under SciPhy along a tree (left) with late divisions** Here, SciPhy data (13 tapes of length 5) was simulated along a tree where the majority of branching events happen toward the past. This corresponds to a scenario where the numbers of edits per cell are independent for the majority of the history of the cell population, resulting in a unimodal distribution of the total number of edits.

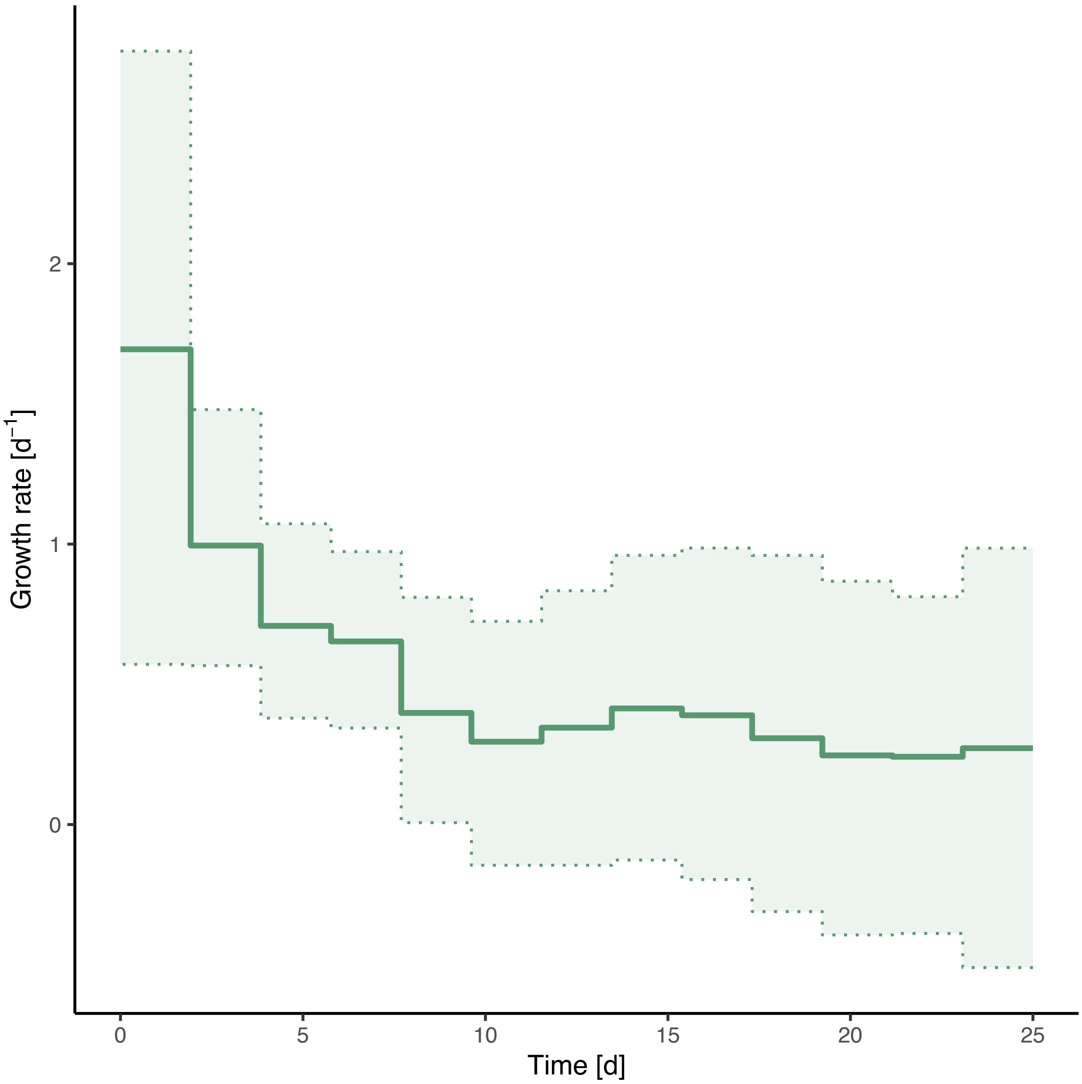


**Figure 15. Time-varying (dynamic) growth rate estimated for the HEK293T cell population, with a sampling proportion fixed to 0.0008** We report the time-varying growth rate estimated using SciPhy, where the growth rate is allowed to vary every 2 days over the duration of the experiment, using an Ornstein-Ulhenbeck (OU) smoothing prior.

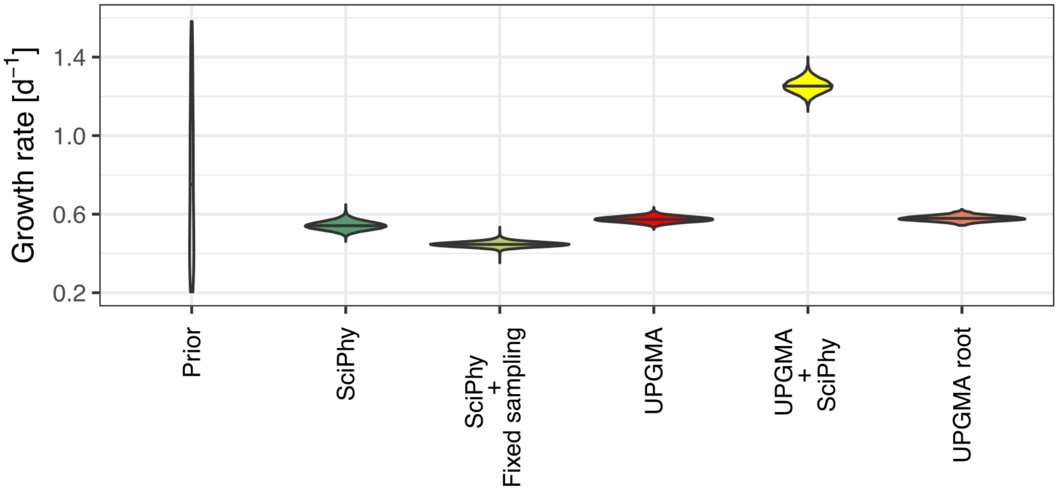


**Figure 16.** **Comparing growth rates estimated with SciPhy and on fixed phylogenies.** We report the constant growth rate estimated using SciPhy, with the sampling proportion fixed to 0.0008 ("SciPhy + Fixed sampling"), with a co-estimated sampling proportion ("SciPhy"), contrasted to inferences based on a fixed UPGMA tree topology. For comparison, we look at the UPGMA tree topology with the root height scaled to 25 days ("UPGMA"), with its root scaled to the median SciPhy tree MRCAs and the UPGMA tree with branch lengths scaled using SciPhy ("UPGMA - SciPhy").

##### *A.2.1 Sampling from prior*

To investigate whether the growth rate is informed mainly by the molecular sequence data (the tapes in our study), or the number of sequenced cells at the end of the experiment and the prior distributions, we re-ran the inference devoid of the sequence data (Appendix Fig. 17). This approach yielded a median growth rate of 0.57 per day (95% HPD interval: [0.48, 0.68] $d^{-1}$), notably different from the estimate with sequence data and aligning with the expected growth rate under a deterministic exponential growth model. This suggests that the slower cell division rate we initially estimated is largely influenced by the sequence data.

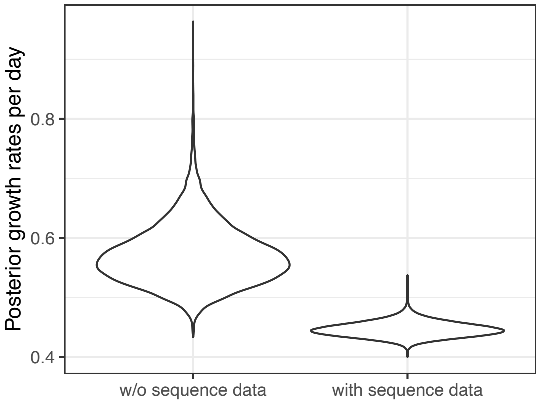


**Figure 17. Comparing growth rate estimates with and without sequence data in the HEK293T analysis.** We report the growth rates estimated using SciPhy as in the main text (with sequence data, right) and compare it to the same analysis just without the sequence data (left).

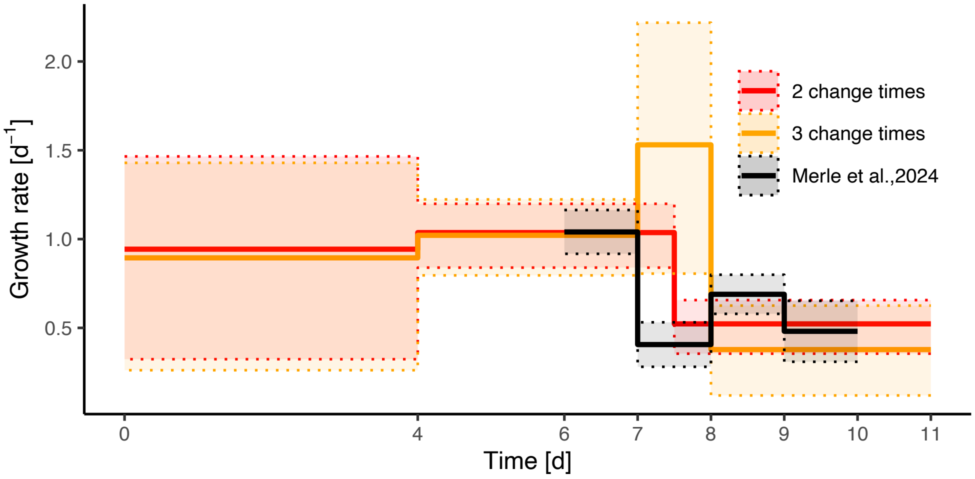


**Figure 18.** **Comparing estimates of the time-varying growth rate for gastruloid data under different time divisions of the experimental timeline and against observed dynamics.** We report the growth rates (median and 95% HPD) estimated using SciPhy under 2 change times (also in main text), and (3 change times) and the growth rates (mean $\pm\sigma$) reported in Merle et al. (2024) for comparison.

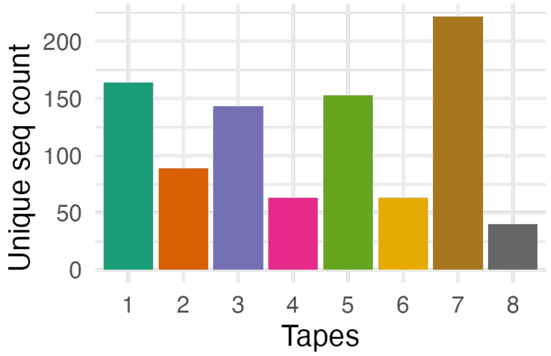


**Figure 19.** We show the number of unique insert sequences for each tape.

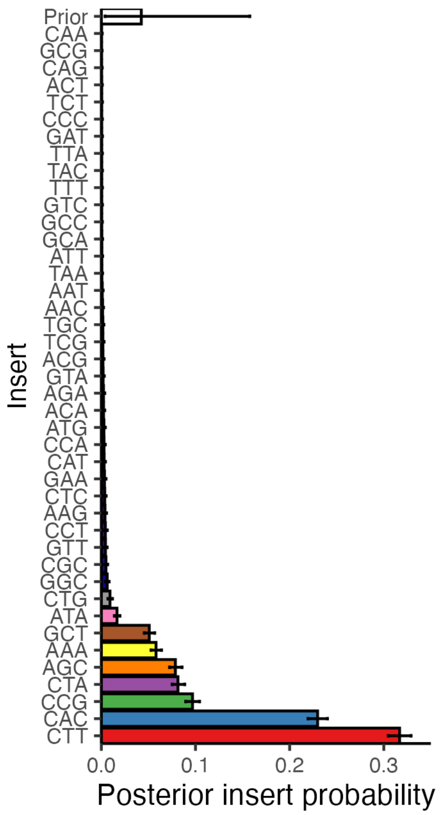


**Figure 20.** We show the posterior insertion probability for every possible insert. Note that we use a highly distinguishable color scheme for the first 9 inserts and a less distinguishable color scheme for the remaining 33 inserts, that have very low insertion probability.

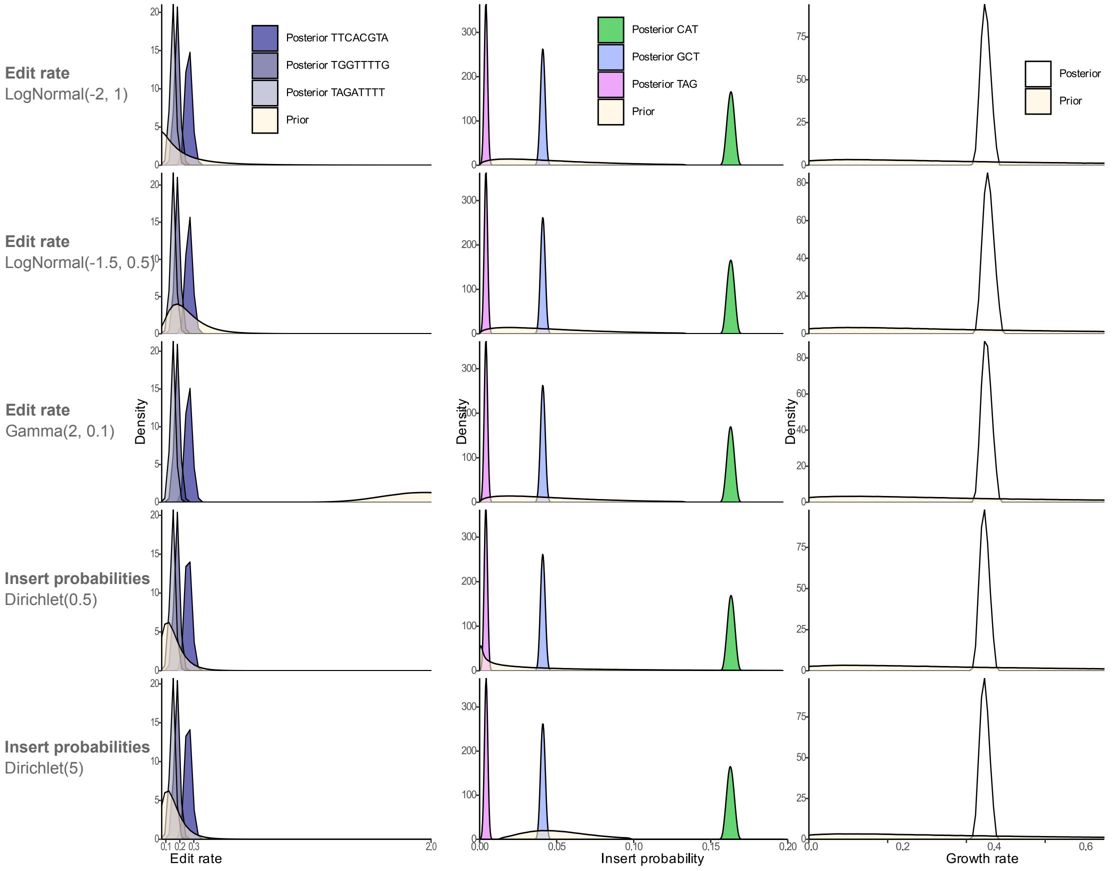


**Figure 21.** Results of the phylodynamic analysis of cell culture growth are robust to different choices of prior distributions.

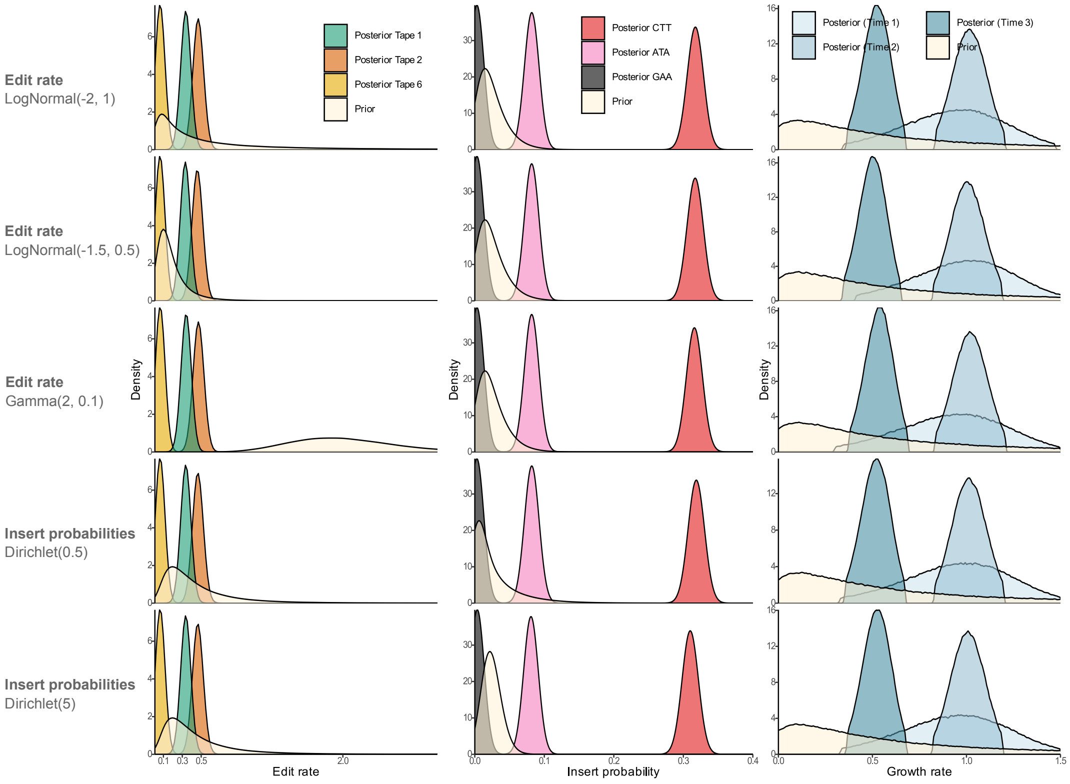


**Figure 22.** Results of the phylodynamic analysis of gastruloid growth are robust to different choices of prior distributions.

##### *A.2.2 Simulation of incomplete tape alignments*

| Parameter name | Symbol | Distribution | HDI |
|---|---|---|---|
| Clock rate | $r$ | Log-normal ($\mu = -2, \sigma = 0.5$) | [0.03, 0.3] |
| Edit probabilities | $f_k$ | Dirichlet ($\alpha = 1.5$) | [2e-4, 2e-1] |
| Heritable barcode loss rate | N/A | Log-normal ($\mu = -5, \sigma = 1.0$) | [0.00095, 0.048] |
| Dropout probability | N/A | Beta ($\alpha = 3, \beta = 7$) | [0.075, 0.6] |

**Table 7.** Distributions for the editing model parameters used to simulate incomplete barcode alignments.

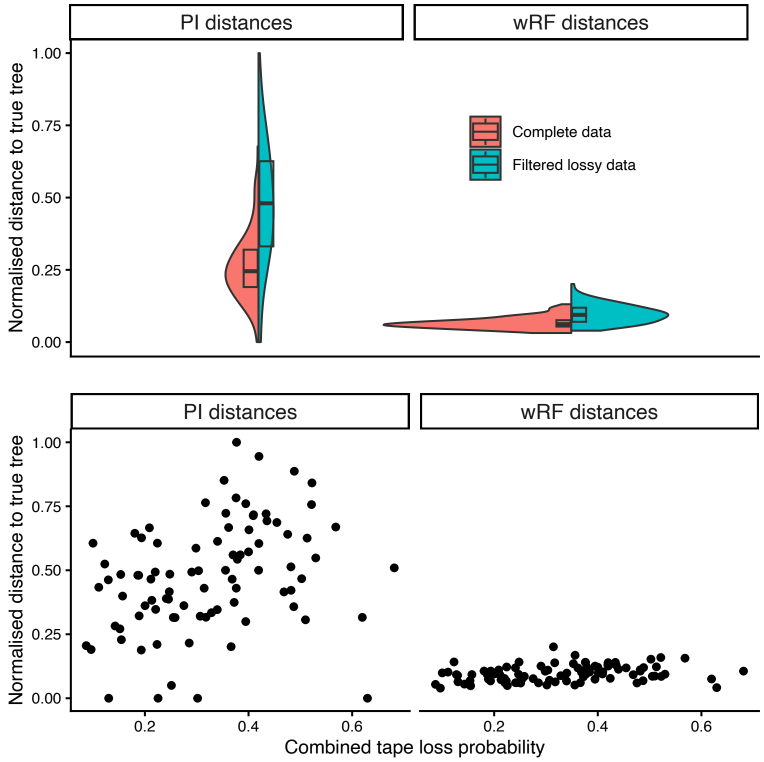


**Figure 23. Robustness of tree inference using SciPhy to sparsity in tape alignments.** In the top panel, for all trees simulated in the validation study, we showcase the distance from the true tree to trees reconstructed with SciPhy (summarized as point estimates with the conditional clade distribution (CCD) algorithm ), using complete alignments of 10 tapes ("Complete data") or lossy alignments ("Filtered lossy data") of 20 tapes (leading to an average of 11 tapes after filtering) as input. In the bottom panel, we additionally show these distances against the combined tape loss (resulting of both heritable probability used for simulation of lossy alignments.

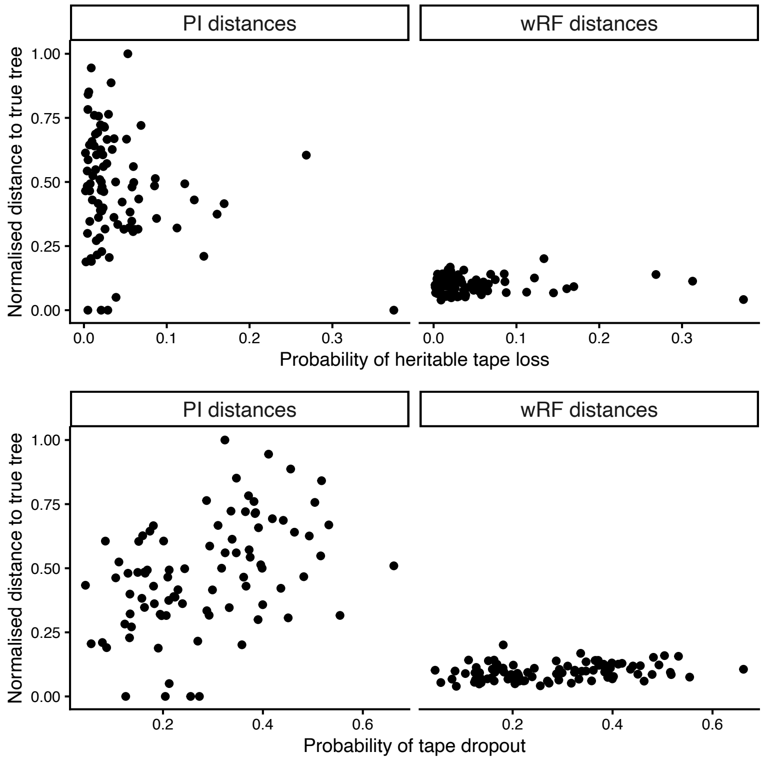


**Figure 24.** **Robustness of tree inference using SciPhy to sparsity in tape alignments.** For all trees simulated in the validation study, we showcase the distance from the true tree to trees reconstructed with SciPhy (summarized as point estimates with the CCD algorithm) using lossy alignments ("Filtered lossy data") of 20 tapes (leading to an average of 11 tapes after filtering) as input. In the top panel, we show these distances against the heritable tape loss probability (or transgene silencing probability) used for simulation, while in the bottom panel we show these distances against the tape dropout probabilities (or probability of loss upon sampling).

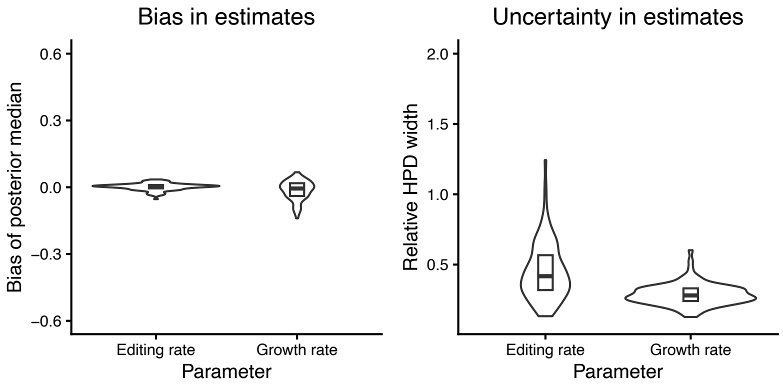


**Figure 25.** **Robustness of parameter inference using SciPhy to sparsity in tape alignments.** For all datasets simulated in the validation study, we showcase the bias (left) and uncertainty (right) in editing and growth estimates obtained using incomplete alignments ("Filtered lossy data" in Figure 23) of 20 tapes as input.

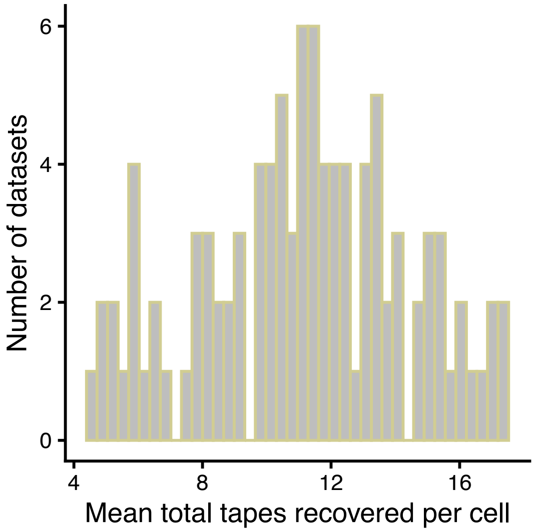


**Figure 26.** **Simulations of incomplete tape alignments.** Mean number of tapes recovered per cell per dataset for the simulations of incomplete alignments, out of the original 20 tapes.

###### A.2.3 Convergence metrics

| Analysis | Posterior $\hat{r}$ | Posterior ESS |
|---|---|---|
| Figure 3 - HEK293T culture | 1.077371 | 1447 |
| Figure 4 - HEK293T SciPhy | id. Fig. 3 | id. Fig. 3 |
| Figure 4 - HEK293T UPGMA ordered | 1.002006 | 6141 |
| Figure 4 - HEK293T UPGMA ordered + root scaling | 0.9983593 | 449 |
| Figure 4 - HEK293T UPGMA ordered + SciPhy scaling | 1.014567 | 297 |
| Figure 5 - Gastruloid | 1.004223 | 833 |

**Table 8.** $\hat{r}$ and ESS convergence metrics for all analyses reported
